# Design of four component T=4 tetrahedral, octahedral, and icosahedral protein nanocages through programmed symmetry breaking

**DOI:** 10.1101/2023.06.16.545341

**Authors:** Sangmin Lee, Ryan D. Kibler, Yang Hsia, Andrew J. Borst, Annika Philomin, Madison A. Kennedy, Barry Stoddard, David Baker

## Abstract

Four, eight or twenty C3 symmetric protein trimers can be arranged with tetrahedral (T-sym), octahedral (O-sym) or icosahedral (I-sym) point group symmetry to generate closed cage-like structures^1,2^. Generating more complex closed structures requires breaking perfect point group symmetry. Viruses do this in the icosahedral case using quasi-symmetry or pseudo-symmetry to access higher triangulation number architectures^3–9^, but nature appears not to have explored higher triangulation number tetrahedral or octahedral symmetries. Here, we describe a general design strategy for building T = 4 architectures starting from simpler T = 1 structures through pseudo-symmetrization of trimeric building blocks. Electron microscopy confirms the structures of T = 4 cages with 48 (T-sym), 96 (O-sym), and 240 (I-sym) subunits, each with four distinct chains and six different protein-protein interfaces, and diameters of 33nm, 43nm, and 75nm, respectively. Higher triangulation number viruses possess very sophisticated functionalities; our general route to higher triangulation number nanocages should similarly enable a next generation of multiple antigen displaying vaccine candidates^10,11^ and targeted delivery vehicles^12,13^.

## Main text

Natural and designed protein nanocages consist of one or multiple unique components arranged with tetrahedral, octahedral, and icosahedral point group symmetry^1,3–7,14–16^. While there are no other point group symmetries, in the icosahedral case viruses access larger and more complex higher triangulation (T) number architectures by interspersing varying numbers of hexagons between 12 pentagonal substructures (pentons), which expand the structure without changing the curvature. Viruses achieve this symmetry breaking by placing at symmetrically non-equivalent positions either the same subunit, but in different conformations (quasi-symmetry)^3–8^, or closely related but distinct subunits (pseudo-symmetry)^9^. The accessing of higher T number icosahedral structures by such symmetry breaking is critical to the remarkable functionality of viruses, such as the ability to package and deliver large nucleic acid cargos. Similarly, the de novo design of higher T number protein assemblies could enable new approaches to nucleic acid delivery and, as the potency of nanoparticle immunogens can increase with increasing valency of display, could lead to more potent vaccines. However, while protein design has had considerable success in designing symmetric assemblies which assemble from identical interacting subunits^1,14–16^, the design of assemblies with multiple identical or nearly identical chains in non-symmetry equivalent positions is an outstanding challenge.

We set out to develop a systematic approach to design higher T number protein assemblies that could generate not only icosahedral (I-sym) architectures but also tetrahedral (T-sym) and octahedral (O-sym) higher T number nanostructures which to our knowledge have not been found in Nature. Of the two routes to breaking symmetry described above, we reasoned that using closely related but distinct subunits (pseudo-symmetry) would have the advantage over quasi-symmetry as it avoids the complexity of designing single subunits with multiple distinct states and sets of interactions. However, pseudo-symmetry requires a set of building blocks that are structurally nearly identical but have distinct interaction surfaces; for example, expanding T = 1 nanocages built from homo-trimers placed on the polyhedral 3-fold symmetry axes to generate a specific higher T number architecture requires heterotrimers with three distinct chains but overall shapes nearly identical to the homotrimer. While there has been progress in designing 3 chain protein heterotrimers^17^, these do not have near 3-fold symmetry nor a homo-trimeric analogue with a near identical structure.

We reasoned that the design of higher T number tetrahedral, octahedral and icosahedral nanostructures could be achieved using ABC-type pseudo-symmetric heterotrimers to precisely program the six distinct interfaces in such structures, provided the complexity of the design challenge could be broken down into a series of individually experimentally validatable steps. We adopted the three step hierarchical approach outlined in Fig. 1. We first design T = 1 cages by arranging C3 symmetric homotrimers along the 3-fold symmetry axis of each architecture (Fig. 1 left two columns)^1,2^. Following experimental validation, we next extract cyclic “crown” like substructures from these cages by replacing the symmetric homotrimers with ABC-type heterotrimers that lack the cage-forming trimer-trimer interface on one of the subunits, and redesigning the crown-forming trimer-trimer interface to be heterodimeric (Fig. 1 third and fourth columns). Following experimental validation, these crown-like structures are arranged along one of the 3-fold axes of T-sym, the 4-fold axis of O-sym, or the 5-fold symmetry axis of I-sym architectures, and an additional symmetric homotrimer is placed on the remaining 3-fold axis (Fig. 1 fifth column). This generates four component T = 4 cages for each cage symmetry, with a hexameric motif between edges of the crowns (*h* = 2 and *k* = 0 in the Caspar and Klug nomenclature^8^).

**Fig. 1.**
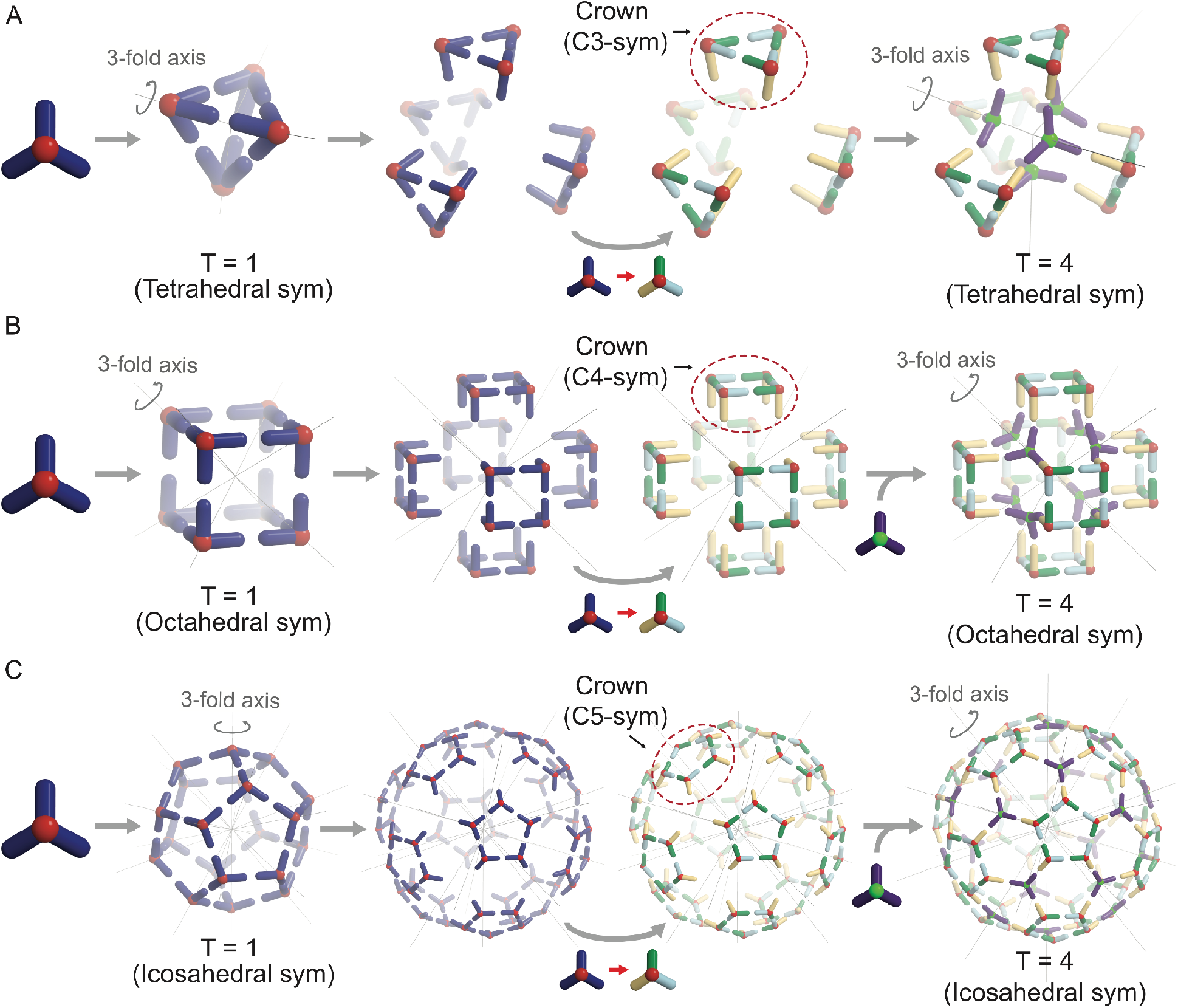
Overview of design strategy. (**A-C**) Schematic illustration of route to T=4 (A) tetrahedral, (B) octahedral, and (C) icosahedral cages. Step 1 (first and second columns): C3 cyclic homo-oligomers are docked and designed into T = 1 cages. Step 2 (third and fourth columns): Shifting parts constituting each face of the T = 1 cages away from the origin along the symmetry axis orthogonal to the face and replacing the homotrimers to ABC-type pseudo-symmetric heterotrimers produce crowns in which one of the three components (yellow) is free to be designed to dock to other building blocks. Step 3 (fifth column): The crowns are docked by homotrimers aligned along the 3-fold symmetry axis of each symmetry, which produces T = 4 cages.

### Design of pseudo-symmetric building blocks

Our design strategy requires homotrimeric building blocks that can be replaced by heterotrimeric blocks that make identical interactions with their neighbors (i.e., with distinct internal interfaces within the heterotrimer, but identical outward facing interfaces), and that can be docked into nanocages with different architectures. As described in the accompanying manuscript^18^, as a first step towards the creation of pseudo-symmetric nanocages, we computationally designed a family of “BGL” ring shaped homo- and heterotrimers which have the same structure but different amino acid sequences at the protomer-protomer interfaces. Using a combination of native mass spectrometry (nMS) and X-ray crystallography, we found that the heterotrimer designs exclusively populate states containing all three chains in equal stoichiometries, and that both homotrimer and heterotrimer designs assemble into symmetric cyclic rings very close to the computational design models^18^.

To enable docking of BGL rings into closed architectures through helix-helix interfaces (Fig. 2A), we rigidly fused onto the rings a variety of helical repeat protein (DHR) “arms”^19^through HelixFuse^20^, yielding “armed” homotrimers ∼10 nm in diameter (Fig. 2B - 2E). From the combinations of the 13 BGLs and 38 DHRs, we generated more than 2000 different armed BGLs and re-designed residues around the junction using Rosetta^21^ (Supplementary Materials Section 1). A total of 39 designs, numbered BGL09_A01 through BGL19_A39 where BGLXX indicates the base BGL design^18^ and AYY identifies the arm attached to the BGL (Table S3) were selected for experimental characterization. AlphaFold (AF2)^22^ structure predictions for 32/39 designs were close to the design models (RMSD < 3.0 Å) (Fig. S1). 37/39 of the designs were soluble following expression in *E. coli*. and purification from immobilized metal affinity chromatography (IMAC), 33/39 had size-exclusion chromatography (SEC) retention volumes consistent with the design models and for 22 the SEC profiles were monodisperse (Fig. S2-S3). The homotrimeric state was confirmed by nMS for 27/39 (Fig. S4-S6), and small-angle X-ray scattering (SAXS) profiles were closely consistent with profiles computed from the design models for 19 (Fig. S7-S8). Negative-stain electron microscopy (nsEM) characterization of the overall shapes of the designs was again consistent with the design models, with curved, straight, wide, and narrow arm arrangements evident in 2D class averages and 3D reconstructions (Fig. 2B, 2D and Fig. S9). We obtained a crystal structure of one of the designs, BGL17_A31, which was close to the design model and even closer to the AF2 prediction (Fig. 2C).

**Fig. 2.**
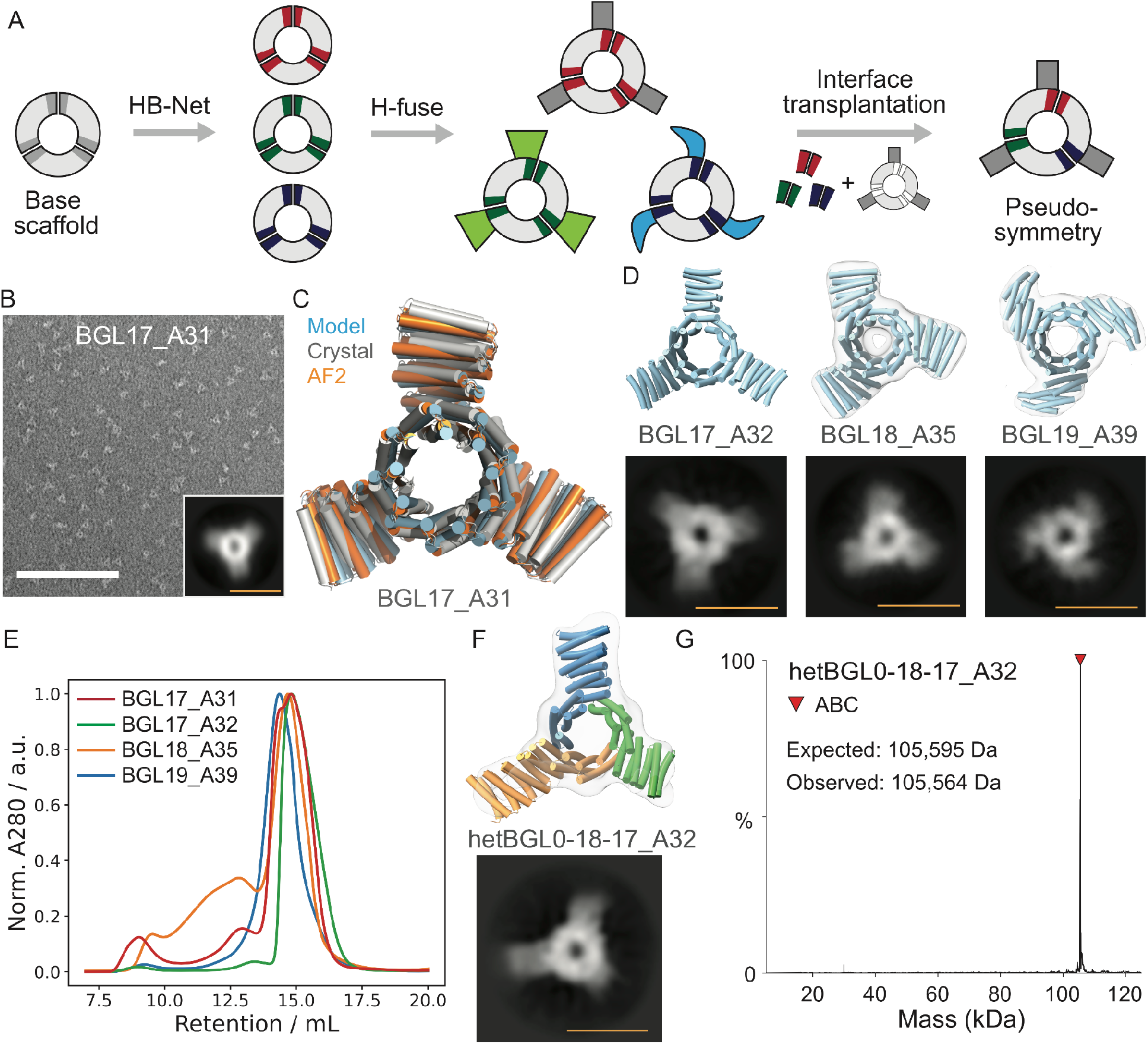
Design of pseudo-symmetric ABC heterotrimers with extensible arms. (**A**) Schematic illustration of design strategy for homotrimers with arms and pseudo-symmetric heterotrimers. (**B**) nsEM micrograph of BGL17_A31 homotrimer and a 2D class average (inset) along 3-fold symmetry axis. (**C**) Structure of BGL17_A31 solved by X-ray crystallography (white) compared to its design model (light blue) and AF2 prediction (orange). (**D**) Top row: superpositions of the 3D reconstructed nsEM map (transparent cloud) to the structure model (blue) of BGL17_A32 (no map) (left), BGL18_A35 (middle) and BGL19_A39 (right). Bottom row: a 2D class average along 3-fold symmetry axis. (**E**) SEC results of homotrimers. (**F**) ABC-type pseudo-symmetric heterotrimer (hetBGL0-18-17_A32) with the same DHR arms of different lengths extending each component. Each color of the model represents each component. Top: superposition of the 3D reconstructed nsEM map (transparent cloud) and model (colors). Bottom: a 2D class average along (pseudo-) 3-fold symmetry axis. (**G**) nMS result of hetBGL0-18-17_A32. Scale bars: (white) 100 nm and (yellow) 10 nm.

To design pseudo-symmetric heterotrimers, we used an interface transplantation approach (Fig. S10-S11), preserving the overall structural C3 symmetry (Supplementary Materials, Section 2). A homotrimer was selected as the host scaffold and three different protomer-protomer interfaces from different homo-oligomeric BGLs (guests) were transplanted onto the subunit-subunit interfaces (Fig. 2A), conserving the residues at the arm junction. We first checked the compatibility between a host and a guest in the homo-oligomer context by symmetrically transplanting the guest interface into the host. We selected seven homotrimers (BGL0_A10, BGL0_A11, BGL17_A31, BGL17_A32, BGL18_A35, BGL19_A38, BGL19_A39) as host scaffolds and four BGLs (BGL0, BGL17, BGL18, BGL19)^18^ as guest interfaces because both groups were expressed at high levels and formed homogenous homotrimers. We experimentally characterized 18 of the 28 combinations (Table S4); 14/18 had SEC peaks at the correct oligomeric size (Fig. S12) and 11/18 had strong nMS signals at the correct masses (Fig. S13). The BGL17_A32 host backbone was found to be compatible with multiple guest interfaces; and we constructed heterotrimers by splicing interfaces from different guests together in different combinations^18^. To enable assignment of chain type by EM, we varied the number of repeat units on the arms protruding from each chain (−1 repeat for the A component and +1 to C component). Three of five heterotrimers (hetBGL0-17-19_A32, hetBGL0-18-17_A32, hetBGL0-19-17_A32) formed ABC-type heterotrimers as shown by SEC, SDS-PAGE, and nMS (Fig. 2F and Fig. S14-S17), and for two of these (hetBGL0-18-17_A32 and hetBGL0-19-17_A32), the expected differences in arm lengths were clearly evident in 2D nsEM averages and 3D reconstructions (Fig. 2F and Fig. S18; in hetBGLXX-XX-XX_AYY, XXs indicate the three BGL interfaces^18^ of the heterotrimer and YY indicates the arm (Table S5)).

### Cage design

We generated base T = 1 tetrahedral, octahedral, and icosahedral cages from the BGL17_A32 homotrimer using RPXdock^23,24^ to sample rotational and translational displacements of the trimer C3 axis along the 3-fold cage axes (Supplementary Materials, Section 3). We found that for the different symmetries, different numbers of DHR-arm repeat units (1.5, 2.5, and 3.5 repeat units for tetrahedral, octahedral, and icosahedral cages respectively) were optimal for docking of the trimers with good shape complementarity between the four helices interacting across the interface because of the twist of the DHR-arm (Fig. 3A, 3B, 3E, 3F, 3I, 3J). The newly generated cage interfaces were designed using ProteinMPNN^25^, and designs for which the AF2 prediction of the arm-arm interface was less than 2.0 Å RMSD from the design model were selected for experimental characterization – these comprise seven tetrahedral cages (Tet_T=1_-1 - Tet_T=1_-7) with 12 subunits and diameters of ∼13nm, eight octahedral cages (Oct_T=1_-1 - Oct_T=1_-8) with 24 subunits and diameters of ∼20nm, and four icosahedral cages (Ico_T=1_-1 - Ico_T=1_-4 with 60 subunits and diameters of ∼40nm. All of the tetrahedral designs had peaks at the expected retention volume (Fig. S20) on SEC, and four (Tet_T=1_-1, Tet_T=1_-2, Tet_T=1_-4, Tet_T=1_-6) were structurally homogeneous by nsEM with 2D class averages and 3D reconstructed nsEM maps (Fig. 3C and S21) matching the design models (Fig. 3B and 3C). Seven of the octahedral cages had single peaks on SEC, and two (Oct_T=1_-2 and Oct_T=1_-4) showed homogenous structures matching the design models by nsEM (Fig. 3F, 3G, S22-S23). One of the icosahedral cages, Ico_T=1_-1, had a single peak on SEC and was close to the design model by nsEM (Fig. 3J and 3K) although imperfectly formed cages were also observed (Fig. S24).

**Fig. 3.**
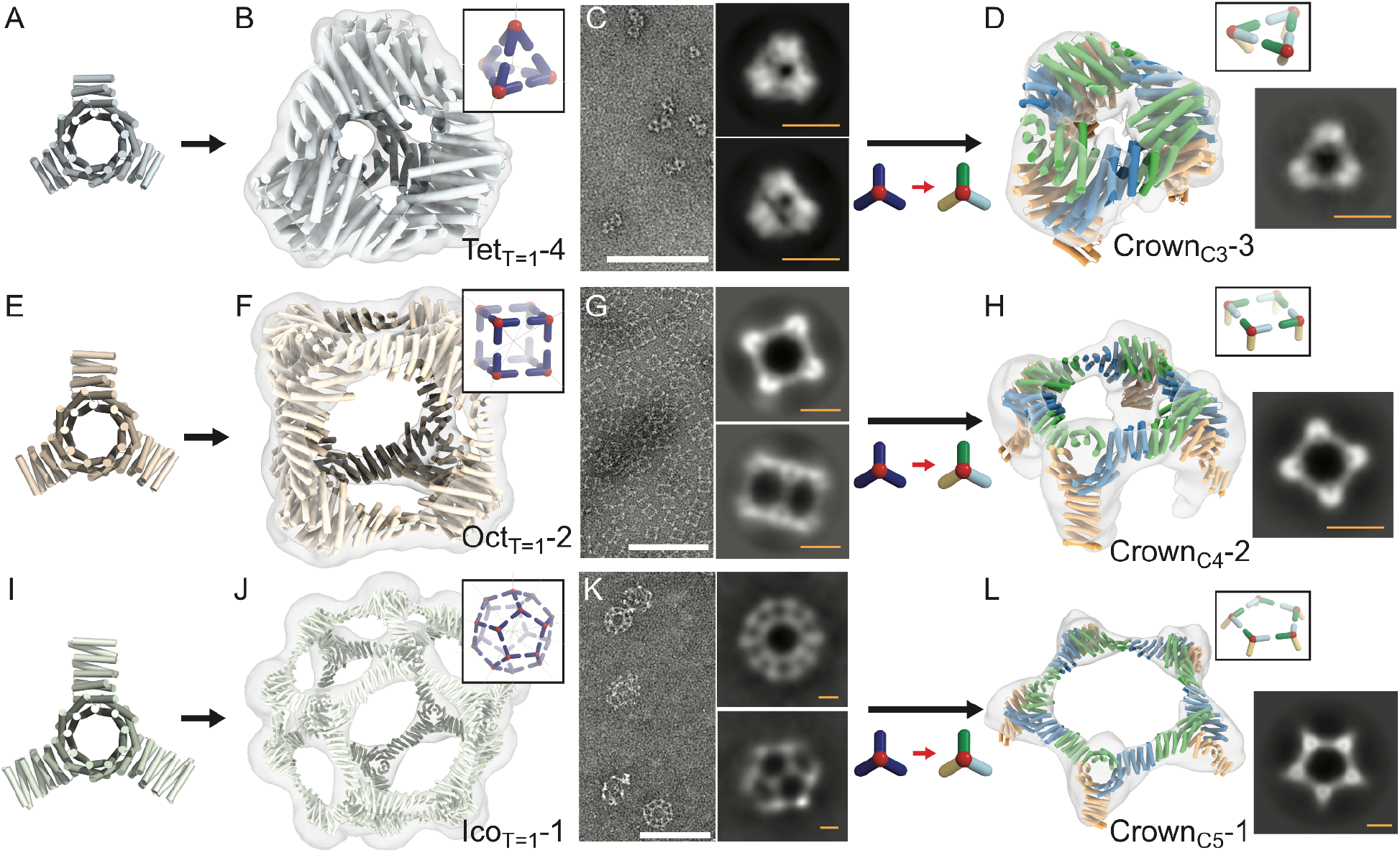
Extraction of homotrimer cycles (crowns) from T=1 cages by pseudo-symmetrization. BGL17_A32 with (**A**) 1.5, (**E**) 2.5 and (**I**) 3.5 repeat units of arms docked into (**B**,**C**) tetrahedral (Tet_T=1_-4), (**F**,**G**) octahedral (Oct_T=1_-2) and (**J**,**K**) icosahedral (Ico_T=1_-1) T=1 cages. (**B**,**F**,**J**) Superpositions of 3D reconstructed nsEM map (transparent cloud) to the design model of cages (colors). (**C**,**G**,**K**) (left) nsEM micrographs and (right) characteristic 2D class averages of the cages. (**D**,**H**,**L**) C3 (Crown_C3_-3), C4 (Crown_C4_-2) and C5 (Crown_C5_-1) crowns made by pseudo-symmetric heterotrimers. (left) Superpositions of 3D reconstructed nsEM map and the model crowns. Each color of the model represents each component (green: ch_A, blue: ch_B, orange: ch_C). (right) 2D class averages along 3-fold, 4-fold and 5-fold symmetry axes. The diameter of crowns is 11nm (C3), 20nm (C4) and 35nm (C5). Scale bars: (white) 100 nm and (yellow) 10 nm.

We next extracted C3, C4, and C5 symmetric cyclic oligomers (which we refer to as crowns because of their shape) from the structurally confirmed T = 1 cages by substituting in the structurally identical pseudo-symmetric hetBGL0-18-17_A32 heterotrimer in place of the BGL17_A32 homotrimer (Fig. 1 third and fourth columns), and designing the chain A (ch_A) and ch_B interfaces to interact at the crown trimer-trimer interface using ProteinMPNN – this was necessary to avoid a potential off target structure possible with the original C2 interface (Fig. S26). The surface of the arm of ch_C, which points outwards from the crown, was redesigned to be entirely polar. We selected designs for which AF2 predicted the ch_A - ch_B interface with RMSD < 2Å and did not predict the ch_A - ch_A or ch_B - ch_B homodimer interfaces to form (Fig. S19). We obtained genes encoding 19 sets of crowns that passed these filters (Crown_C3_-1 - Crown_C3_-5 for C3 crowns, Crown_C4_-1 - Crown_C4_-7 for C4 crowns, Crown_C5_-1 - Crown_C5_-7 for C5 crowns), and the three chains for each crown were expressed separately in independent *E. coli* cultures. Following expression, the amount of each protein was estimated by SDS-PAGE gel densitometry, and appropriate amounts of culture were combined to achieve mixtures with stoichiometric amounts of the three chains which were co-lysed and co-purified. By SEC, 4/5 C3, 5/7 C4, and 2/7 C5 crowns had peaks at the expected elution volumes (Fig. S26-S31) containing three distinct bands by SDS-PAGE (Fig. S32), indicating that the complexes were heterotrimers. nsEM 2D class averages and 3D reconstructed nsEM maps matched well with the crown design models (Fig. 3D, 3H, 3L). Thus symmetric substructures can be extracted from larger symmetric assemblies by substituting homotrimers with pseudosymmetric heterotrimers.

In the final step of our hierarchical design approach, we designed T = 4 cages by combining the experimentally confirmed crowns with the BGL17_A32 homotrimer (Fig. 1 last column) (Supplementary Materials, Section 3). The C3, C4, and C5 crowns were aligned with the 3-fold, 4-fold, and 5-fold axes of tetrahedral, octahedral, and icosahedral architectures, and BGL17_A32 was aligned to the remaining 3-fold axis. This generates assemblies in which the heterotrimer arms pointing outward from the crowns (ch_C of heterotrimer) interact with the arms of the homotrimer (ch_ho). To find optimal docking interfaces, we used RPXdock, sampling the lengths of the interacting arms and the rotations and translations along the common axis, and designed sequences using proteinMPNN for the highest RPX scoring models for each symmetry. Designs were filtered based on formation of the designed interface, and lack of formation of the self interfaces, in AF2 predictions (Fig. S19). Very few initial designs for the octahedral architecture avoided the formation of self interfaces according to AF2 predictions, so to decrease the probability of self interaction we performed explicit negative design using proteinMPNN^25^ against the predicted self interfaces. We experimentally tested 14 sets of T = 4 cage designs (Tet_T=4_-1 - Tet_T=4_-5, Oct_T=4_-1 - Oct_T=4_-4, and Ico_T=4_-1 - Ico_T=4_-5) that passed the AF2 filters. The four components were expressed independently in different *E. coli*. cultures, mixed with 1:1:1:1 stoichiometry, and co-lysed. The lysed samples were purified using IMAC and SEC, and the cage structures were characterized by nsEM (Fig. S35-S47). As described in the following paragraphs, the major species in each case was the designed T=4 structure; we also observed minor species of smaller off-target T=1-like cages (Fig. S37, S42, S45).

The T = 4 tetrahedral cage (Tet_T=4_-2) has a tetrapod shape (diameter: 33 nm) with the four C3 crowns pointing outward, and the homotrimers bridging the crowns closer to the center of the cage and facing inward (Fig. 4A, 4D, 4G, and S36). The homotrimer-heterotrimer distance (11.5 nm) is almost twice as long as the heterotrimer-heterotrimer distance (6.0 nm) due to the DHR-arm length difference between components (1.5 repeat units for ch_A and ch_B, 3 repeat units for ch_C and ch_ho), and the interior volume is a tetrahedral channel 6.0 nm in width (Fig. 4D). Overall, the structure maps to a T = 4 Goldberg polyhedra with tetrahedral symmetry^26^, in which the hexagonal motifs between triangles are highly elongated (Fig. 4A). These structural features are evident in the nsEM map (Fig. 4D), micrographs (Fig. 4E), and (2D average classes (Fig. 4F), and the design model is closely consistent with the reconstructed nsEM map (Fig. 4G). The design model could be readily relaxed to fit the nsEM 3D map with all four components clearly within density (Fig S36); overall the relaxed model matches well with the design model, with the exception of a slight twist of the overall structure resulting from curvature in the DHR-arm near the homotrimer-heterotrimer interface (Fig. 4D and S36).

**Fig. 4.**
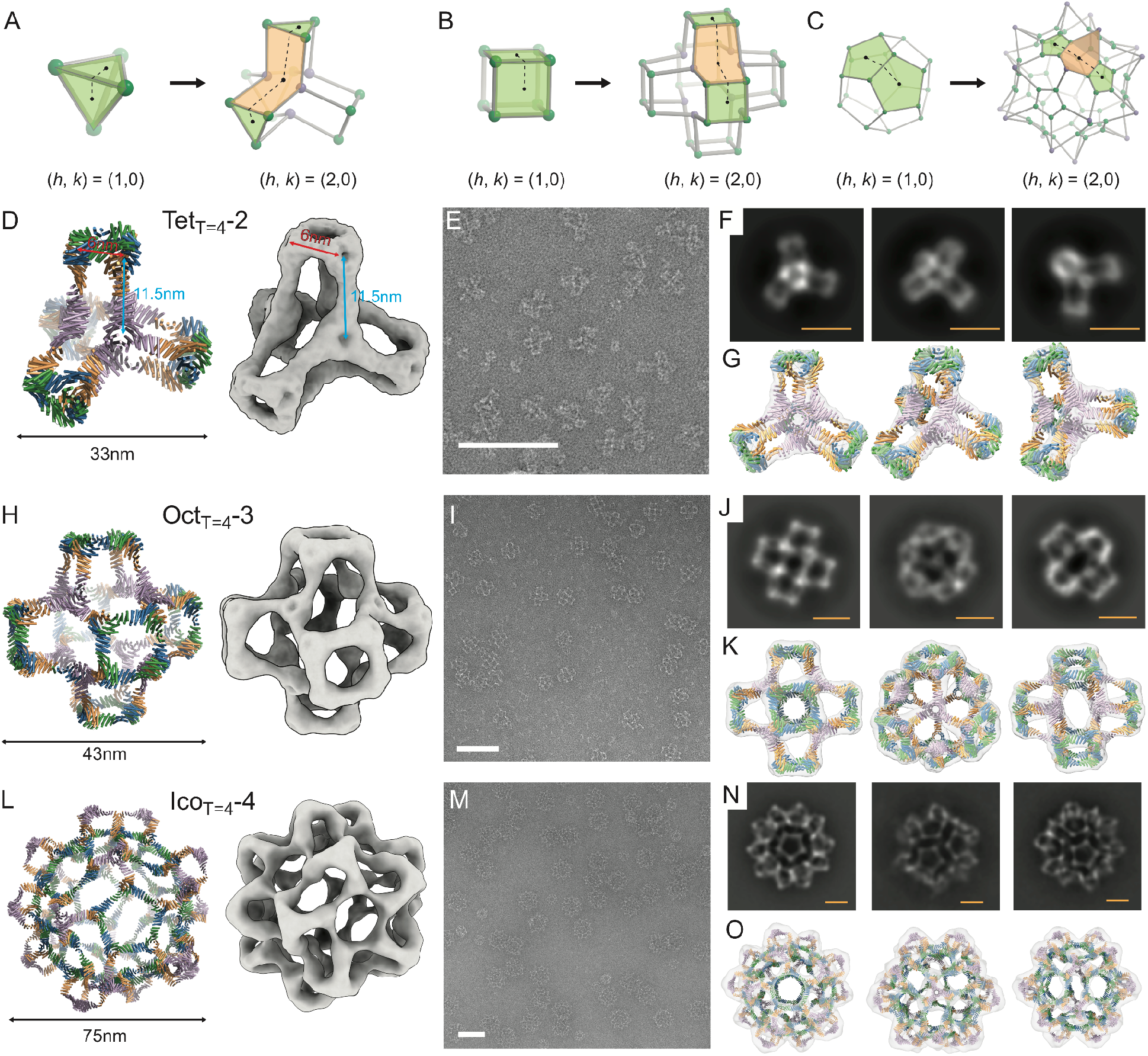
NsEM characterization of designed T = 4 tetrahedral, octahedral, and icosahedral protein cages. **(A-C)** Ball-and-stick models of (left) T=1 and (right) T=4 cages for each symmetry, defined by the Caspar and Klug nomenclature ^8^. Four component T = 4 (**D**-**G**) tetrahedral cage with 48 subunits, (**H**-**K**) octahedral cage with 96 subunits, (**L**-**O**) icosahedral cage with 240 subunits. (**D, H, L**) (left) Structure models of the T=4 cages and (right) 3D reconstructed nsEM map. Each color of the model represents each component (green: ch_A, blue: ch_B, orange: ch_C, purple: ch_ho). (**E, I, M**) nsEM micrographs. (**F, J, N**) Characteristic 2D class averages of nsEM. (**G, K, O**) Superpositions of 3D reconstructed nsEM map (gray density) to the structure model of cages (colors). Scale bars: (white) 100 nm and (yellow) 20 nm.

The T = 4 octahedral cage (Oct_T=4_-3) has a 3D cross shape structure (diameter: 43 nm) with the original cubic shape of the T=1 structure repeated 6 times and shifted away from the origin to positive and negative values of the three 4 fold symmetry axes along x, y and z (Fig. 4B and 4H). Six C4 crowns form the outward faces of the structure along the x, y, z axes which are connected by eight homotrimers placed in a cubic arrangement closer to the center of the cage; as for the tetrahedral cage, the homotrimers and heterotrimers face in opposite directions (Fig. S40). The overall architecture is that of a T = 4 Goldberg polyhedra with octahedral symmetry^26^, with an elongated hexagon bridging the square faces (Fig. 4B) and a 10 nm cavity in the center. Homogeneous populations of cages are observed in nsEM micrographs (Fig. 4I), and 2D class averages along the 2, 3 and 4 fold symmetry axes are closely consistent with the design model (Fig. 4J). The nsEM 3D reconstruction is very close to the overall structure (Fig. 4K), but a slightly curved connection between crowns and homotrimers leading to a slight twist of the overall structure (Fig. S40). To characterize the structure at higher resolution, we collected cryoEM datasets of the Oct_T=4_-3 cage and generated a 3D reconstruction which following refinement resulted in a 3D cryoEM map with 6.87 Å resolution with clear secondary structure features (Fig. 5A-5D). Following relaxation via molecular dynamics^27^, the design model fits well into the cryoEM map (Fig. 5E-5G), with the crown and homotrimer substructures and the individual chains clearly defined (Fig. S41). Around the 4-fold symmetry axes, ch_A and ch_B of the hetBGL0-18-17_A32 heterotrimers (Fig. 5B and 5E, green and blue) form square motifs; the 5 helices in each DHR-arm in both chains are clearly evident in the cryoEM map (Fig 5E). BGL17_A32 homotrimers are placed along the 3-fold symmetry axes (Fig. 5D); the DHR arm of each subunit has 6 helices and at the end forms an interface with ch_C of the heterotrimer (Fig. 5F and 5G, purple and orange). The twist of the C4 crown substructures relative to the design model arises from shifts at the homotrimer-heterotrimer interface (Fig. S41).

**Fig. 5.**
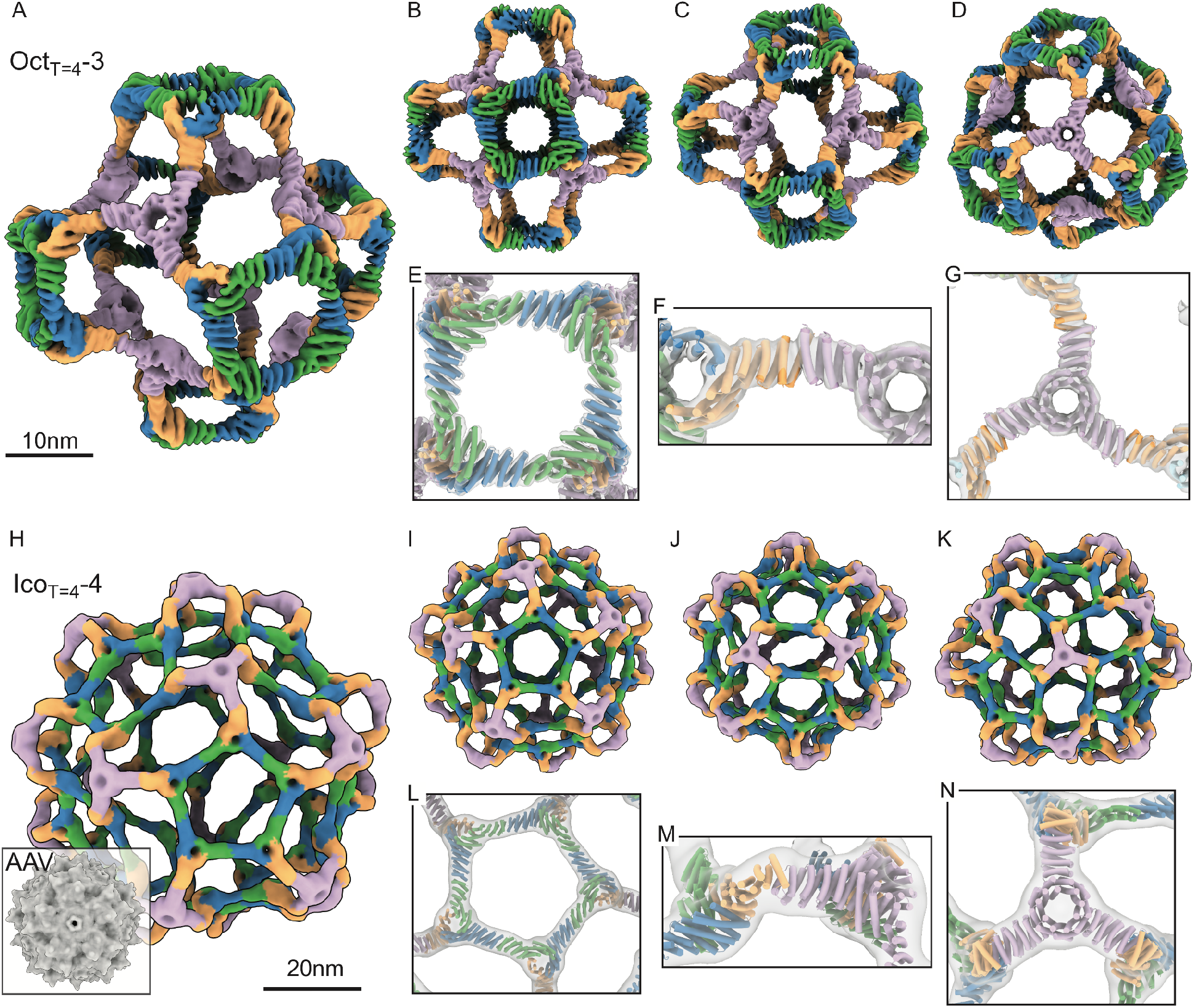
CryoEM characterization of T = 4 octahedral and icosahedral protein cages. (**A**-**D**) 3D cryoEM map of Oct_T=4_-3 from different views. (**E**-**G**) Overlay between the cryoEM map (gray transparent) and the protein model relaxed into the map (colors), for each substructure: (E) C4 crown, (F) the homotrimer-heterotrimer interface, and (G) homotrimer. (**H**-**K**) 3D cryoEM map of Ico_T=4_-4 from different views. The inset of (H) is Adeno-associated virus (AAV) capsid shown for size comparison. (**L**-**N**) Overlay between the cryoEM map (continuous density) and the relaxed protein model (colors), for each substructure: (L) C5 crown, (M) the homotrimer-heterotrimer interface, and (N) homotrimer. Both the cryoEM map and protein model are colored by components (green: ch_A, blue: ch_B, orange: ch_C, purple: ch_ho).

The T = 4 icosahedral cage (Ico_T=4_-4) consists of twelve C5 crowns (pentons) connected by twenty outward facing homotrimers (Fig. 4C, 4L). Largely homogeneous 75 nm size cages were identified by nsEM and dynamic light scattering (Fig. 4M and S44). SDS-PAGE showed clear bands corresponding to each component suggesting that all four chains are present (Fig S43). The 2D class averages (Fig. 4N) and 3D nsEM reconstructions (Fig. 4L, 4O) have the overall designed shape but the orientations of the C5 crowns and homotrimers appeared inverted from the design model (Fig. S47). We collected cryoEM images, and 3D reconstruction and refinement of the cryoEM data yielded a 3D cryoEM map with 13.15 Å resolution (Fig. 5H-5K), in which the holes at the center of trimers and the orientations of trimers are clearly identified. The design model with inverted components fits well into the cryoEM density following relaxation (Fig. 5L-5N; Fig. S48). The overall structure has the architecture of a T = 4 Goldberg polyhedron with icosahedral symmetry^26^, with boat-type hexagonal motifs placed between pentagons (Fig. 4C). Surrounding the 5-fold symmetry axes are pentons formed from hetBGL0-18-17_A32 heterotrimers (Fig 5H and 5I, green, blue and orange). The pentons are bridged by BGL17_A32 homotrimers which form tripod-like protrusions on the 3-fold symmetry axis (Fig 5H and 5K, purple). On the two fold axes are boat-type distorted hexagons with two homotrimers and four heterotrimers on the vertices; two of the edges are formed by interacting heterotrimer subunits, and four edges by interacting homotrimer and heterotrimer subunits (Fig 5J). The outer diameter of the Ico_T=4_-4 cage is about three times larger than that of Adeno-associated virus (AAV) capsid (Fig. 5H), and the inner diameter of the empty pore at the center of the Ico_T=4_-4 cage is ∼50 nm (volume ∼ 6.55 x 10^4^ nm^3^), which can be used to package diverse cargos.

## Conclusion

Our T = 4 assemblies are a considerable advance in compositional and structural complexity over previous nanocage designs which have one or two distinct components and up to three distinct interfaces: the designs described here have four distinct core structural components and six distinct interfaces. The success of the design strategy required that the six interfaces be orthogonal – this is achieved by using distinct hydrogen bond networks^18,28^ within the heterotrimers, and different arrangements of interacting helices at the heterotrimer-heterotrimer and heterotrimer-homotrimer interfaces (Fig S33). Our design approach is not limited to T = 4 cages but can be extended to higher T; for example T = 7 cages with 420 subunits can be constructed by incorporating a second heterotrimer – either based on another BGL homodimer, or using a different set of designed heterotrimers^17^(see Fig. S49, S50 for routes to T = 7 and T = 9). To our knowledge, pseudo-symmetric tetrahedral and octahedral assemblies have not been observed in Nature; with four distinct chains and 8 termini available for functionalization, these provide starting points for sophisticated multicomponent materials and crystals^29^. With an 75 nm diameter, our T=4 icosahedral design has considerably more interior volume available for packaging nucleic acid and other cargoes than previously designed 1 and 2 component cages; the accompanying Dowling et al. manuscript describes a route to even larger albeit less homogenous structures using two component heterotrimers^30^. For vaccine and delivery applications, the four distinct structural components of our T = 4 assemblies provide many more opportunities for further functional elaboration than the one and two component nanoparticles designed to date: for vaccines, presentation of multiple antigens and immunomodulators, and for targeted delivery, incorporation of distinct designed modules for targeting and endosomal escape.

## Acknowledgements

We thank Erin C. Yang, Quinton M. Dowling, Neil P. King for helpful discussions; Helen Eisenach for advice on explicit negative design; Xinting Li for help with mass spectrometry analysis of proteins; Caixuan Liu for advice on CryoEM data processing. Native mass spectrometry measurements were provided by Florian Busch, Andrew Norris, Nicholas Horvath, and Sean Cleary of the NIH-funded Resource for Native Mass Spectrometry Guided Structural Biology at The Ohio State University (NIH P41 GM128577 awarded to Vicki Wysocki). Small-angle X-ray scattering was conducted at the Advanced Light Source (ALS), a national user facility operated by Lawrence Berkeley National Laboratory on behalf of the Department of Energy, Office of Basic Energy Sciences, through the Integrated Diffraction Analysis Technologies (IDAT) program, supported by DOE Office of Biological and Environmental Research. Additional support comes from the National Institute of Health project ALS-ENABLE (P30 GM124169) and a High-End Instrumentation Grant S10OD018483. This research used resources of the National Energy Research Scientific Computing Center (NERSC), a U.S. Department of Energy Office of Science User Facility located at Lawrence Berkeley National Laboratory, operated under Contract No. DE-AC02-05CH11231 using NERSC award BER-ERCAP0022018.

## Author contributions

Conceptualization and design protocol: S. Lee, R. D. Kibler, Y. Hsia, D. Baker; Computational design of oligomers: R. D. Kibler, S. Lee, Y. Hsia; Computational design of cages: S. Lee, R. D. Kibler; Protein synthesis and characterizations: S. Lee, R. D. Kibler; CryoEM: A. J. Borst, A. Philomin, R. D. Kibler, S. Lee; X-ray Crystallography: M. A. Kennedy and B. Stoddard; Supervision: D. Baker; Writing – original draft: S. Lee, R. D. Kiber, D. Baker; All authors read and contributed to the manuscript.

