## Supplementary Material for "Design of four component T=4 tetrahedral, octahedral, and icosahedral protein nanocages through programmed symmetry breaking"

**This file includes:**

**Section 1.** *In silico* design of homo-oligomers
**Section 2.** *In silico* design of pseudo-symmetric hetero-oligomers
**Section 3.** *In silico* design of cages and crowns
**Section 4.** Experimental methods and materials
Figs. S1 to S51
Tables S1 to S16
References

### Section 1. *In silico* design of homo-oligomers

1.1. Helical fuse

The HelixFuse protocol from (*1*) was used to combinatorially fuse together a set of 13 BGLs and 38 DHRs (Table S3) by structurally superimposing terminal helix residues (*2*) in either direction (“AB”: N-terminus of a DHR to the C-terminus of a BGL; “BA”: C-terminus of a BGL to the N-terminus of a DHR), only requiring the DHR to extend outward from the center of the BGL. Up to 11 residues of the BGL and an entire repeat of the DHR were allowed to be deleted during the procedure. After the C-beta atoms are superimposed, a RMSD check across 6 residues was performed to ensure that the fusion results in a continuous helix. Outputs were filtered to remove significant clashes (Rosetta centroid energy < 5.0), ensure at least 3 helices in contact with the overlapped region, and remove fusions with poor shape complementarity (<0.5).

1.2. Sequence design (Rosetta)

Sequence design was carried out with RosettaScripts (*3*) (hufse_sym_hdf5.xml) with sequence choices informed by fragments with similar sequence (via StructProfileMover). Because of the presence of hydrogen bond networks in the BGL interfaces, a scoretype designed to avoid buried unsatisfied hydrogen bond networks was used (*4*) and the locations of HBNets were not allowed to be changed. The interface between the BGL and DHR defined by InterfaceByVector and any residues superimposed over were redesigned and sidechain optimized with a basic fixed backbone relax (sidechain minimize -> repack (no design) -> sidechain minimize). The redesign was done again with a fixed core to optimize surface and boundary. After sequence design, designs were selected with more stringent shape complementarity (sc > 0.55), no more than 50 alalines, and at most 10 continuous apolar residues to identify designs likely to be well behaved and rigid across the junction point. In total, the building block library generated in silico by HF consists of 1556 AB and 589 BA fusions.

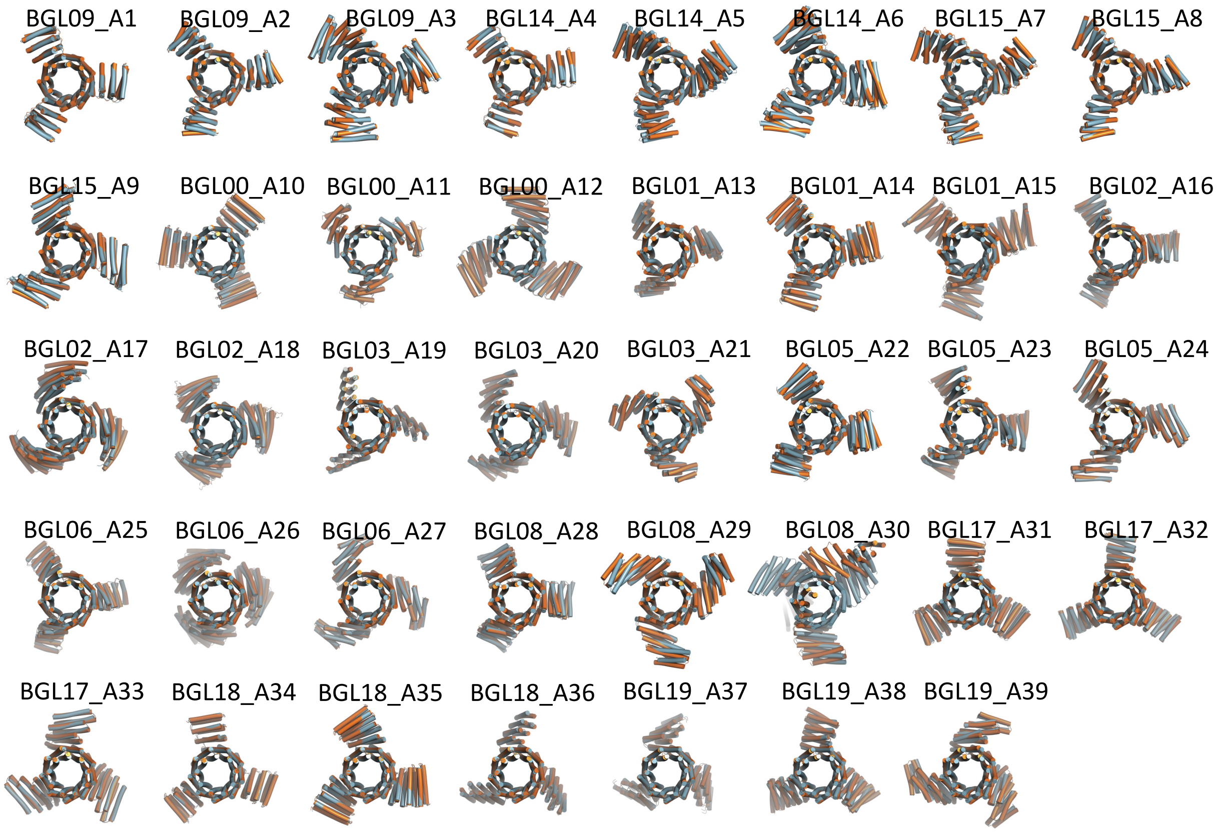


**Fig. S1.** Overlays of model structure (light blue) of homotrimers (BGL09_A01 – BGL19_A39), and their AF2 predictions (orange).

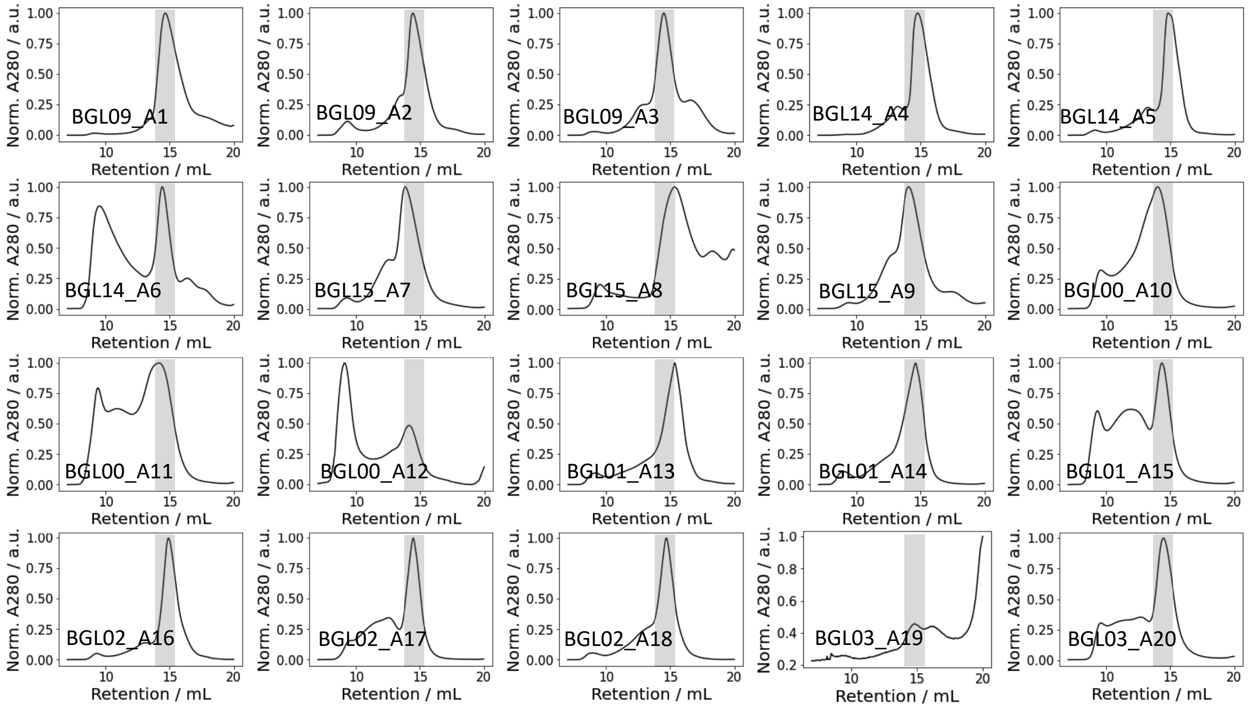


**Fig. S2.** SEC traces of homotrimers (BGL09_A01 – BGL03_A20) obtained using S200 column. Transparent grey box (~ 15ml) of each trace indicates an expected retention volume of the trimers.

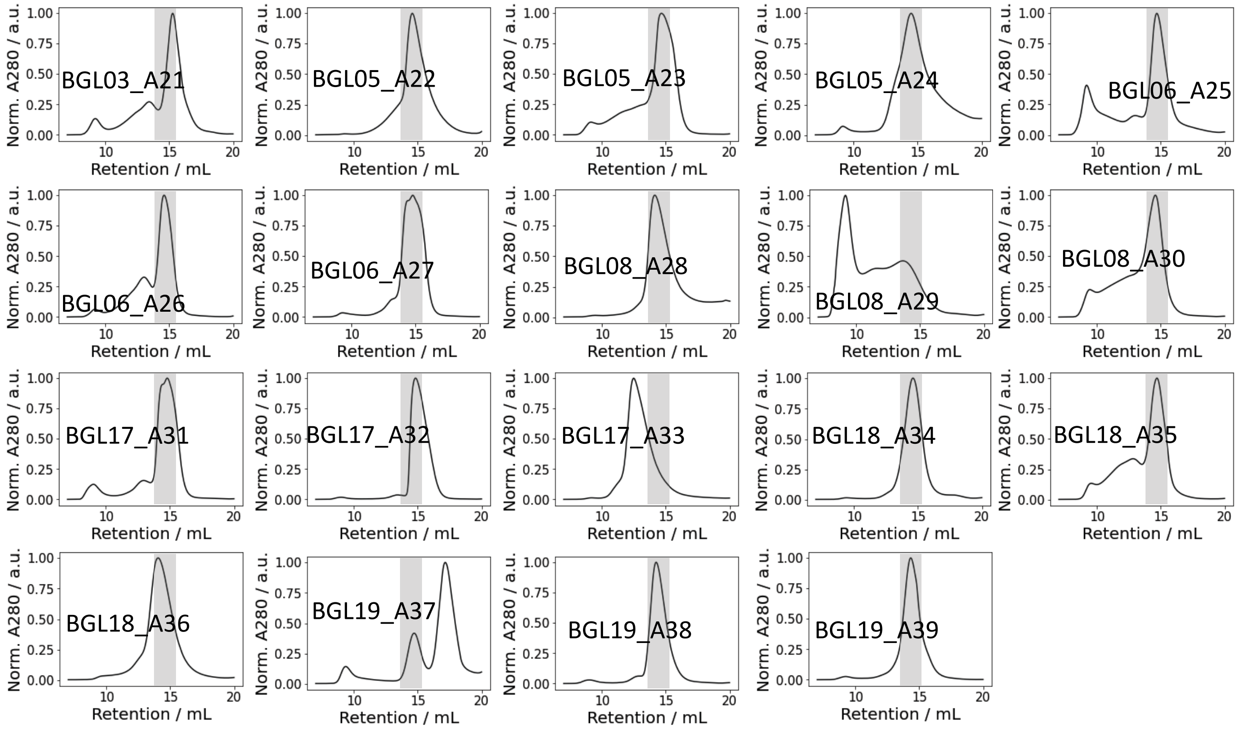


**Fig. S3.** SEC traces of homotrimers (BGL03_A21 – BGL19_A39) obtained using S200 column. Transparent grey box (~ 15ml) of each trace indicates an expected retention volume of the trimers.

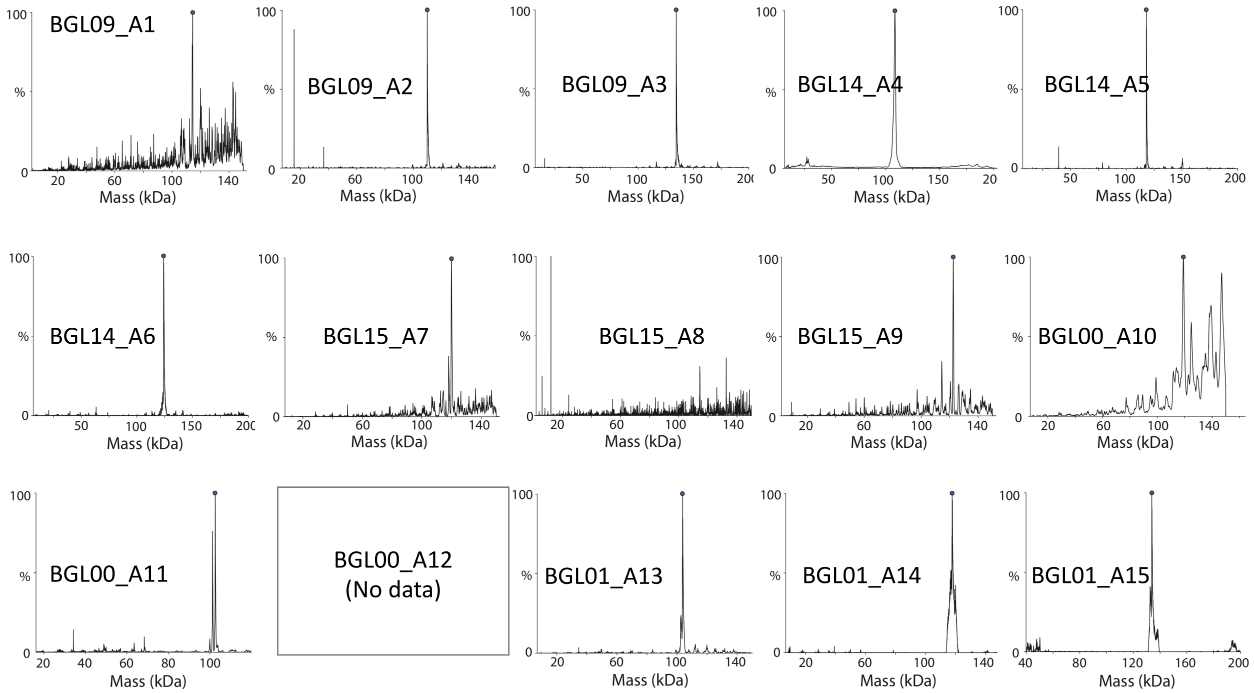


**Fig. S4.** native mass spec results of homotrimers (BGL09_A01 – BGL01_A15). Signals for trimers are marked with a small dot.

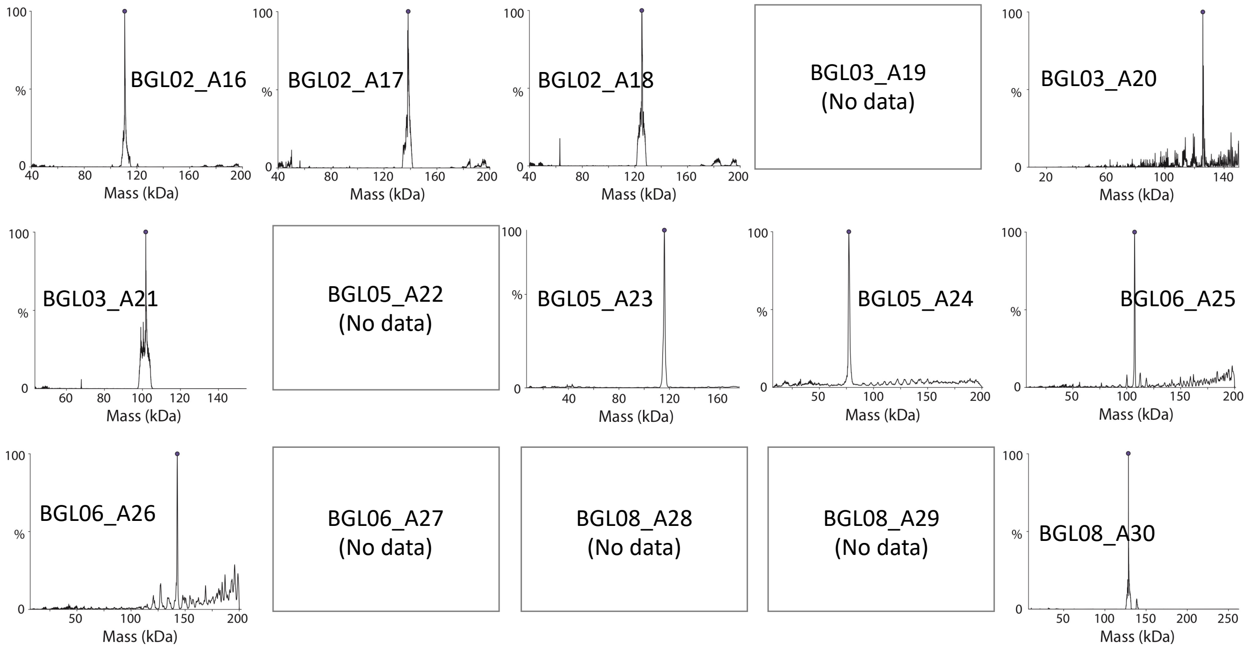


**Fig. S5.** native mass spec results of homotrimers (BGL02_A16 – BGL08_A30). Signals for trimers are marked with a small dot.

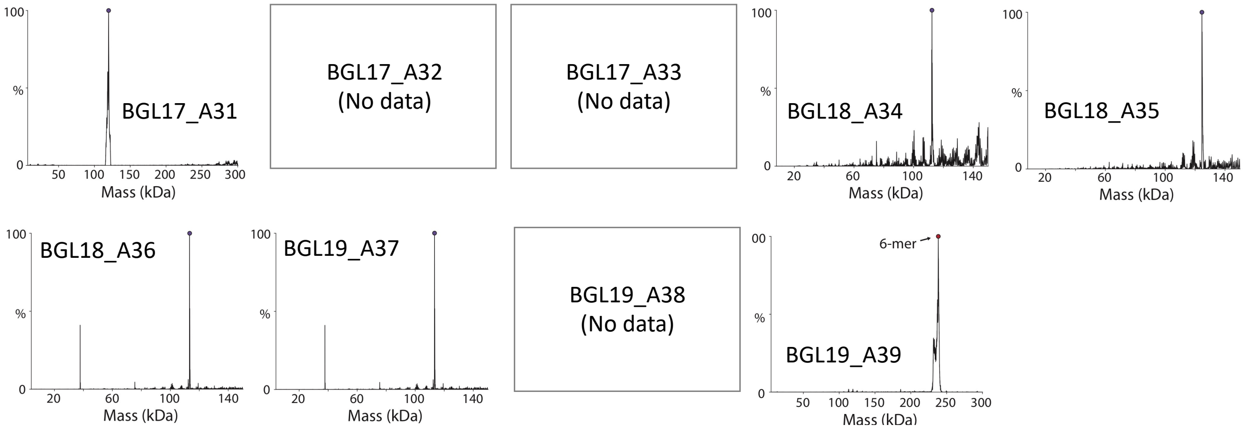


**Fig. S6.** native mass spec results of homotrimers (BGL17_A31 – BGL19_A39). Signals for trimers are marked with a small dot.

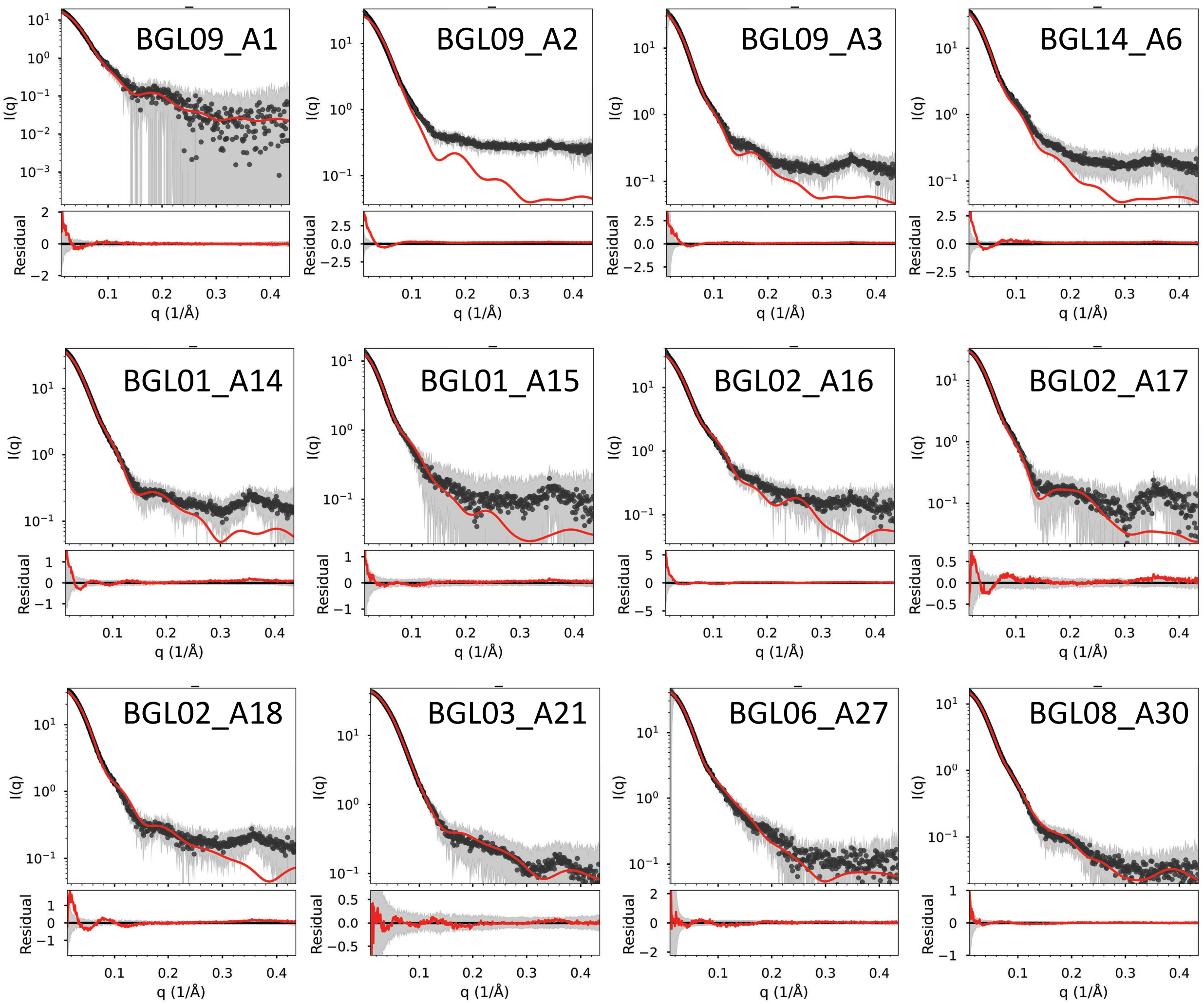


**Fig. S7.** Small angle X-ray scattering (SAXS) results of homotrimers. For each system, top panel is scattering intensity (red: calculated from model, black dots: experimental data, grey: the standard deviation of the averaged frames). (Bottom of each panel) The residual of the fit between experimental data and the computed profile, shown on a linear scale.

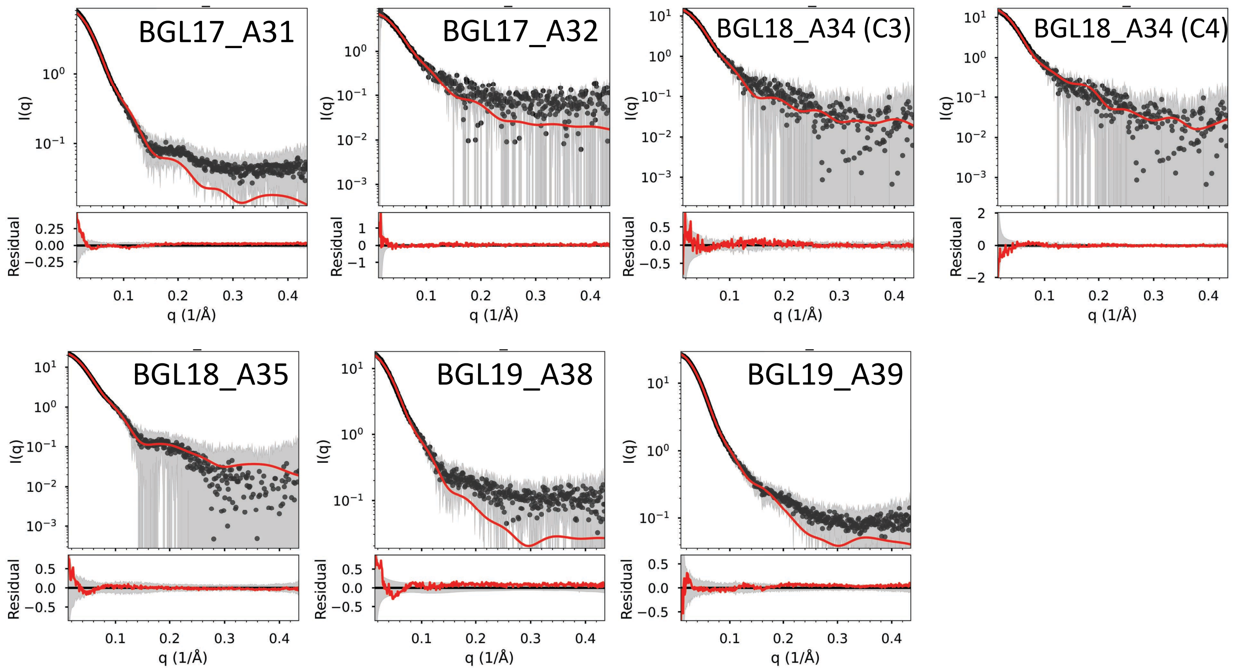


**Fig. S8.** Small angle X-ray scattering (SAXS) results of homotrimers. For each system, top panel is scattering intensity (red: calculated from model, black dots: experimental data, grey: the standard deviation of the averaged frames). (Bottom of each panel) The residual of the fit between experimental data and the computed profile, shown on a linear scale.

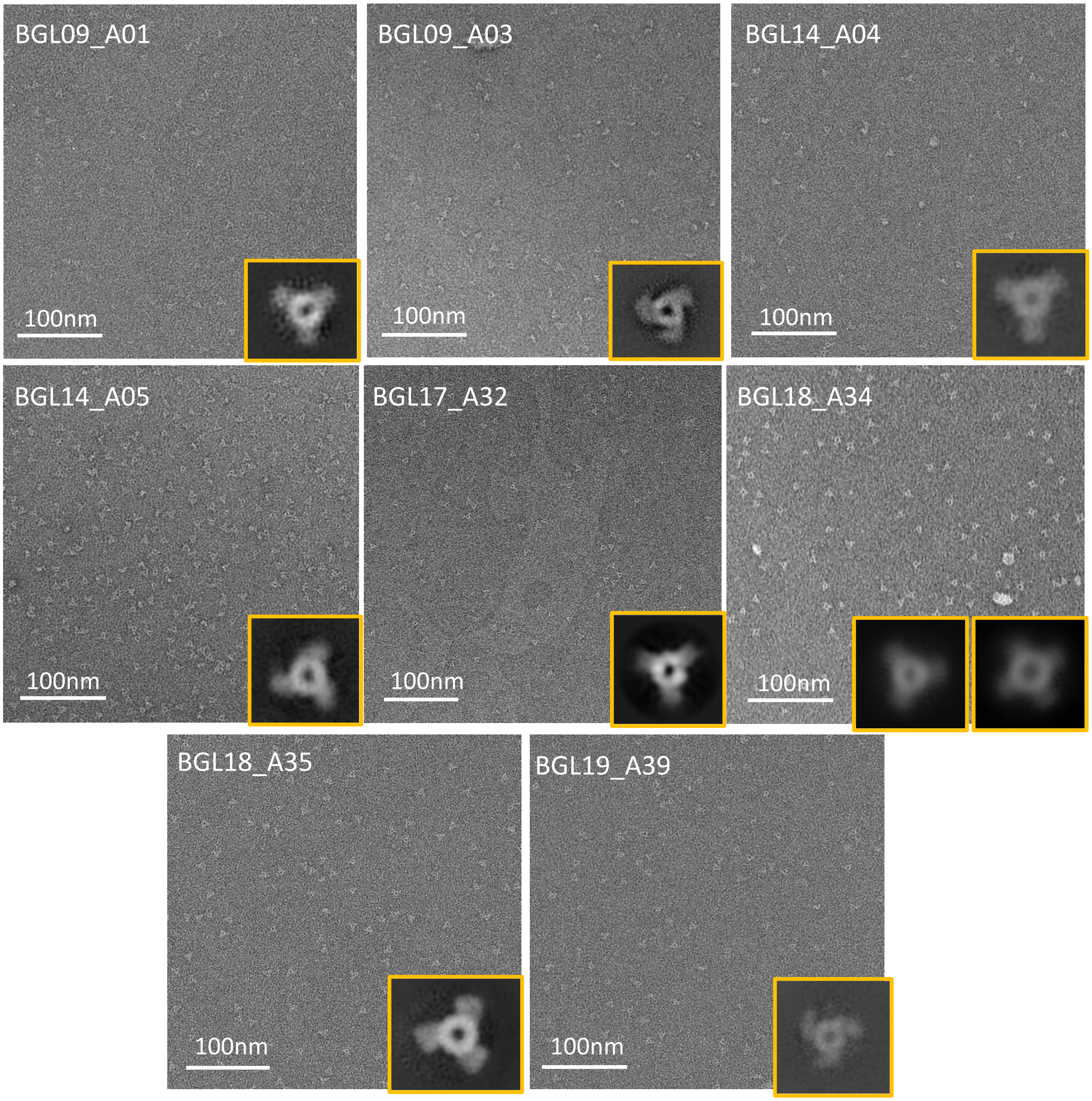


**Fig. S9.** Negative stained electron micrographs (nsEM) of homotrimers (BGL09_A01, BGL09_A03, BGL14_A04, BGL14_A05, BGL17_A32, BGL18_A34, BGL18_A35, BGL19_A39). Insets are 2D average classes along 3-fold symmetry axis of the oligomers. Note that BGL18_A34 is a mixture of C3 and C4 (off-target) oligomers.

### Section 2. *In silico* design of pseudo-symmetric hetero-oligomers

The basic idea of designing pseudo-symmetric hetero-oligomers is to transplant three different protomer-protomer interfaces of BGLs (*5*) (guest) into a homotrimer scaffold (host), which gives three different chains that form a heterotrimer but the same structure with the homotrimer.

We first tested the compatibility between a guest interface and a host scaffold as a homo-oligomer, following steps below:

(i) Identifying residues (of the host) involving in protomer-protomer interfaces.

(ii) Identifying residues (of the host) involving in DHR-arm binding.

(iii) The residues (of the host) identified in (i) but not involving in (ii) are defined as 'mutable residues'.

(iv) Identifying residues (of the BGL guest) involving in protomer-protomer interfaces.

(v) Mutating the 'mutable residues' to the residues in (iv).

A schematic diagram of the interface transplantation process is shown in Fig. S10. After validating interface transplanted homotrimers in experiment (Fig. S12–S13), we combined them into heterotrimers (Fig. S11). Protein sequences of the interface transplanted homotrimers are in Table S4. For a heterotrimer, a host and three guests are selected (Table S1), and the protomer-protomer interface of each guest is transplanted into one of the protomer-protomer interfaces of the host (Fig. S14-S18). Protein sequences of the heterotrimers are in Table S5.

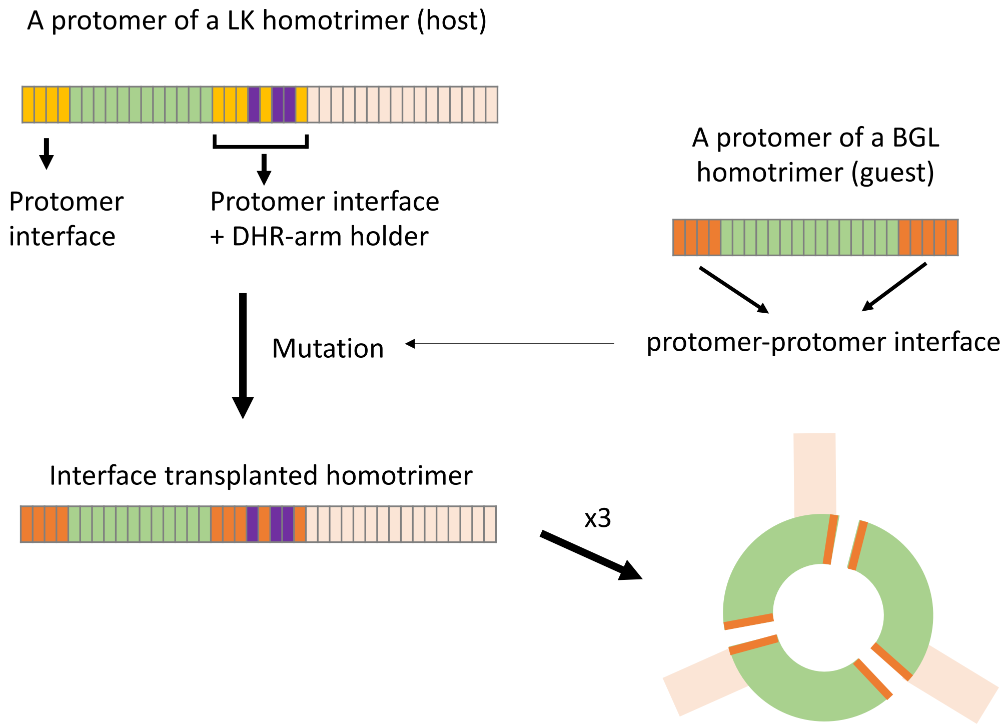


**Fig. S10.** Schematic diagram of the interface transplantation of homotrimers.

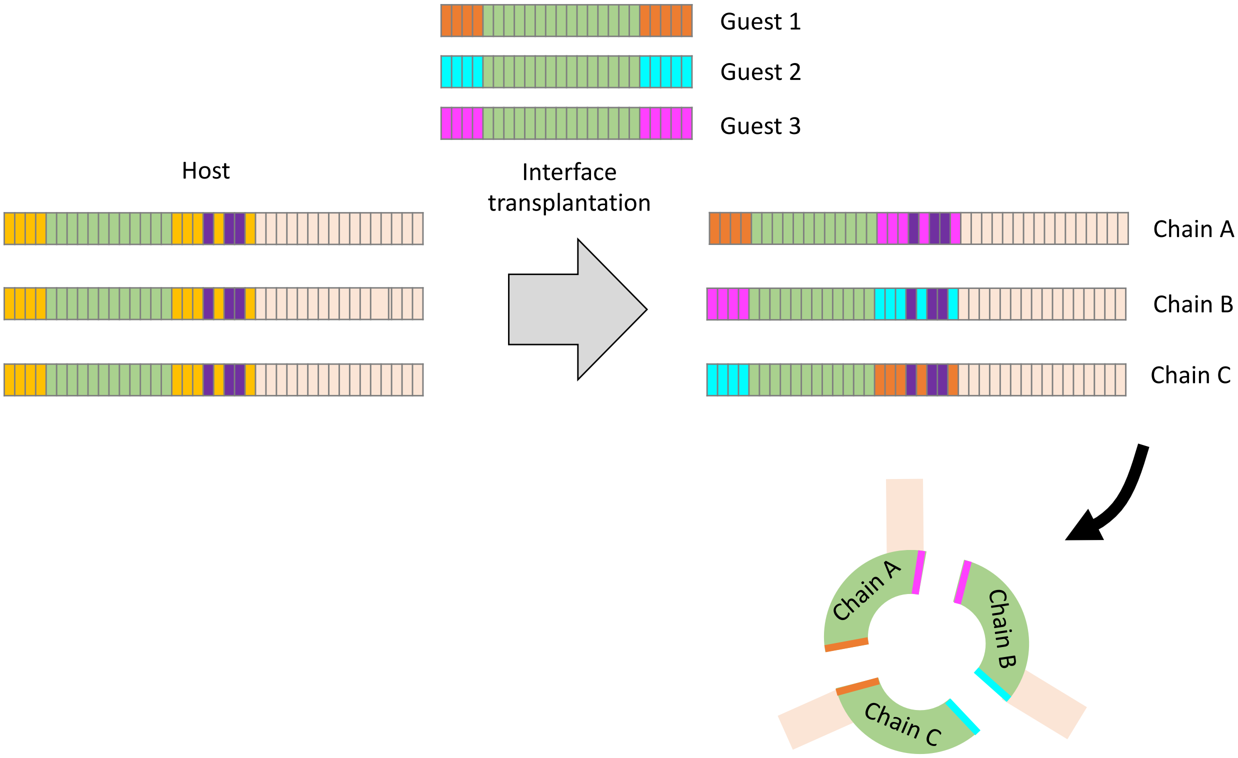


**Fig. S11**. Schematic diagram of the interface transplantation of heterotrimers.

**Table S1.** Host and guest combinations for the interface transplantation of heterotrimers.

| ID | Host | Guest 1 | Guest 2 | Guest 3 |
|---|---|---|---|---|
| hetBGL0-17-18_A32 | BGL17_A32 | BGL00 | BGL17 | BGL18 |
| hetBGL0-17-19_A32 | BGL17_A32 | BGL00 | BGL17 | BGL19 |
| hetBGL0-18-17_A32 | BGL17_A32 | BGL00 | BGL18 | BGL17 |
| hetBGL0-19-17_A32 | BGL17_A32 | BGL00 | BGL19 | BGL17 |
| hetBGL17-18-19_A32 | BGL17_A32 | BGL17 | BGL18 | BGL19 |

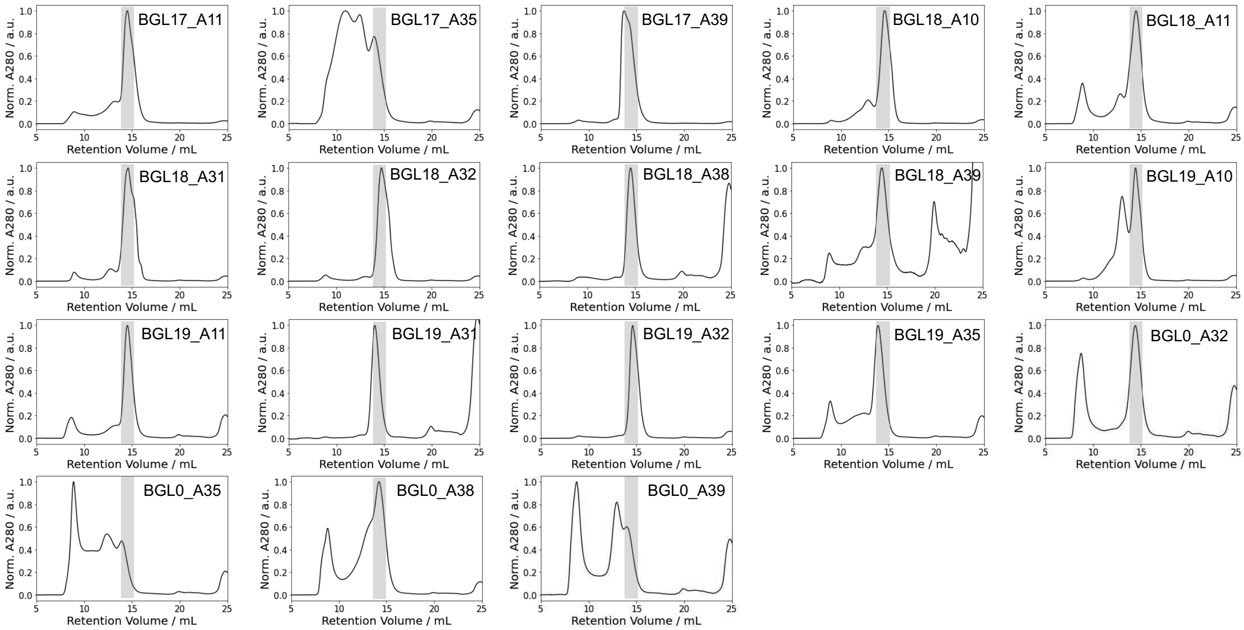


**Fig. S12.** SEC traces of 18 interface-transplanted homotrimers obtained using S200 column. Transparent grey box (~ 15ml) of each trace indicates an expected retention volume of the trimers.

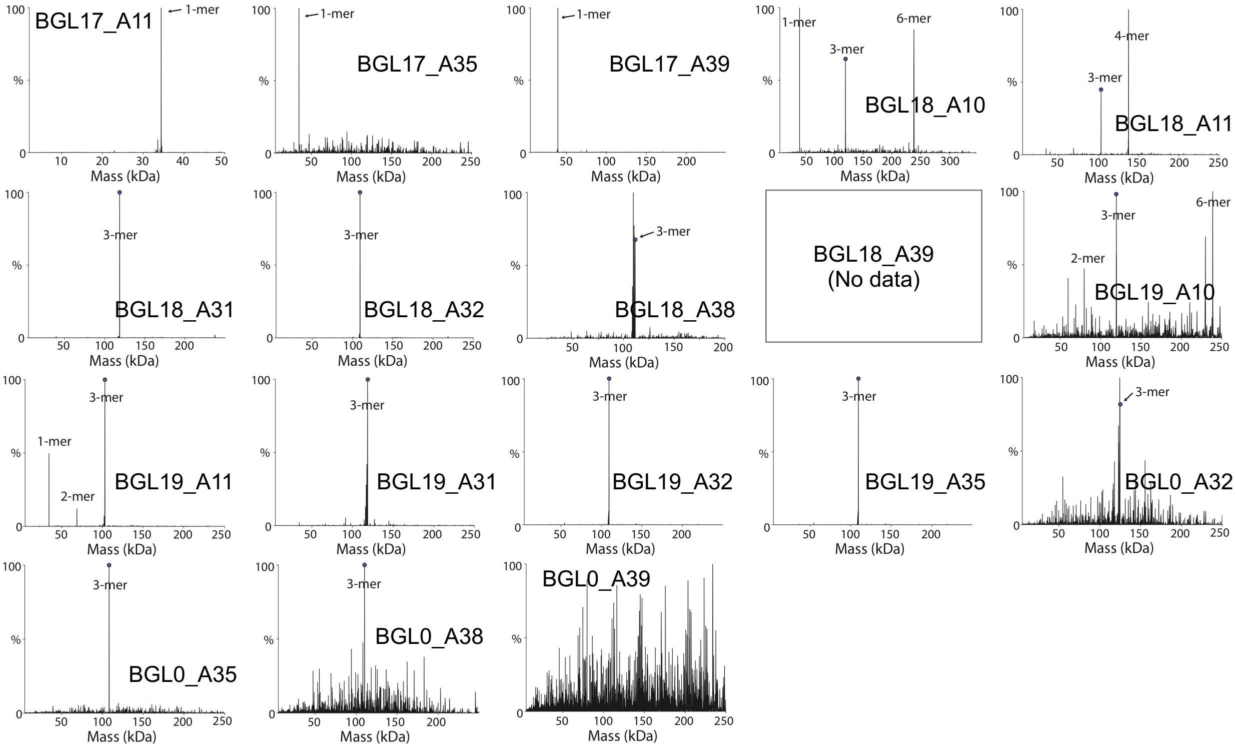


**Fig. S13.** Native mass spec results of interface-transplanted homotrimers. Signals for trimers are marked with a small dot.

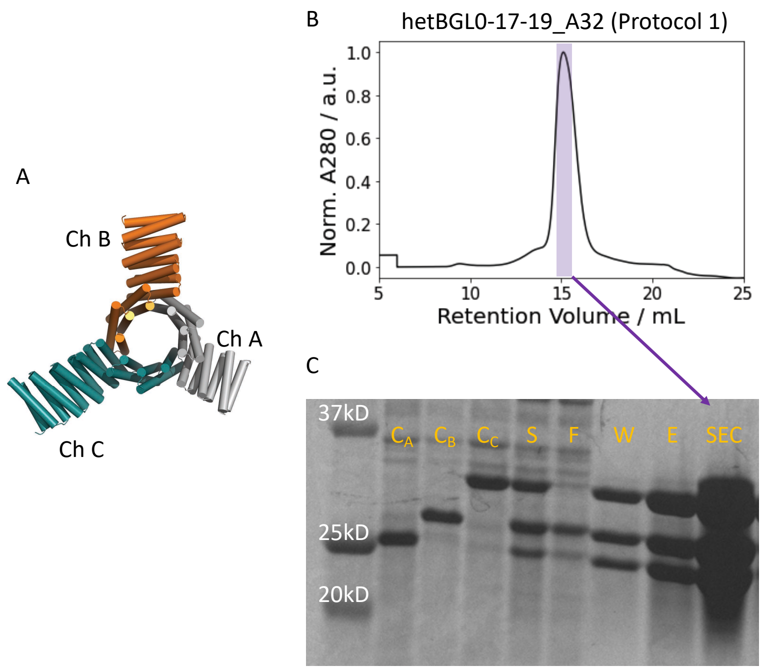


**Fig. S14.** Pseudo-symmetric heterotrimers (hetBGL0-17-19_A32). (**A**) Model structure. (**B**) SEC trace and (**C**) SDS-PAGE results from protocol 1. (**D**) SEC trace and (**E**) SDS-PAGE results from protocol 2. (**C**, **E**) C: E. Coli culture. S: Soluble fraction. F: Flow through from IMAC. W: Washed fraction. E: Elution. SEC: SEC peak. $C_X$ indicates E. Coli culture of chain-X before mixing with other chains.

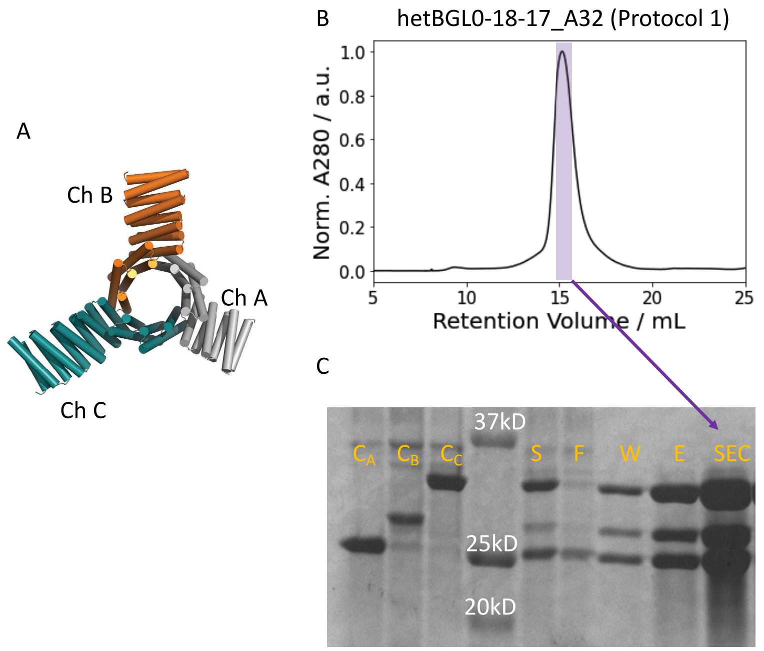


**Fig. S15.** Pseudo-symmetric heterotrimers (hetBGL0-18-17_A32). (**A**) Model structure. (**B**) SEC trace and (**C**) SDS-PAGE results from protocol 1. (**D**) SEC trace and (**E**) SDS-PAGE results from protocol 2. (**C**, **E**) C: E. Coli culture. S: Soluble fraction. F: Flow through from IMAC. W: Washed fraction. E: Elution. SEC: SEC peak. $C_X$ indicates E. Coli culture of chain-X before mixing with other chains.

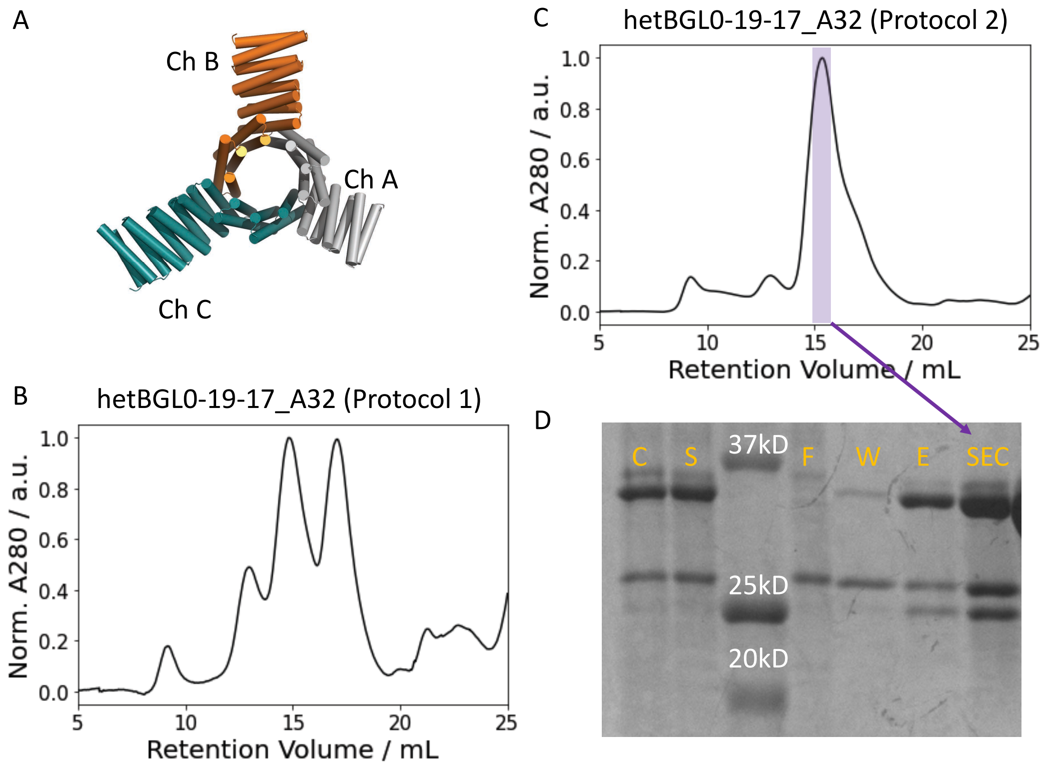


**Fig. S16.** Pseudo-symmetric heterotrimers (hetBGL0-19-17_A32). (**A**) Model structure. (**B**) SEC trace from protocol 1. (**C**) SEC trace and (**D**) SDS-PAGE results from protocol 2. (**D**) C: E. Coli culture. S: Soluble fraction. F: Flow through from IMAC. W: Washed fraction. E: Elution. SEC: SEC peak.

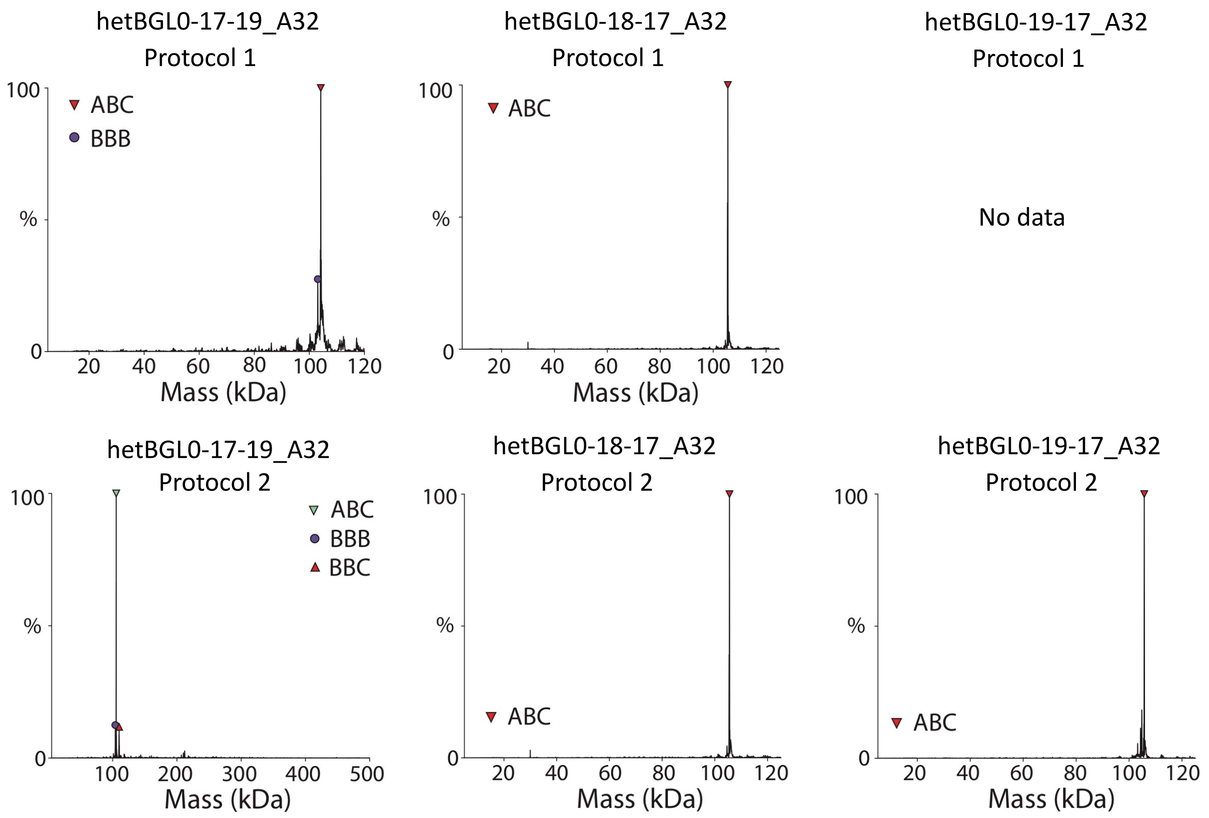


**Fig. S17.** Native mass spec results of pseudo-symmetric heterotrimers (hetBGL0-17-19_A32, hetBGL0-18-17_A32, hetBGL0-19-17_A32) made from (top row) protocol 1 and (bottom row) protocol 2.

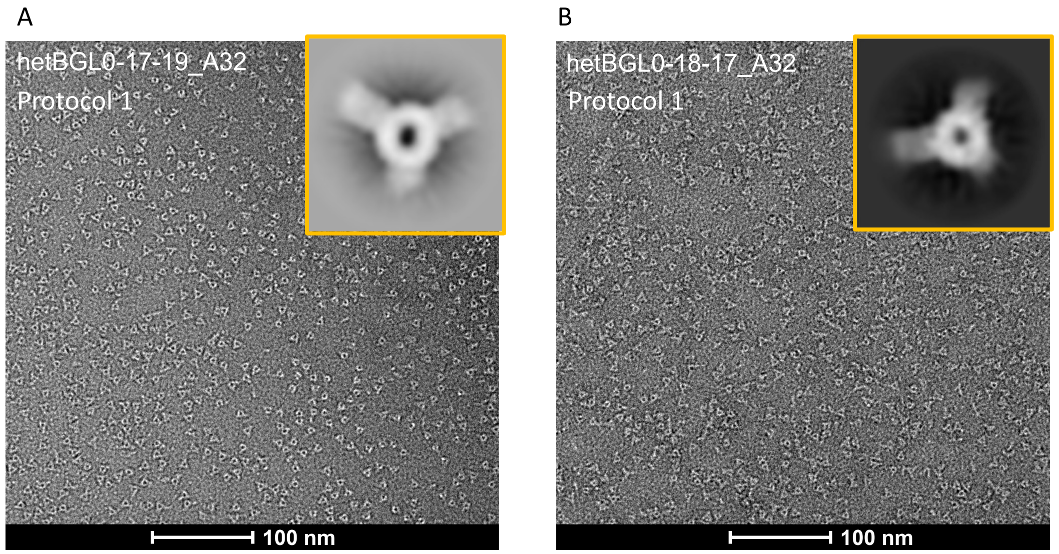


**Fig. S18.** nsEM of pseudo-symmetric heterotrimers (hetBGL0-17-19_A32 and hetBGL0-18-17_A32) obtained from protocol 1. Insets are 2D average classes along 3-fold symmetry axis of the oligomers.

### Section 3. *In silico* design of cages and crowns

3.1. Docking

Cages docking was carried out using the residue-pair transform docking (RPXdock) which is available at https://github.com/willsheffler/rpxdock, and details of the RPXdock can be found in a recently published paper (*6*). Briefly, the RPXdock samples docking configurations using input building blocks for a given symmetry by translating and rotating the building blocks along given symmetry axes. For T = 1 cages, T3, O3 and I3 symmetries were used for tetrahedral, octahedral, and icosahedral symmetric cages, respectively. Input scaffolds are BGL17_A32-based backbones, which are the same backbones with $Tet_{T=1}$-4 (for T3), $Oct_{T=1}$-2 (for O3), and $Ico_{T=1}$-1 (for I3). For crowns, we did not run additional dockings. Instead, we reused the docking interfaces verified from T = 1 cages. For T = 4 cages, T33, O43 and I53 symmetry were used. For T33, input scaffolds are: (1) C3 crown (made by backbones of $Tet_{T=4}$-2_chA, $Tet_{T=4}$-2_chB, $Tet_{T=4}$-2_chC) along a 3-fold symmetry axis, (2) a homotrimer (made by backbones of $Tet_{T=4}$-2_ho) along another 3-fold symmetry axis. For O43, input building blocks are: (1) C4 crown (made by backbones of $Oct_{T=4}$-3_chA, $Oct_{T=4}$-3_chB, $Oct_{T=4}$-3_chC) along a 4-fold symmetry axis, (2) a homotrimer (made by backbones of $Oct_{T=4}$-3_ho) along a 3-fold symmetry axis. For I53, input building blocks are: (1) C5 crown (made by backbones of $Ico_{T=4}$-4_chA, $Ico_{T=4}$-4_chB, $Ico_{T=4}$-4_chC) along a 5-fold symmetry axis, (2) a homotrimer (made by backbones of $Ico_{T=4}$-4_ho) along a 3-fold symmetry axis. Each docking configuration is scored by RPX-score, and dockings with top 3 scores for each symmetry were selected to carry out sequence design.

3.2. Sequence design

Sequence design for the cages was carried out using a deep learning based software ProteinMPNN which is available at https://github.com/dauparas/ProteinMPNN, and details (algorithms etc.) are well described in a recently published paper (*7*). ProteinMPNN was used for designing only for residues forming the protein-protein interfaces for the cages or crowns (Table S6-S14). For T=1 cage design, protomer sequences were tied to be identical to design homo-dimeric interfaces, and cysteines were disallowed. For crown design, we reused the docking interfaces verified in T=1 cages, but now protomer sequences were not tied to design hetero-dimeric interfaces, and cysteines were disallowed. For T=4 cage, the docking interface between chain C of crowns and homo-trimer were designed, and designed sequences were not tied to design hetero-dimeric interfaces. Cysteines were disallowed. For each design, 32 sequences for each temperature (0.1, 0.2, 0.4, 0.6, 0.8 and 1.0) were generated.

The quality of the newly designed interfaces was assessed by AlphaFold2 (AF2) (*8*) using model 4 and $N_{recycle} = 8$ (https://github.com/deepmind/alphafold). The new interfaces for cages and crowns are located at DHR-arm part (not the BGL ring part), thus we conducted AF2 only with the DHR-arm parts instead of running entire protomers to reduce computation time. For homo-dimeric interface design, we selected designs that show backbone RMSD < 2.0Å (computed with pymol) to the original protomer in AF2 prediction and $N_{Met} < 2$ at the interface for experimental test. For hetero-dimeric interface designs, we additionally checked the self-interacting of the designed interface of each chain, using AF2 (Fig. S19). We only used the designs that show a weak self-interaction (Fig. S19).

For T=4 octahedral cage design, we performed explicit negative design implemented in proteinMPNN (*7*). If AF2 predicts the formation of self interface (*i.e.* homodimeric interface), we provided the self interface scaffold as an input of the negative design and ran the sequence design again to decrease probability of the formation of the self interface. We used the beta value = -0.5, which is a coefficient for the negative design in proteinMPNN.

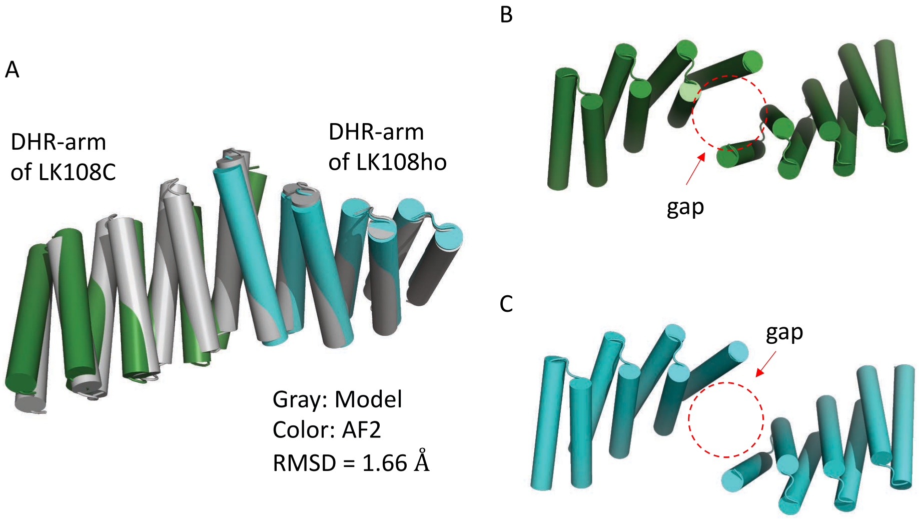


**Fig. S19. An example of computational validation of designed interface using AF2**. Only DHR-arm part was used for AF2 prediction. (**A**) Interface between $Oct_{T=4}$-3_chC and $Oct_{T=4}$-3_ho. Model (Gray) and AF2 prediction (green and cyan). (**B-C**) In self-association check, AF2 predict that the homo-dimeric interface of (**B**) $Oct_{T=4}$-3_chC and (**C**) $Oct_{T=4}$-3_ho show a gap at the interface. We assumed that this gap indicates a weak (or no) self-interaction. For all other hetero-dimeric interface designs of this work, we only used the interface designs that show the weak self-interaction.

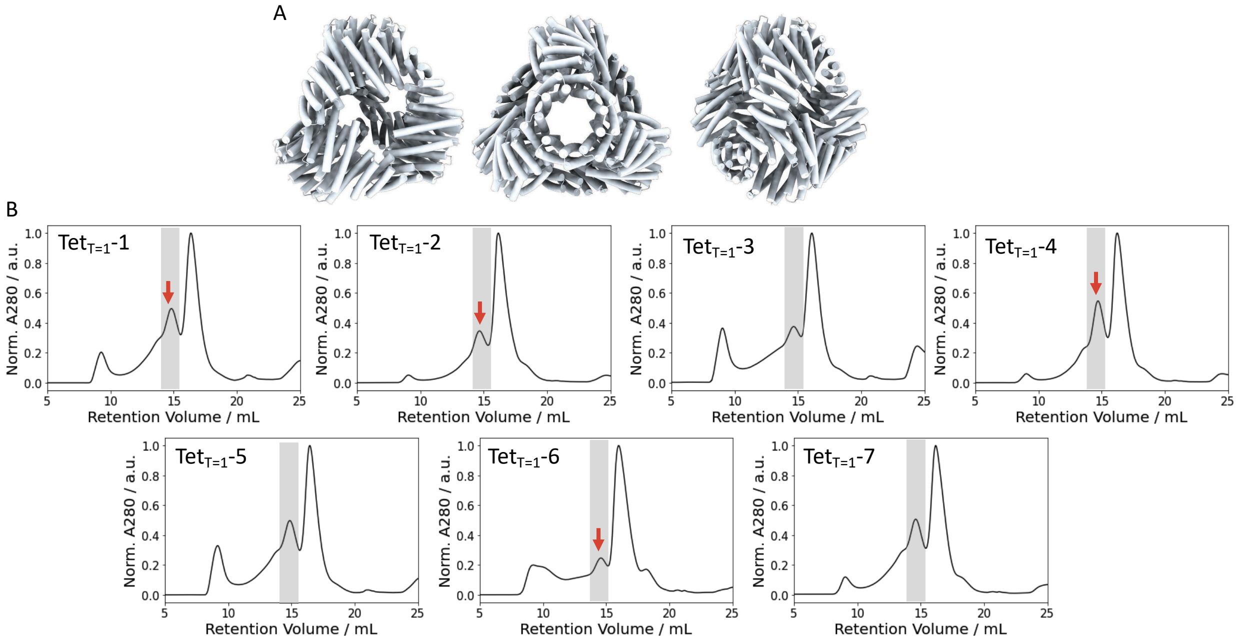


**Fig. S20**. (**A**) Structure model of T=1 tetrahedral cages (symmetry: T3). (**B**) SEC traces of the cages ($Tet_{T=1}$-1 – $Tet_{T=1}$-7) obtained from S6 column. Transparent grey box (~ 15ml) of each trace indicates an expected retention volume of the cage. Red arrows indicate where nsEM in Fig. S21 were obtained.

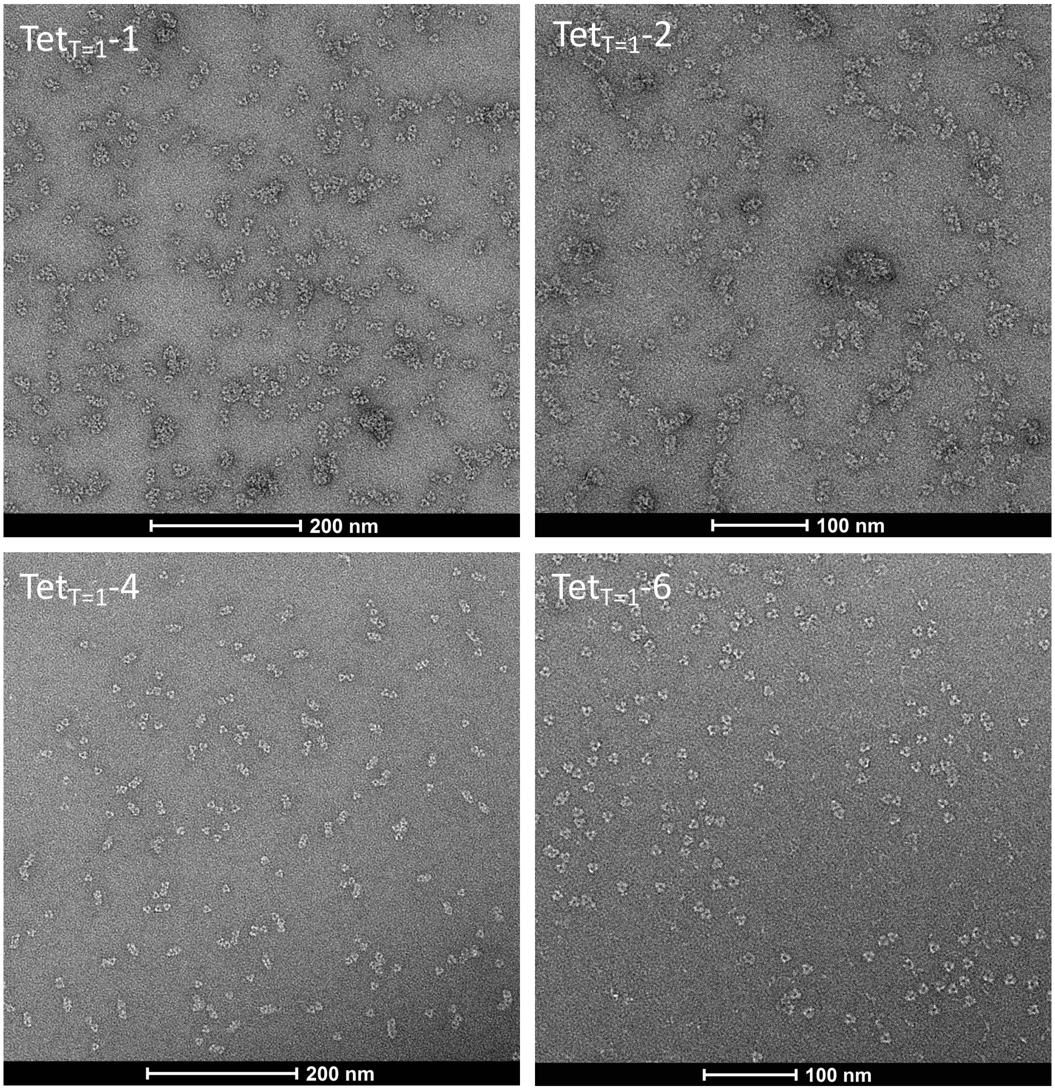


**Fig. S21.** nsEM images of tetrahedral cages (T=1, Symmetry: T3) ($Tet_{T=1}$-1, $Tet_{T=1}$-2, $Tet_{T=1}$-4, $Tet_{T=1}$-6) obtained from SEC peaks shown in Fig. S20 (red arrows).

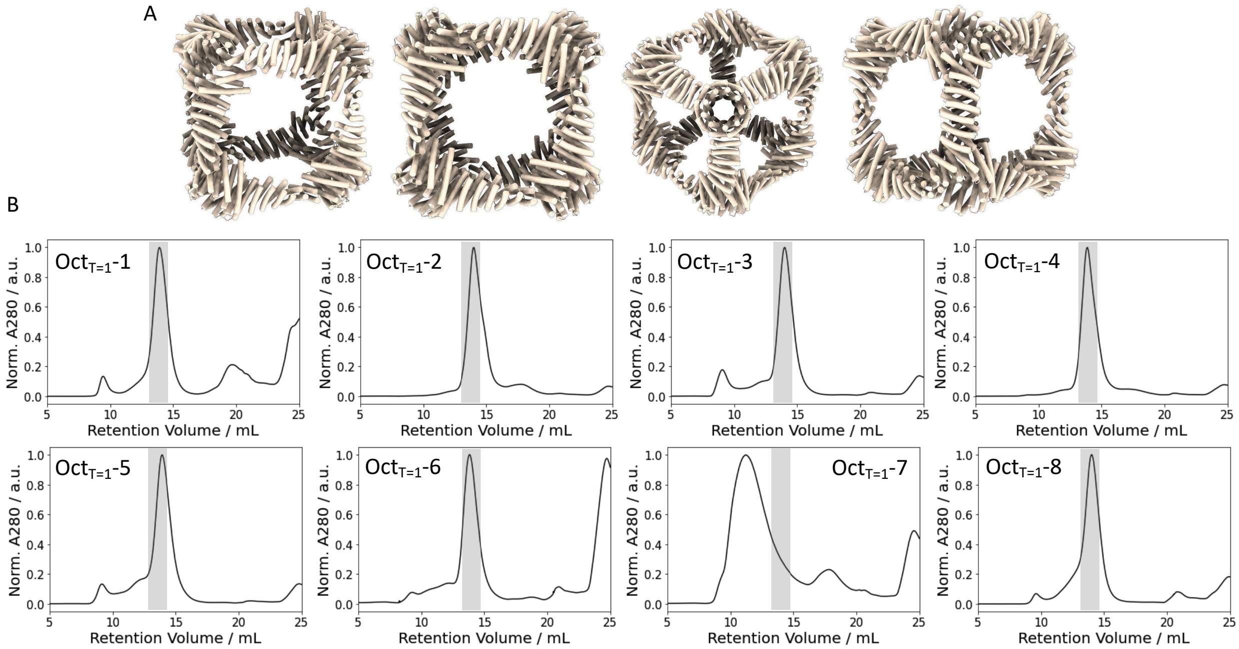


**Fig. S22**. (**A**) Structure model of T=1 octahedral cages (symmetry: O3). (**B**) SEC traces of the cages (Oct$_{T=1}$-1 – Oct$_{T=1}$-8) obtained from S6 column. Transparent grey box (~ 13ml) of each trace indicates an expected retention volume of the cage.

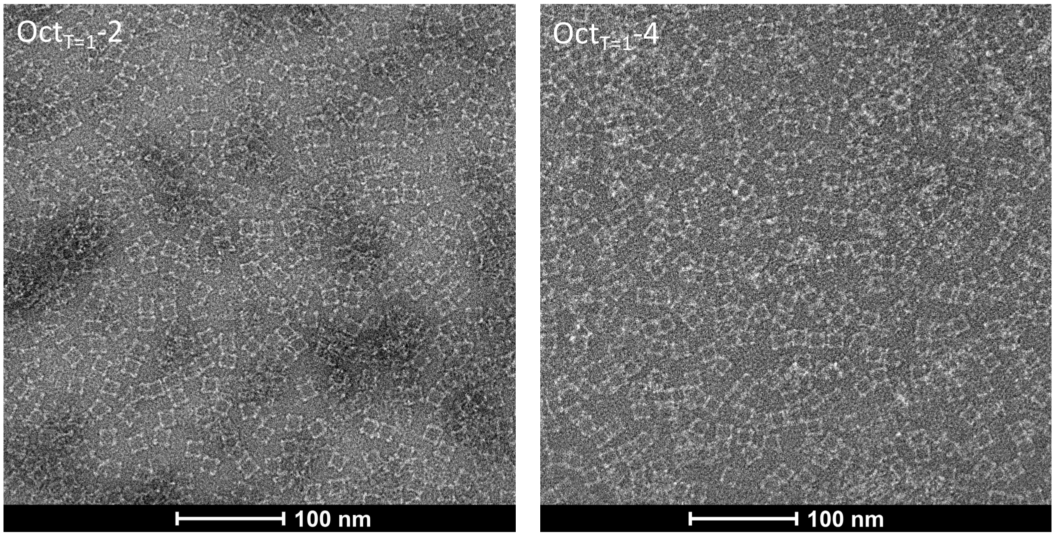


**Fig. S23.** nsEM images of octahedral cages (T=1, Symmetry: O3) (Oct$_{T=1}$-2 and Oct$_{T=1}$-4) obtained from SEC peaks shown in Fig. S22 (light grey box).

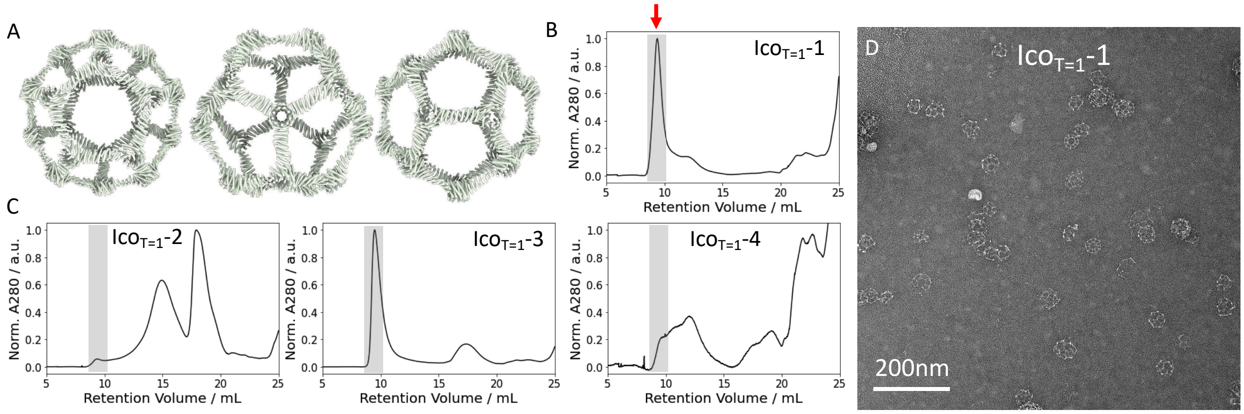


**Fig. S24**. (**A**) Structure model of T=1 icosahedral cage (symmetry: I3). (**B, C**) SEC traces of the cages ($Ico_{T=1}$-1– $Ico_{T=1}$-4) obtained from S6 column. Transparent grey box (~ 9ml) of each trace indicates an expected retention volume of the cage. (**D**) An nsEM image obtained from SEC peaks (red arrow) shown in (B).

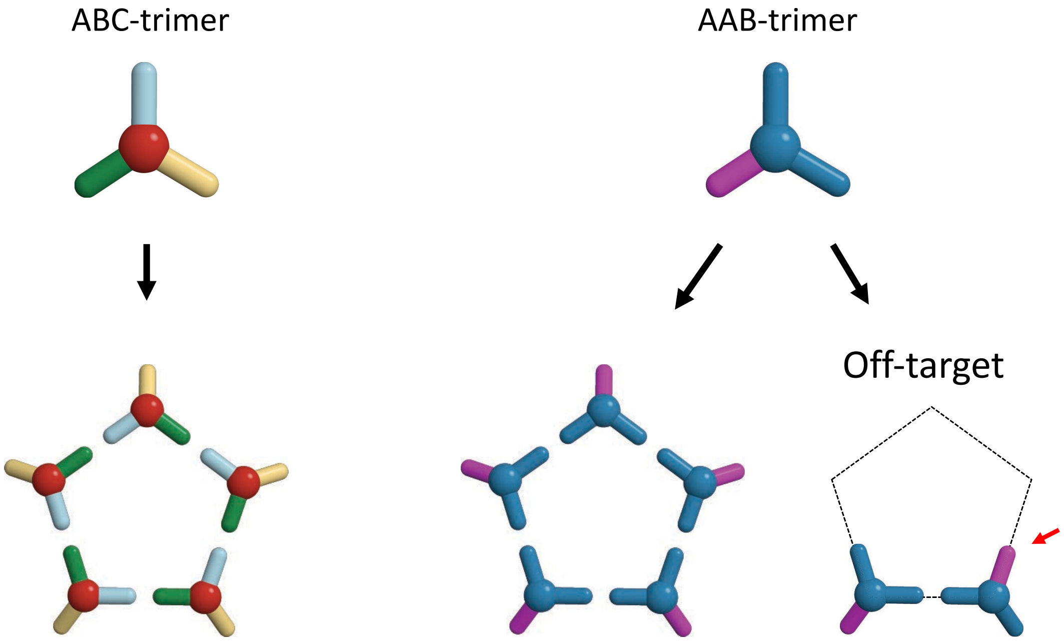


**Fig. S25.** Two types of heterotrimers (ABC-type and AAB-type) forming C5 crowns. If orthogonal interfaces are given, ABC heterotrimers can form C5 crowns exclusively without forming off-targets, while AAB-trimers can form off-targets (bottom right).

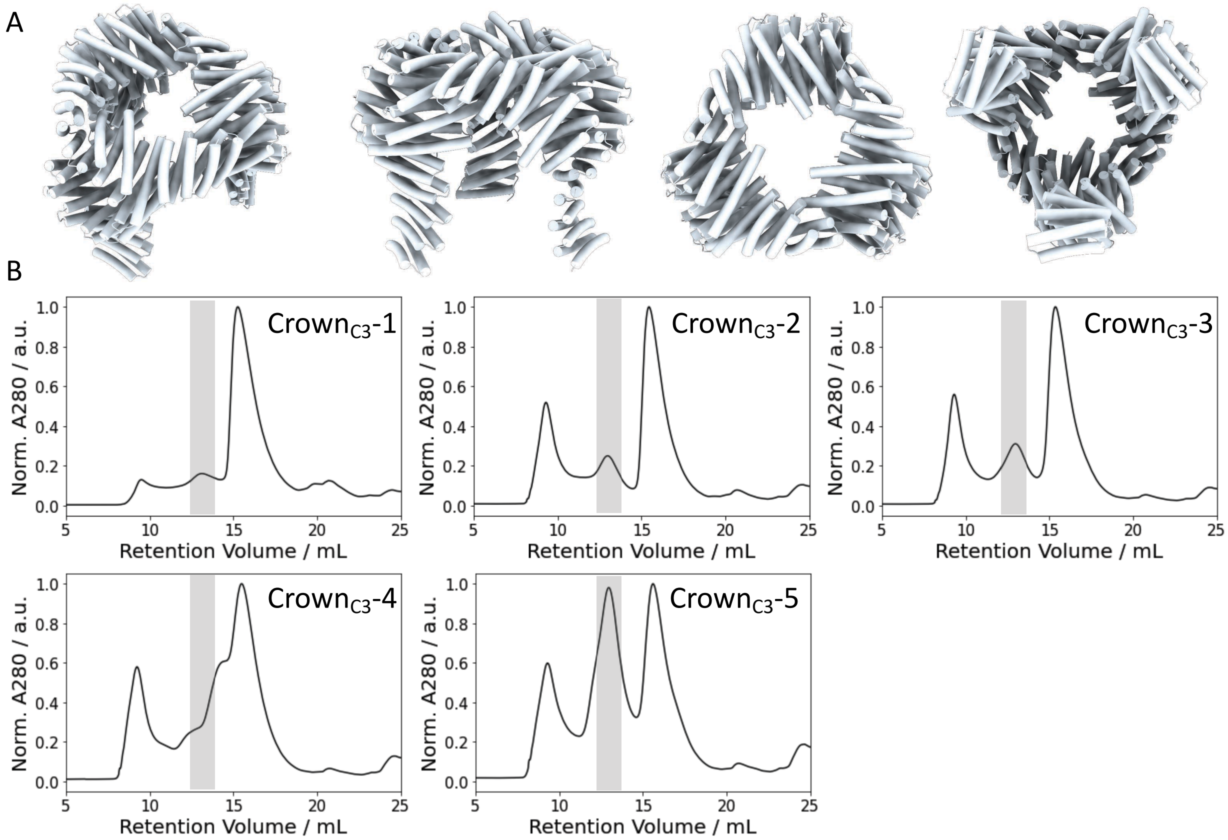


**Fig. S26.** (**A**) Structure model of C3 crowns. (**B**) SEC traces of the crowns (Crown$_{C3}$-1 – Crown$_{C3}$-5) obtained from S200 column. Transparent grey box (~ 13ml) of each trace indicates an expected retention volume of the crown.

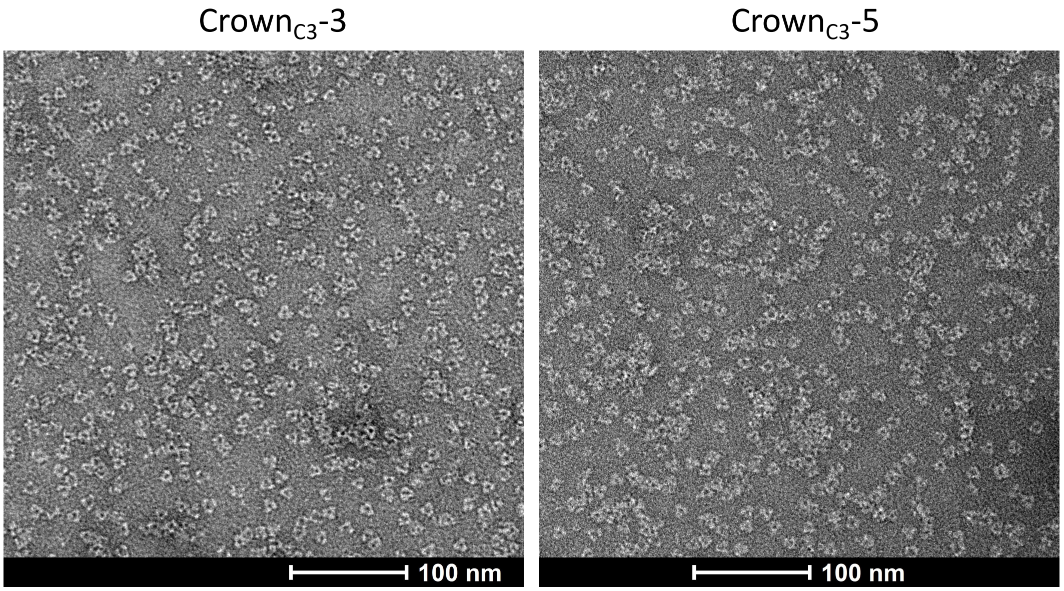


**Fig. S27.** nsEM images of C3 crowns (Crown$_{C3}$-3 and Crown$_{C3}$-5) obtained from SEC peaks shown in Fig. S26 (light grey box).

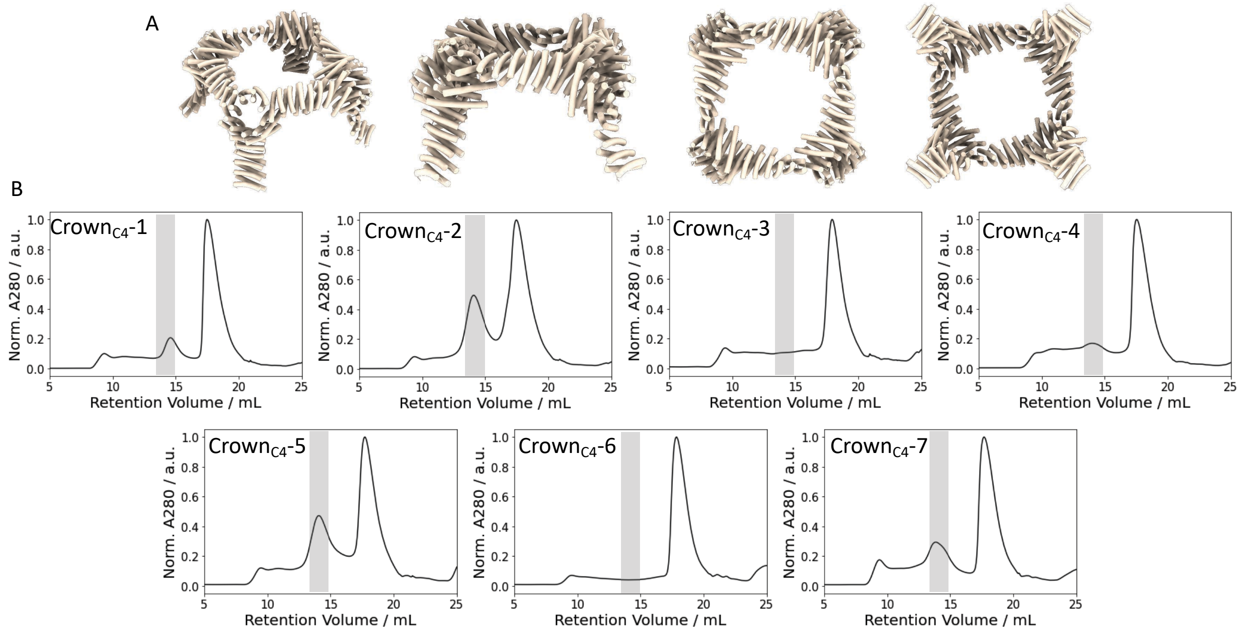


**Fig. S28.** (**A**) Structure model of C4 crowns. (**B**) SEC traces of the crowns (Crown$_{C4}$-1 – Crown$_{C4}$-7) obtained from S6 column. Transparent grey box (~ 14ml) of each trace indicates an expected retention volume of the crown.

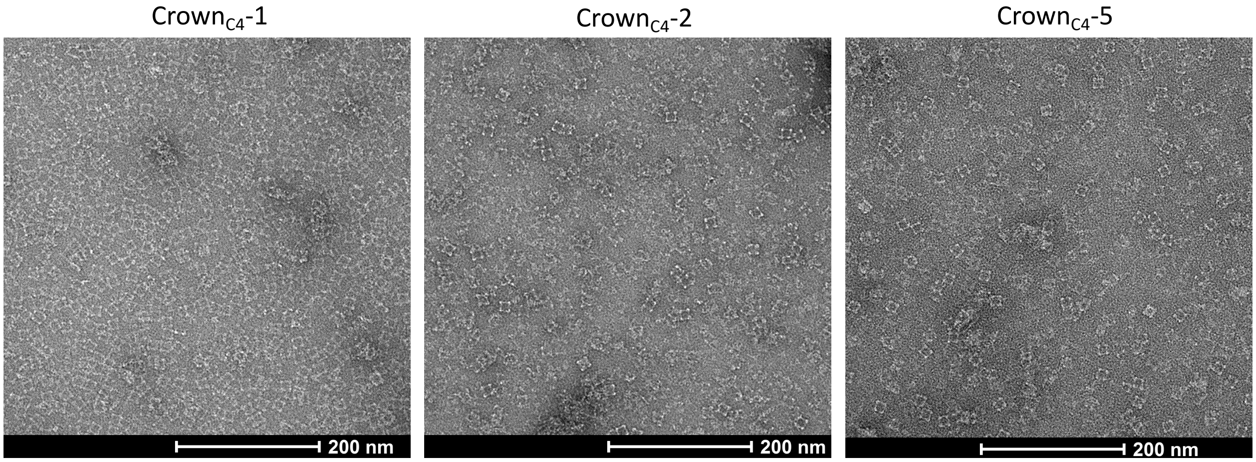


**Fig. S29.** nsEM images of C4 crowns (Crown$_{C4}$-1, Crown$_{C4}$-2, Crown$_{C4}$-5) obtained from SEC peaks shown in Fig. S28 (light grey box).

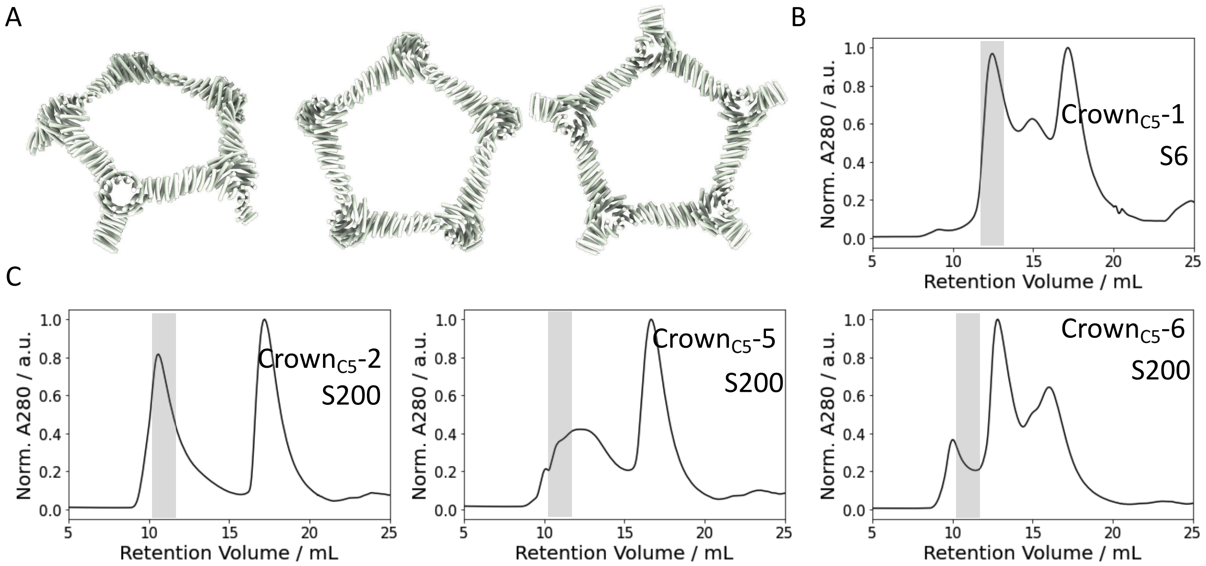


**Fig. S30.** (**A**) Structure model of C5 crowns. (**B, C**) SEC traces of the crowns (Crown$_{C5}$-1, Crown$_{C5}$-2, Crown$_{C5}$-5, Crown$_{C5}$-6) obtained from (**B**) S6 column and (**C**) S200 column. Transparent grey box of each trace indicates an expected retention volume of the crown.

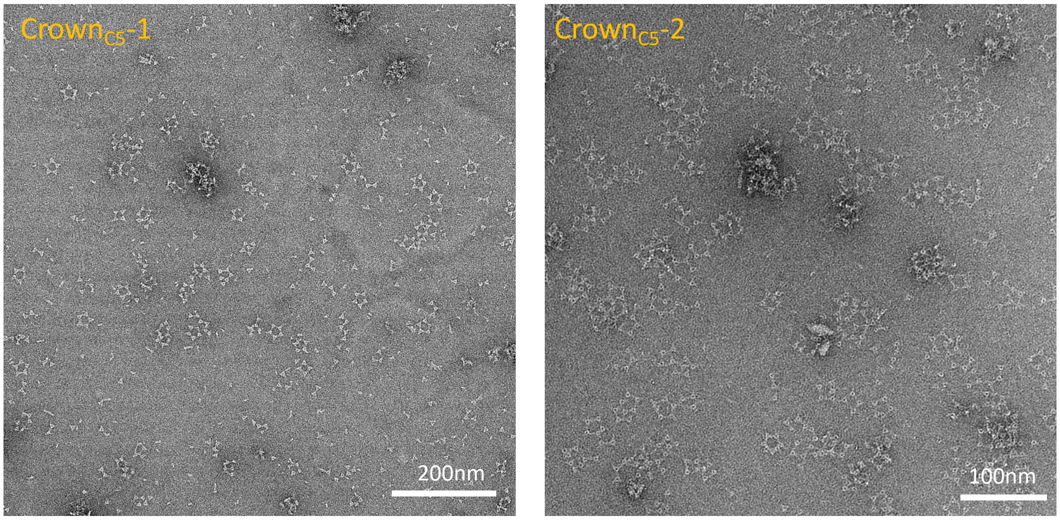


**Fig. S31.** nsEM images of C5 crowns (Crown$_{C5}$-1, Crown$_{C5}$-2) obtained from SEC peaks shown in Fig. S30 (light grey box).

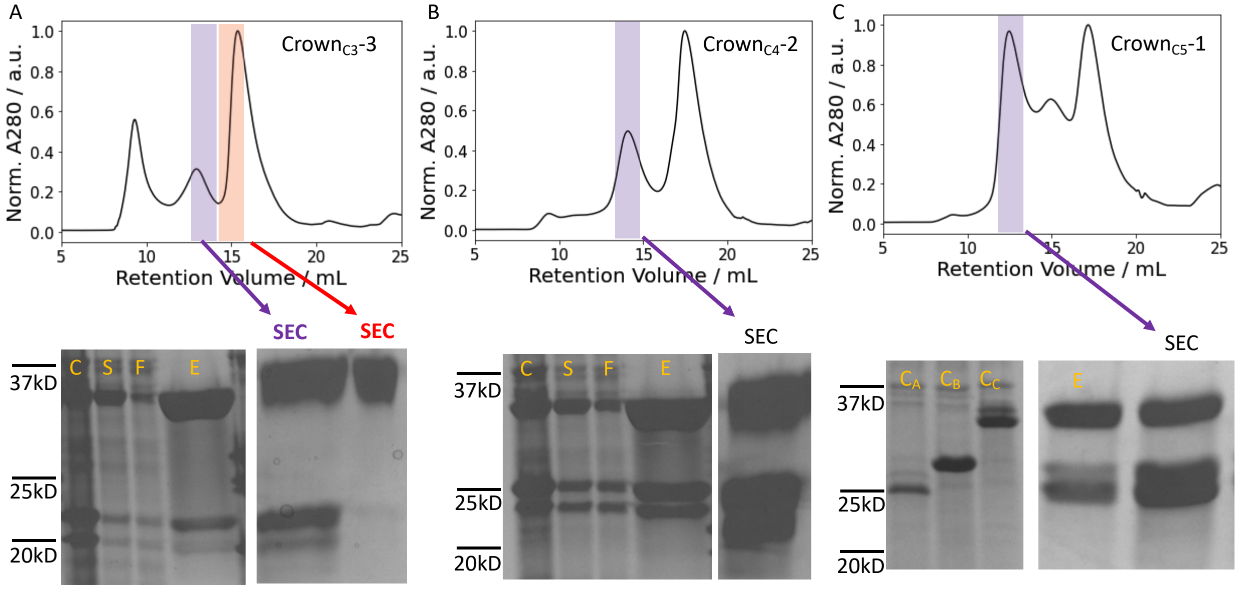


**Fig. S32.** (**A-C**) (top row) SEC trace and (bottom row) SDS-PAGE results of (**A**) C3 crown (Crown$_{C3}$-3), (**B**) C4 crown (Crown$_{C4}$-2), and (**C**) C5 crown (Crown$_{C5}$-1). In SDS-PAGE; C: E. Coli culture. S: Soluble fraction. F: Flow through from IMAC. E: Elution. SEC: SEC peak. $C_X$ indicates E. Coli culture of chain-X before mixing with other chains. For Crown$_{C3}$-3 and Crown$_{C4}$-2, a fraction of E. Coli culture was taken for the SDS-PAGE after mixing the three chains. For Crown$_{C5}$-1, a fraction of E. Coli culture of each chain was taken for the SDS-PAGE before mixing.

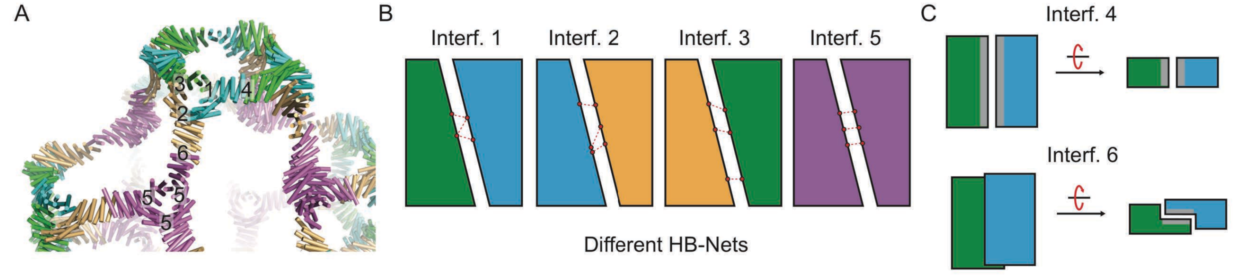


**Fig. S33.** Schematic diagram of six different interfaces of T=4 octahedral cage (Oct$_{T=4}$-3).

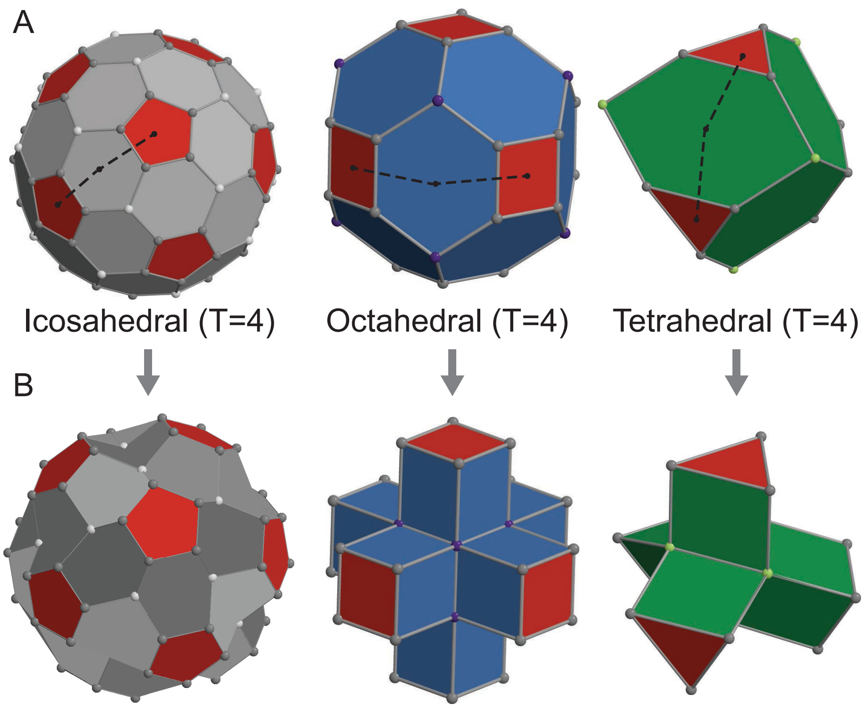


**Fig. S34.** T=4 polyhedra with (**A**) convex and (**B**) concave shapes.

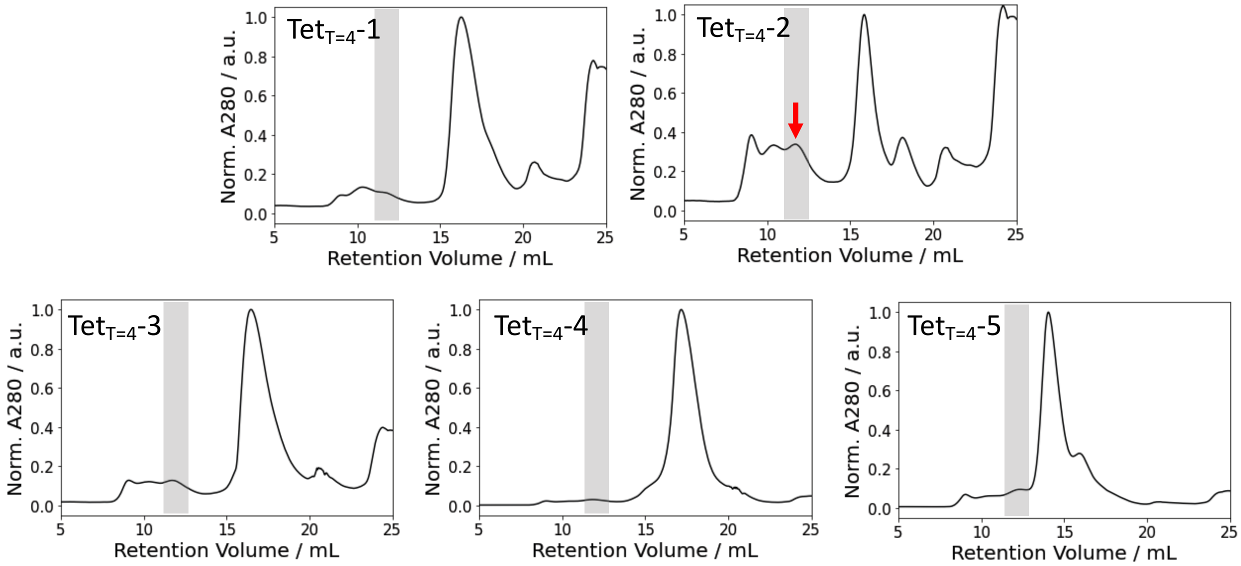


**Fig. S35.** SEC traces of the T=4 tetrahedral cages ($Tet_{T=4}$-1 – $Tet_{T=4}$-5) obtained from S6 column. Transparent grey box of each trace indicates an expected retention volume of the cage. A red arrow indicates the sample where the nsEM image of Fig. 4E was obtained.

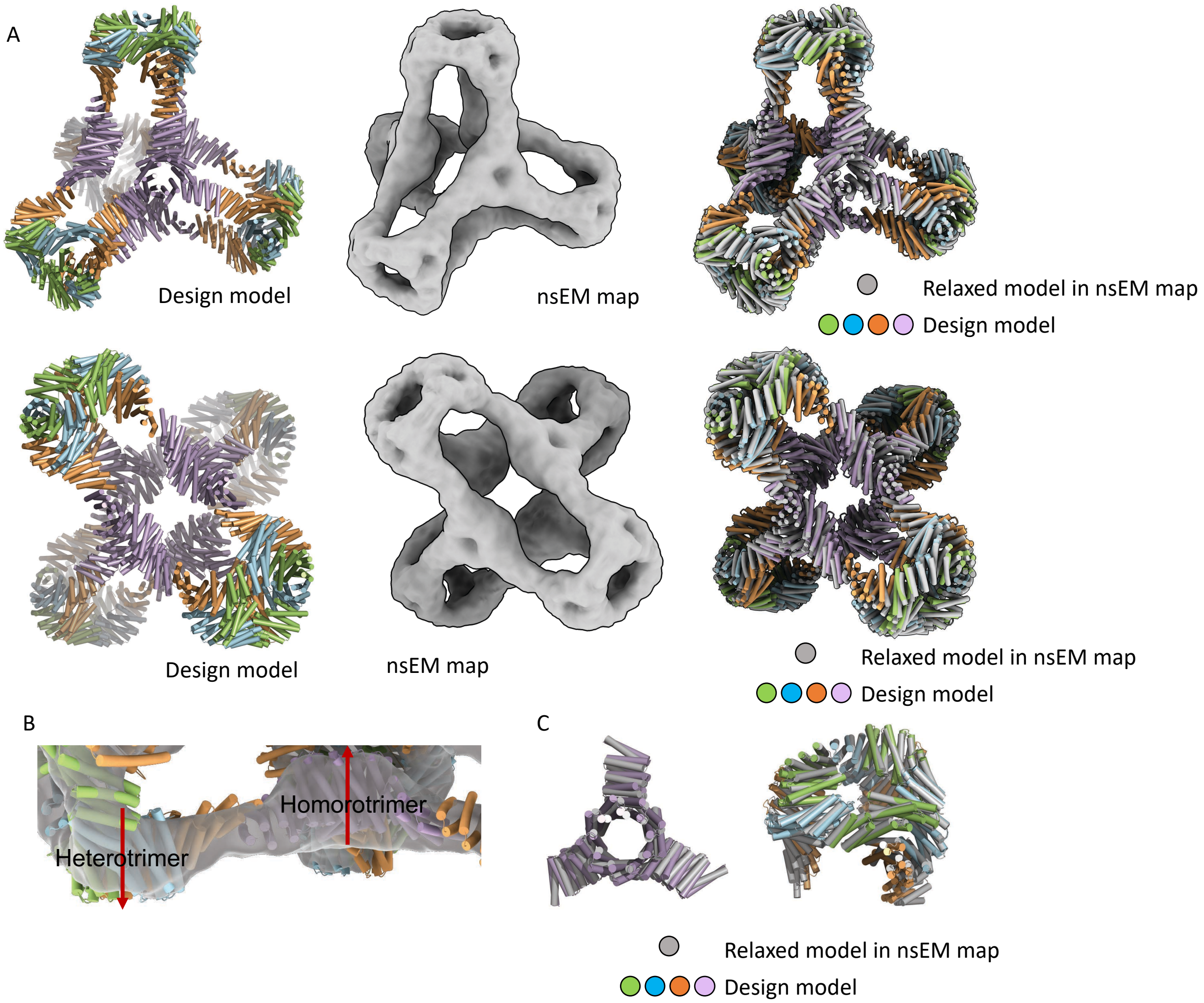


**Fig. S36.** (**A**) T=4 tetrahedra (Tet$_{T=4}$-2) (left) design model, (middle) 3D reconstruction from nsEM, and (right) overlay between a relaxed model in nsEM map (grey) and the design model (colors). Top and bottom rows were obtained from different viewing directions. (**B**) Overlay between nsEM map (grey transparent) and design model (colors). (**C**) Overlay between between a relaxed model in nsEM map (grey) and the design model (colors) for each substructures (left: homotrimer, right: C3 crown).

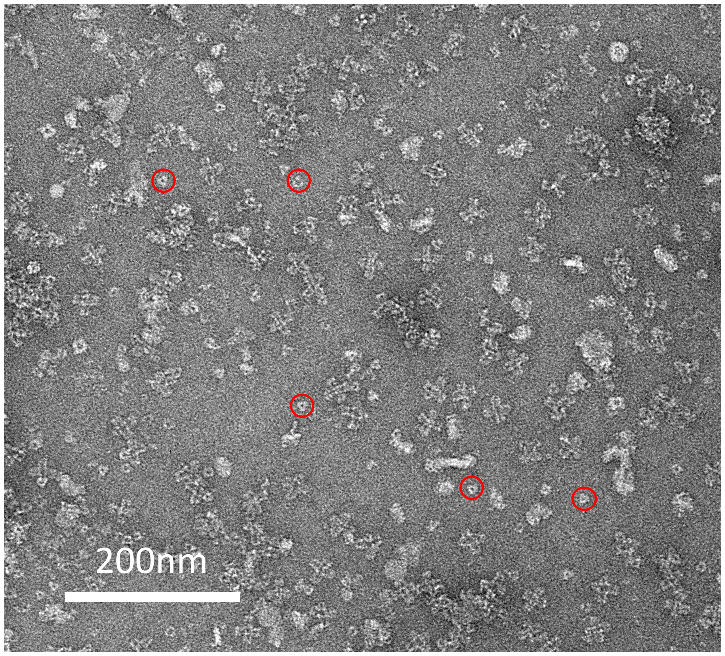

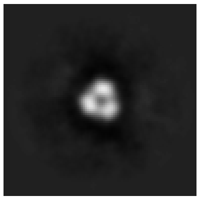

**Fig. S37.** Off-targets (red circles in the left micrograph) in T=4 tetrahedra system ($Tet_{T=4}$-2). (right) 2D class average of the off-targets.

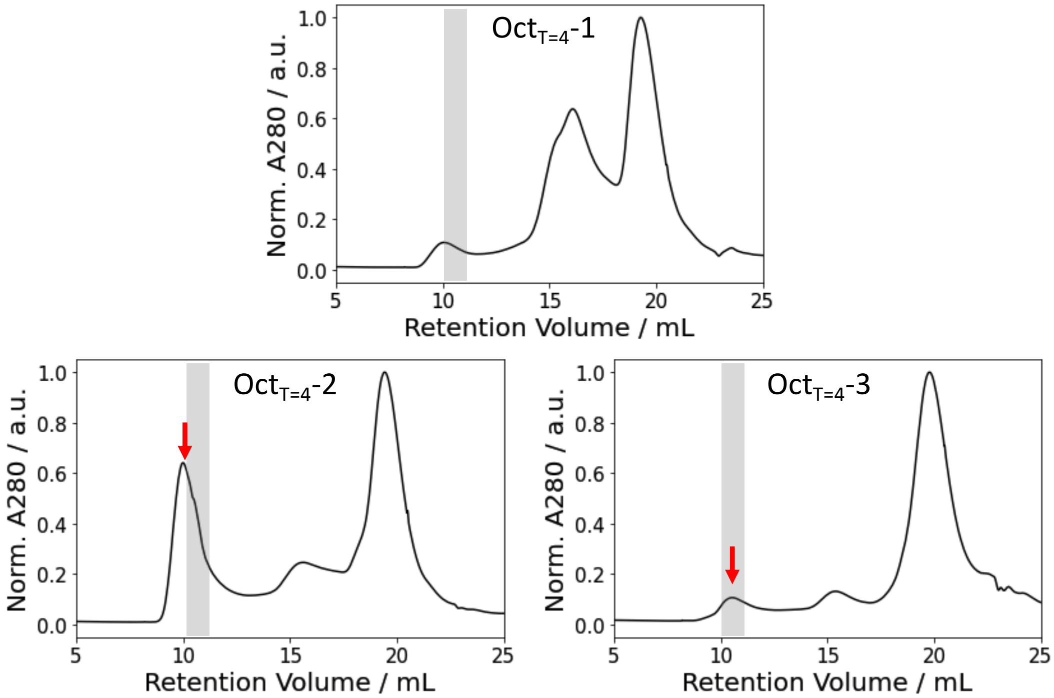


**Fig. S38.** SEC traces of the T=4 octahedral cages ($Oct_{T=4}$-1, $Oct_{T=4}$-2, $Oct_{T=4}$-3) obtained from S6 column. Transparent grey box of each trace indicates an expected retention volume of the cage. Red arrows indicate the peak where the nsEM images of Fig. 4I and Fig. S39 was obtained.

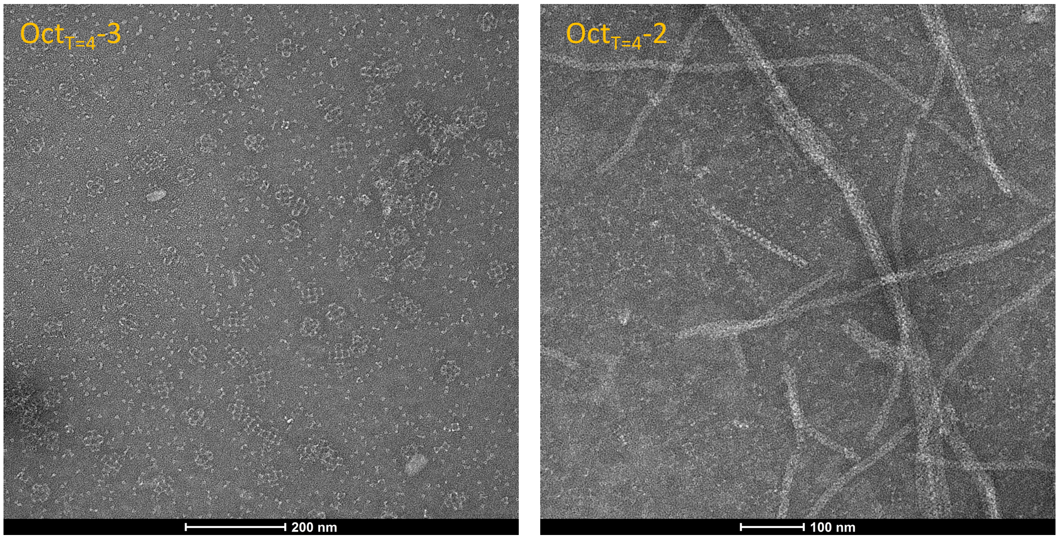


**Fig. S39.** nsEM images of T=4 octahedral cages (Oct$_{T=4}$-3 - left, Oct$_{T=4}$-2 - right) obtained from SEC peaks shown in Fig. S38 (red arrows). Right figure shows off-target formation.

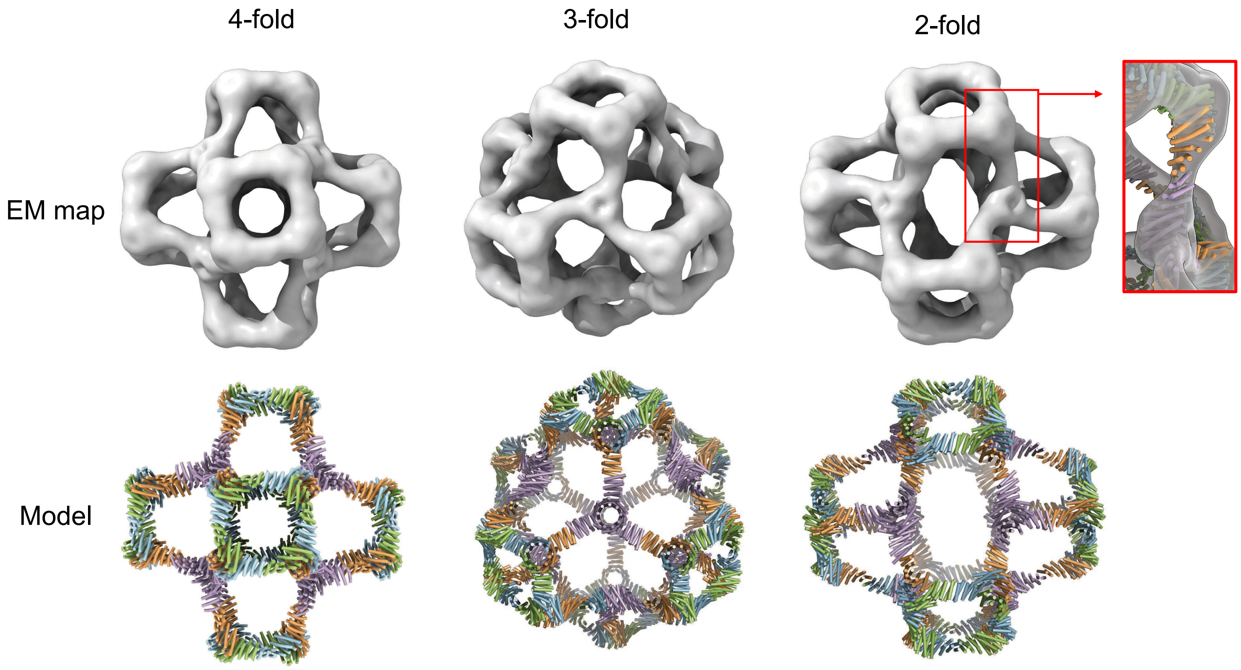


**Fig. S40.** T=4 octahedra (Oct$_{T=4}$-3) (bottom) model and (top) 3D reconstructions from nsEM, viewed from different characteristic symmetry axes: (left) 4-fold, (middle) 3-fold, and (right) 2-fold. The inset on the right is the overlay between nsEM map and the design model around the homotrimer-heterotrimer interface.

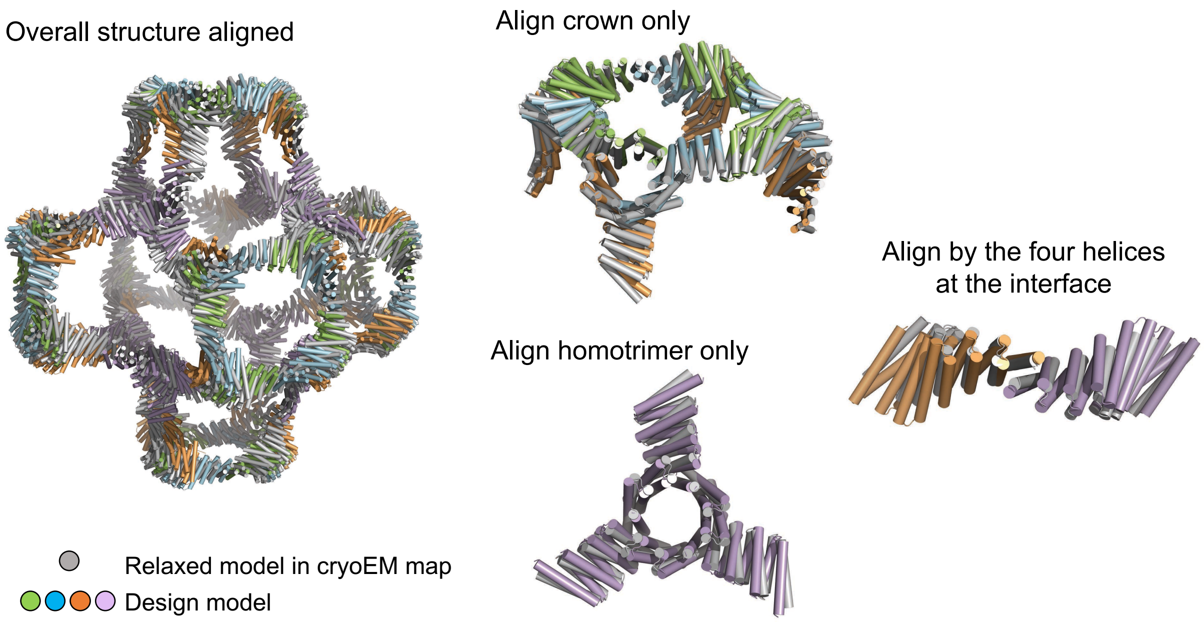


**Fig. S41.** Overlay between the relaxed model structure in cryoEM map (grey) and the design model structure (colors) of T=4 octahedra ($Oct_{T=4}$-3).

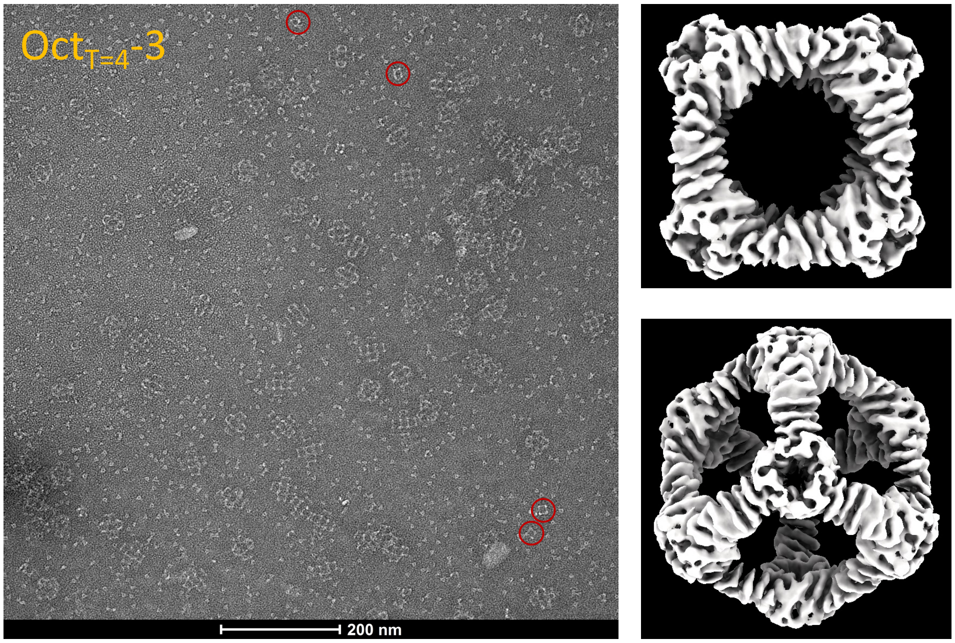


**Fig. S42.** O3 off-targets of T=4 octahedral cage system ($Oct_{T=4}$-3), identified in (left) a nsEM micrograph (red circles) and (right) 3D reconstruction of cryo-EM map.

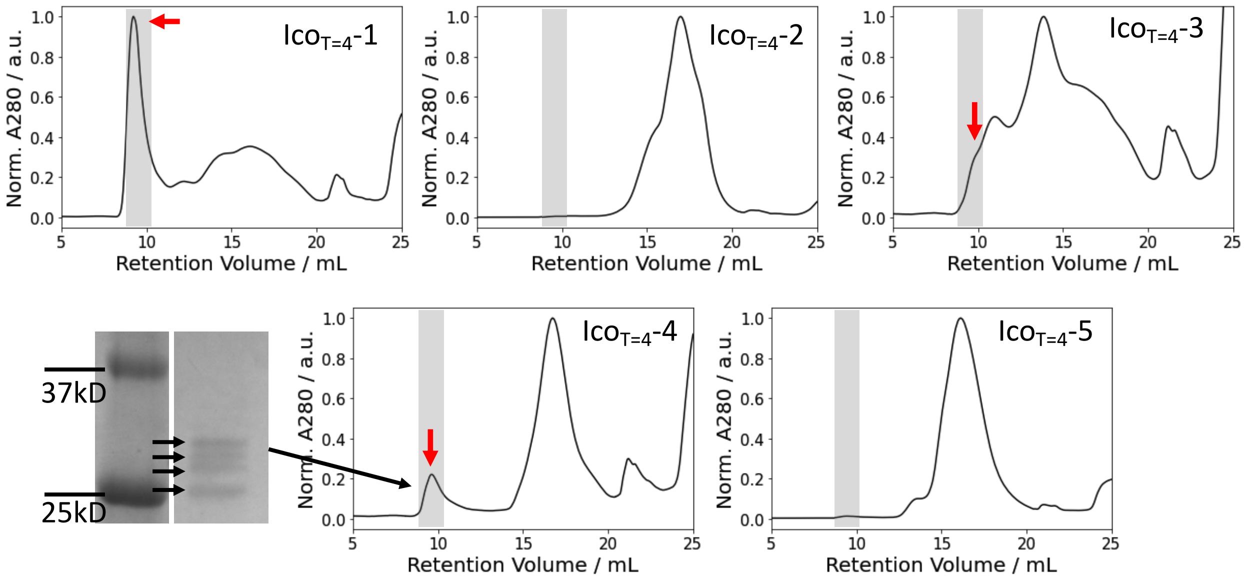


**Fig. S43.** SEC traces of the T=4 icosahedral cages (Ico$_{T=4}$-1– Ico$_{T=4}$-5) obtained from S6 column and SDS-PAGE result of a peak of Ico$_{T=4}$-4 system. Transparent grey box of each trace indicates an expected retention volume of the cage. Red arrows indicate the peak where the nsEM images of Fig. 4M and Fig. S44-S46 was obtained.

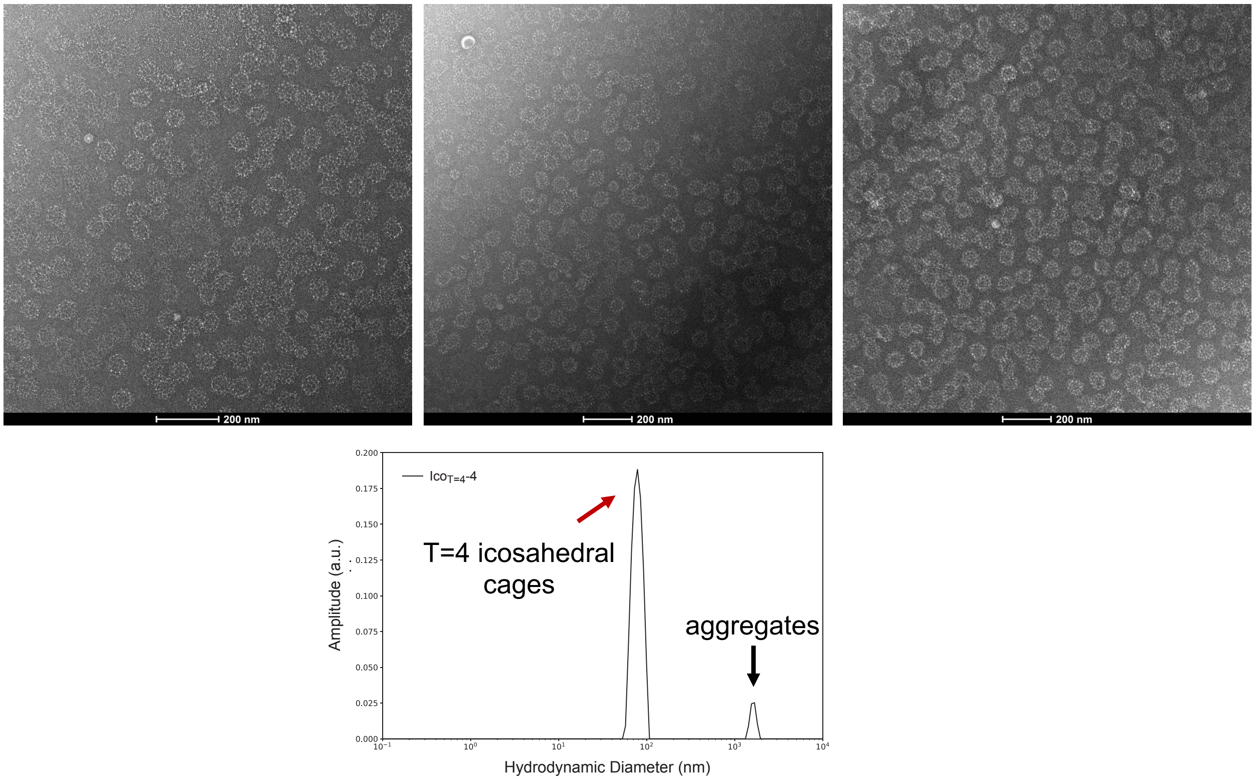


**Fig. S44.** Homogenous Ico$_{T=4}$-4 cages identified by (top row) nsEM micrographs from diverse stain thicknesses and (bottom row) dynamic light scattering (DLS) obtained at 25 $^{o}$C. Hydrodynamic diameter measured by DLS is 83.09 nm and the polydispersity index is 0.149.

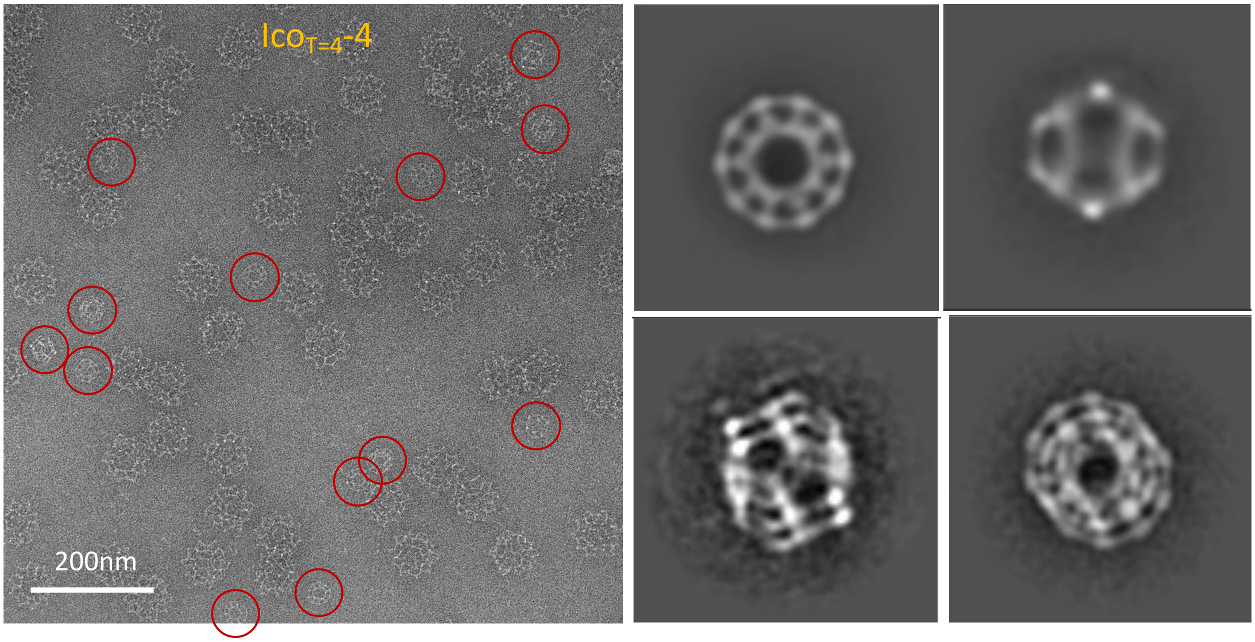


**Fig. S45.** I3 off-target cages identified from T=4 icosahedral cage sample. (left) Red circles are off-target cages. (right four small panels) 2D class averages of the off-targets. Top two are I3 cages and bottom two are entangled cages.

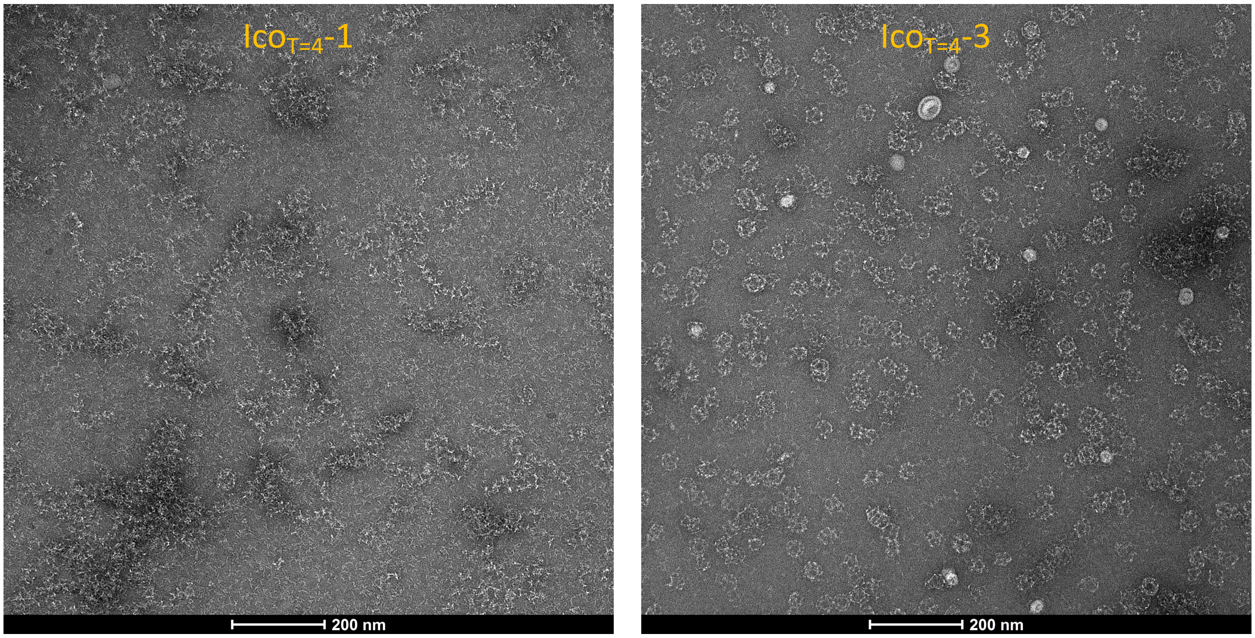


**Fig. S46.** nsEM images of (left) Ico$_{T=4}$-1 and (right) Ico$_{T=4}$-3 obtained from SEC peaks shown in Fig. S43 (red arrows).

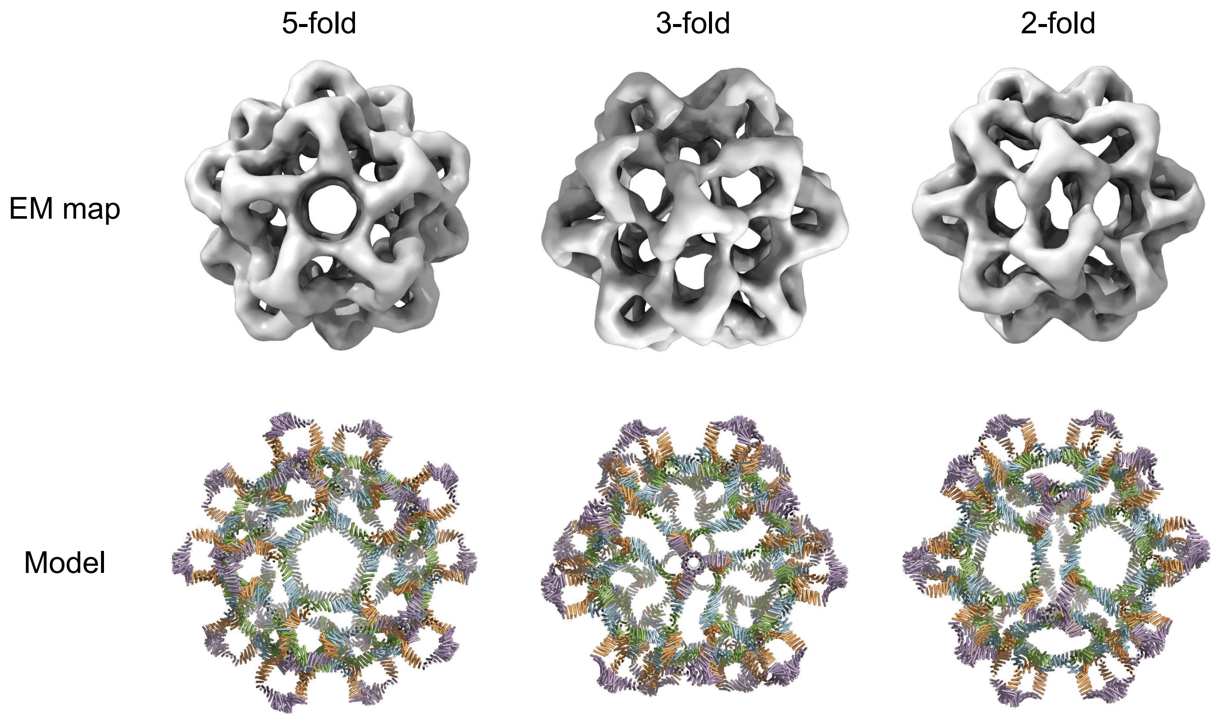


**Fig. S47.** T=4 icosahedra (Ico$_{T=4}$-4) (bottom) model and (top) 3D reconstructions from nsEM, viewed from different characteristic symmetry axes: (left) 5-fold, (middle) 3-fold, and (right) 2-fold.

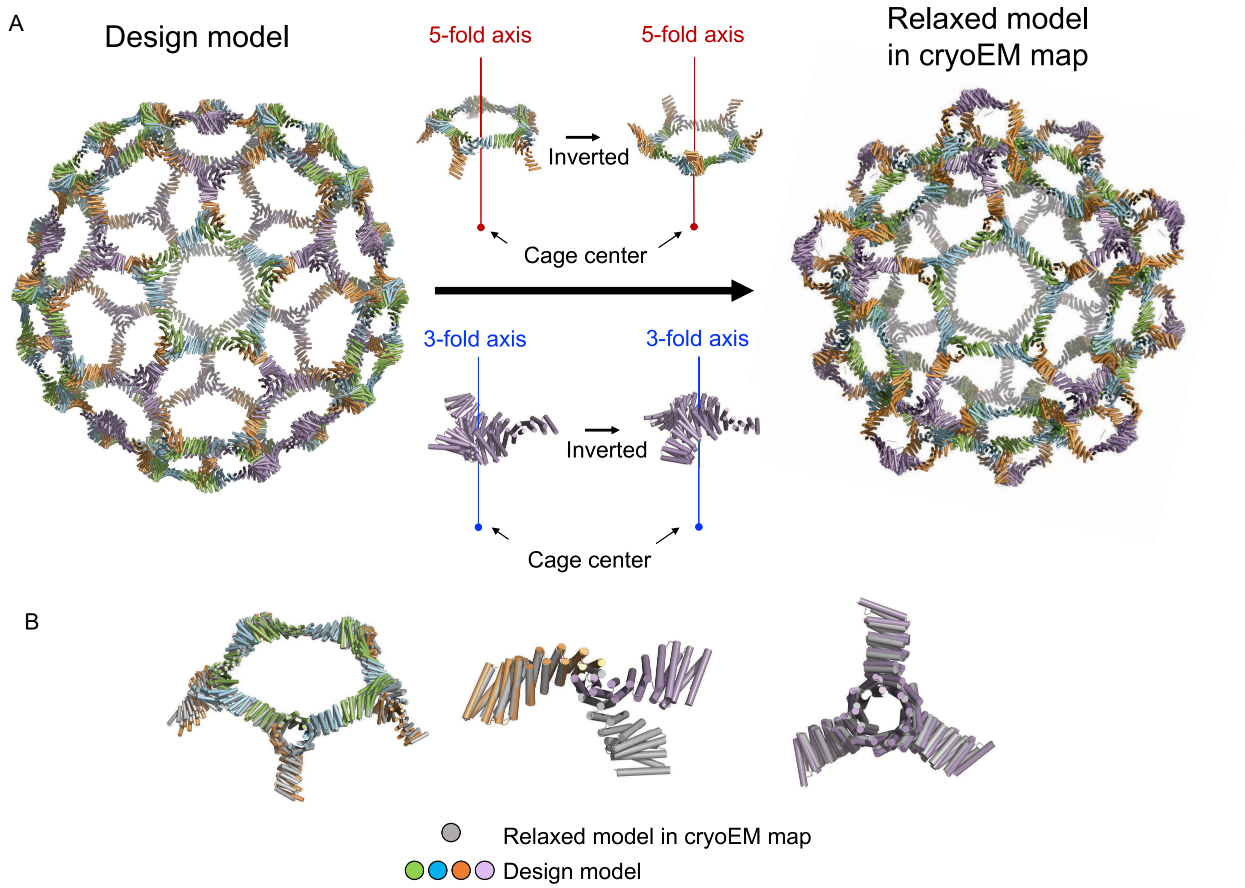


**Fig. S48.** (**A**) Structure difference between the design model (the model structure used when designing C5-C3 interfaces) and the relaxed model in cryoEM 3D map (shown in Fig. 5H). (**B**) Overlay between the design model (colors) and the relaxed model (grey) for each substructure. (left) C5 crown, (middle) heterotrimer-homotrimer interface, and (right) homotrimer.

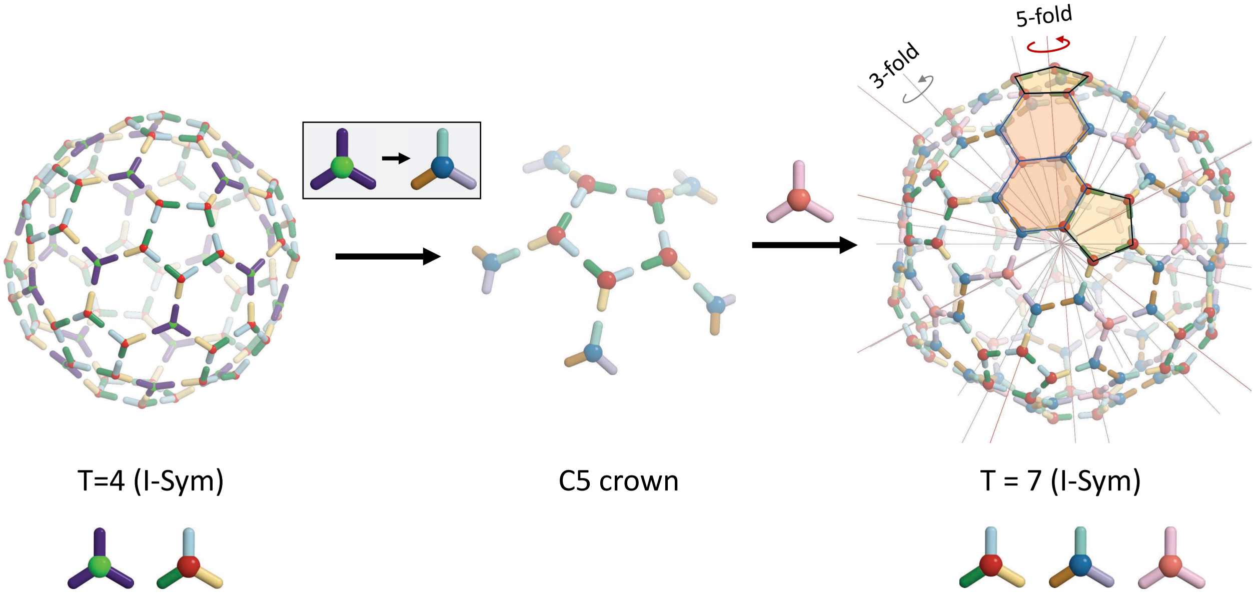

**Fig. S49**. Schematic diagram of design steps from T = 4 to T = 7 icosahedral cage (420 subunits). T = 7 cage requires two heterotrimers and one homotrimer.

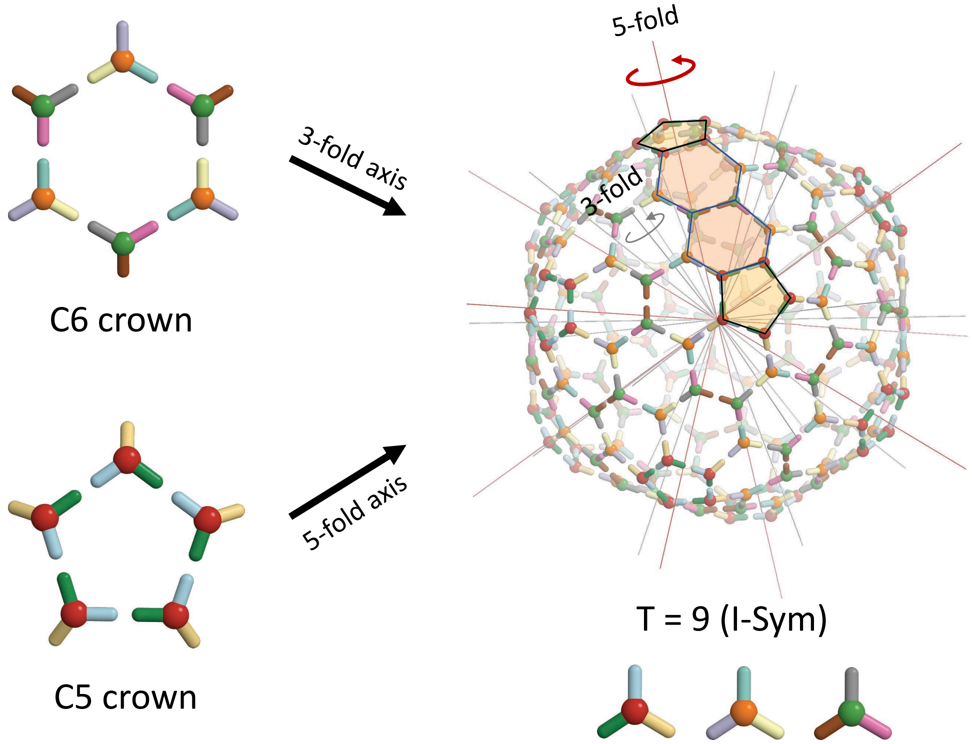


**Fig. S50**. Schematic diagram of T = 9 icosahedral cage (540 subunits) made by three heterotrimers.

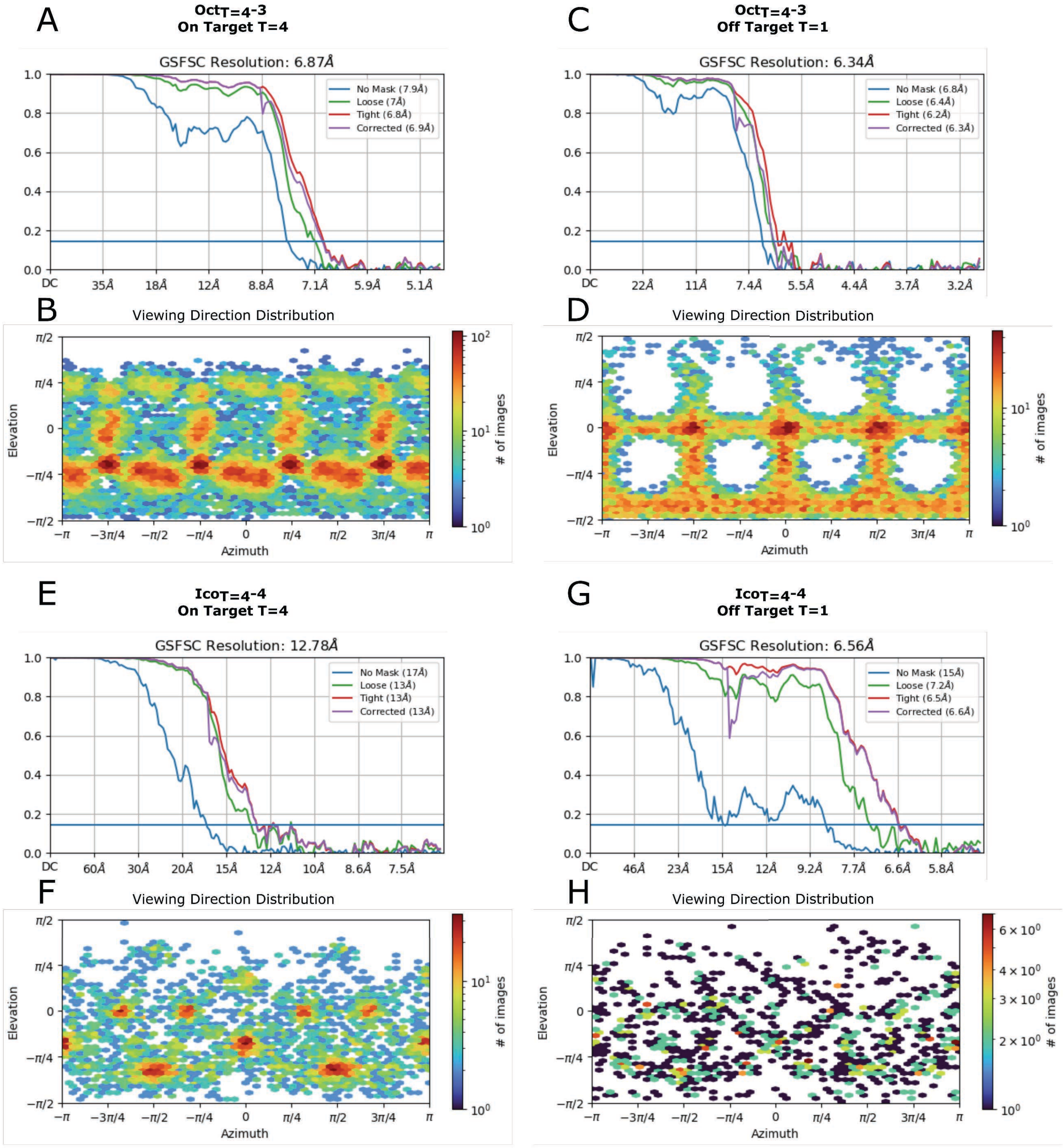


**Fig. S51.** CryoEM data processing plots for $Oct_{T=4}$-3 and $Ico_{T=4}$-4. (**A**, **C**, **E**, **G**) Gold Standard Fourier Shell Correlation curves. (**B**, **D**, **F**, **H**) Viewing angle direction distributions.

### Section 4. Experimental methods and materials

The experimental methods and materials used in this study are mostly same methods and materials described in our previous study (*5*).

4.1. Buffer and media recipes
All buffers and media were made using Milli-Q filtered water.

Autoinduction media (TBM-5052):
1.2% [wt/vol] tryptone, 2.4% [wt/vol] yeast extract, 0.5% [wt/vol] glycerol, 0.05% [wt/vol] D--glucose, 0.2% [wt/vol] D--lactose, 25 mM Na2HPO4, 25 mM KH2PO4, 50 mM NH4Cl, 5 mM Na2SO4, 2 mM MgSO4, 10 µM FeCl3, 4 µM CaCl2, 2 µM MnCl2, 2 µM ZnSO4, 400 nM CoCl2, 400 nM NiCl2, 400 nM CuCl2, 400 nM Na2MoO4, 400 nM Na2SeO3, 400 nM H3BO3

Lysis buffer:
25 mM Tris, 300 mM NaCl, 20 mM imidazole, 10% glycerol, pH 8.0 at room temperature

Wash buffer:
25 mM Tris, 300 mM NaCl, 40 mM imidazole, 10% glycerol, pH 8.0 at room temperature

Elution buffer:
25 mM Tris, 300 mM NaCl, 300 mM imidazole, 100 mM EDTA, 10% glycerol, pH 8.0 at room temperature

Het Lysis buffer:
25 mM Tris, 300 mM NaCl, 10 mM imidazole, 10% glycerol, pH 8.0 at room temperature

Het Wash buffer:
25 mM Tris, 300 mM NaCl, 20 mM imidazole, 10% glycerol, pH 8.0 at room temperature

Het Elution buffer:
25 mM Tris, 300 mM NaCl, 50 mM imidazole, 10% glycerol, pH 8.0 at room temperature

SEC buffer:
25 mM Tris, 300 mM NaCl, pH 8.0 at room temperature

SAXS buffer:
25 mM Tris, 300 mM NaCl, 2% glycerols, pH 8.0 at room temperature

TEV buffer:
25 mM Tris, 100 mM NaCl, 0.5 mM EDTA, 1mM DTT, pH 8.0 at room temperature

4.2. Construction of synthetic genes
All synthetic genes were ordered from either Integrated DNA Technologies Inc. (Coralville, IA, USA) (IDT) using pET29b+ vectors with kanamycin resistance, and genes were reverse translated and codon optimized using Domesticator (https://github.com/rdkibler/domesticator_3).

See Table S3-S14 for full protein sequence. Where indicated, hexa-histidine tag (histag) was added for purification, and a tobacco etch virus protease (TEVp) cleavage site (ENLYFQG) was added for the cases that the histag needs to be removed, such as crystallization, SAXS.

4.3. Protein expression

Plasmids (100ng) were transformed into chemically competent *E. coli* expression strain, BL21(DE3) or BL21(DE3)Star, for protein expression following manufacturer's protocol, with the exception of using 10ul competent cells per reaction. Following transformation and recovery, the entire transformation products were used to inoculate 1 mL Luria-Bertani (LB) medium containing 100 ug/mL kanamycin and grown at 37°C with shaking at 225 rpm overnight. 500ul of overnight cultures were diluted into 50 mL TBM-5052 supplemented with 100 ug/mL kanamycin in 250 mL baffled flasks and incubated at 37°C with shaking at 225 rpm for 18-24 hours.
For systems using hetero-oligomers (pseudosymmetric hetero-oligomers, crowns, and T=4 cages), we used two different protocols: *protocol 1* and *protocol 2*.

*Protocol 1*: Each component was transformed and incubated independently. After the incubation, we first checked relative expression level between components using SDS-PAGE (section 4.5) based on the intensity of bands. Then, the incubated E. coli cultures of each component were mixed with a correct stoichiometry for target assemblies (1:1:1 for hetero-trimers and crowns. 1:1:1:1 for T=4 cages). After mixing the E. coli cultures, we harvested proteins following *Section 4.4*.

*Protocol 2*: Plasmids of all components were transformed into E. coli together. For example, in hetero-trimer expression (3 components – A, B, C), 300ng of plasmids (100ng of A + 100ng of B + 100ng of C) were transformed into 10ul E. coli cells and incubated as described in this section. Then, proteins were harvested following *Section 4.4*.

4.4. Protein purification

*Immobilized metal affinity chromatography (IMAC)*

Cultures were harvested by centrifugation at 4000 rcf for 10 minutes, culture supernatant decanted, and pellets resuspended to 30 mL in Lysis buffer. 300ul PMSF (100mM in 100% EtOH) is added immediately prior to sonication at 70% power for 5 minutes. "Lysate" fractions are saved, and then lysates were clarified by ultracentrifugation at 18,000 rcf at 12°C for at least 30 minutes and applied to 1.5 mL Ni-NTA resin (Qiagen) pre-equilibrated with Lysis buffer and packed into Econo-Pac columns (Bio-Rad) for gravity chromatography. The columns were washed twice with 15 mL Wash buffer and eluted with 10 mL Elution buffer. Hetero-oligomers were purified according to a similar procedure, except they used the "Het" variants of the the Lysis, Wash, and Elution buffers as this was found to improve the yield of single-his-tagged complexes over non-specific multiple-his-tagged complexes which can arise at high concentrations and likely dominate binding to the Ni-NTA resin. Samples prepared for crystallization were treated similarly, except 500 mL cultures were used and lysate was divided among six gravity columns.

*Size-exclusion chromatography (SEC)*

IMAC eluted samples were concentrated using 10k MWCO spin concentrators and were purified using a Superdex 200 10/300 increase (for oligomers) or Superdex 6 10/300 increase (for crowns and cages) columns (Cytiva) in SEC buffer using an ÄKTA pure system (Cytiva). SEC traces were also used to qualitatively determine homogeneity and quantitatively measure total yield by A280 absorbance integrated over the collected fractions using Unicorn (Cytiva).

Note that, for T=4 icosahedral cages ($Ico_{T=4}$-4), we kept the IMAC eluted sample on room temperature for 24 hours *before concentrating and running SEC*. The 24h equilibration was important to generate homogeneous samples, and we think that even longer equilibration time could be helpful to further increase the level of homogeneity. Given the complexity of this sample (four components, six different interfaces, 240 subunits), a relatively slow assembly kinetics seems reasonable, thus, giving an enough equilibration time (>24h) is critical to get homogeneous samples.

For all other systems, the equilibration time is not a critical issue, so we concentrated the elution sample right after IMAC and ran SEC within a few hours. But, giving enough equilibration time (>24h) may not harm the sample quality, rather possibly increase the homogeneity.

*TEVp cleavage*

Purification and mass tags were buffer exchanged into TEV buffer and cleaved with TEVp at a ratio of 1 mg TEV per 100 mg substrate for 24-72h at room temperature. After TEV cleavage, samples were exchanged into Lysis buffer and passed over a Ni-NTA gravity column and washed with 10ml lysis buffer. Flowthrough was collected, concentrated using 10k MWCO spin concentrators, and purified once again by SEC.

4.5. Sample analysis

*SDS-PAGE*

Samples were diluted 1:1 with 2x Laemmli Sample Buffer (Bio-Rad) without Beta-mercaptoethanol and 15ul were loaded onto AnykD™ Criterion™ TGX™ Precast Midi Protein Gels (Bio-Rad). Ladder was 10ul of Precision Plus Protein™ Kaleidoscope™ Prestained Protein Standards (Bio-Rad). Gels were run at 300V for 18 minutes, then stained using an eStain™ L1 Protein Staining System (Genscript). Stained gels were imaged using a Chemidoc XRS+ (Bio-Rad)

*Liquid chromatography mass spectrometry (LC-MS)*

To identify the molecular mass of each protein and thus verify sample identity and integrity, intact mass spectra was obtained via reverse-phase LC/MS on an Agilent G6230B TOF on an AdvanceBio RP-Desalting column, and subsequently deconvoluted by way of Bioconfirm using a total entropy algorithm.

*Native mass spectrometry (nMS)*

Oligomeric state of SEC-purified and LC-MS verified samples was analyzed by on-line buffer exchange MS in 200 mM ammonium acetate using a Vanquish ultra-high performance LC system coupled to a Q Exactive ultra-high mass range Orbitrap mass spectrometer (Thermo Fisher Scientific). A self-packed buffer exchange column was used (P6 polyacrylamide gel, BioRad). The recorded mass spectra were deconvolved with UniDec version 4.2+.

*Negative stain electron microscopy (nsEM)*

SEC purified samples were diluted to (0.005 mg/ml for oligomers, 0.05 – 0.1 mg/ml for crowns and cages) using SEC buffer immediately before application for 45s to glow discharged thick carbon film-coated 400 mesh copper grids (CF400-CU TH) (Electron Microscopy Sciences). Grids were then stained and dried immediately twice using 2% uranyl formate. Dried grids were screened on a 120 kV Talos L120C transmission electron microscope. The E. Pluribus Unum (EPU) (FEI Thermo Scientific) software was used for automated data collection. Data processing was carried out in CryoSPARC™ (Structura Biotechnology Inc).

*CryoEM Sample Preparation*

To prepare cryoEM sample grids, 3 µL of 0.5 - 1.0 mg/mL of protein in 25 mM Tris and 300 mM NaCl at pH 8.0 was applied to glow-discharged Quantifoil R 2/2 300 mesh copper grids overlaid with a thin layer of carbon. Vitrification was performed on a Mark IV Vitrobot with a wait time of either 5 or 7.5 seconds, a blot time of 0.5 seconds, and a blot force of either 0 or -1 before being immediately plunged frozen into liquid ethane. The sample grids were clipped following standard protocols before being loaded into the microscope for imaging.

*CryoEM Data Collection*

Data collection was performed automatically using either Leginon (*9*) or SerialEM to control either a ThermoFisher Titan Krios 300 kV TEM equipped with a standalone K3 Summit direct electron detector with an energy filter or a ThermoFisher Glacios 200 kV equipped with a standalone K3 Summit direct electron detector (*10*). Both samples were collected using counting mode, with random defocus ranges spanning between -0.7 and -1.8 µm using image shift, with one shot per hole on the Glacios for $Oct_{T=4}$-3 or multiple shots per hole on the Titan Krios for $Ico_{T=4}$-4. 1,320 and 1,048 movies were collected with a pixel size of 0.44 Å for $Oct_{T=4}$-3 with a total dose of ~50 $e^{-}/Å^{2}$, and 6,678 movies were collected with a pixel size of 0.84 Å for $Ico_{T=4}$-4 with a total dose of ~61 $e^{-}/Å^{2}$.

*CryoEM Data Processing*

All data processing was carried out in CryoSPARC and CryoSPARC Live (*11*). Details for each system are below:

[Data processing details for $Ico_{T=4}$-4]

Alignment of movie frames was performed using Patch Motion with an estimated B-factor of 500 $Å^{2}$, with an output F-crop factor of 0.5. Defocus and astigmatism values were estimated using Patch CTF with default parameters. An initial set of 75 particles were picked manually and extracted with a box size of 1,280 pix binned to 420px. These were sorted into 10 classes using 2D Classification without forcing max poses/shifts. One class showing the 5-fold axis was subsequently used as the input for Template Picking, which produced 35,042 particles. These particles were extracted with a larger box size of 1428 px to better encompass the particle and binned to 360px. 50 classes were created by 2D classification and two classes representing recognizable particles (4,912) were used to initiate 3 3D classes using Ab-Initio reconstruction with icosahedral symmetry. Template picking was evidently failing to pick several good particles, so an additional set of 12,780 particles were picked using Manual Picker and extracted to a box size of 1,428px binned to 360px, resulting in 9,692 extracted particles, including particles of a contaminating T=1 species. All particles were sorted into 50 classes by 2D Classification and separated into classes containing the larger T=4 particles and the smaller T=1 particles.

The T=4 particles had a slightly preferred orientation favoring angles with the 5-fold axis centered, though particles with other orientations were observed during manual picking. Homogenous refinement with per-particle defocus optimization and enforced icosahedral symmetry was performed using hand-picked particle set and the prior 3D reconstruction of the T=4 species as input. Resolution was improved from 15.7 Å to 13.2 Å by non-uniform refinement, which used icosahedral symmetry, initial lowpass resolution of 20 Å, GSFSC split resolution of 8 A, large batch sizes (epsilon = 0.0001 and snr factor 70), per-particle defocus optimization, and a tight dynamic mask (near = 3Å, far = 5Å, starting at 15Å resolution). A final Local Refinement with icosahedral symmetry using pose/shift gaussian priors and rotation and shift re-centering after every iteration was used to improve the map resolution to 12.8 Å, keeping the output of this job as the final sharpened map. UCSF ChimeraX (*13*) was used to visualize the map. Final maps were deposited to the EMDB under accession number EMD-40260 (Fig. S51E-F). Ab initio reconstruction from the hand-picked particle set, heterogenous refinement, and subsequent non-uniform refinement of the largest class without enforcing symmetry at any step was performed to validate the use of icosahedral symmetry above, and the resulting maps superimpose well, with slight anisotropy likely due to poor angular particle distribution (data not shown).

The T=1 particles (1,172) displayed 2-fold and 5-fold views. These were re-extracted with a box size of 1096 px binned to 360 px and used to perform Ab-initio Reconstruction with icosahedral symmetry. The largest class (1,093 particles) was refined using Non-uniform refinement with icosahedral symmetry, per-particle defocus optimization, and tight dynamic masks (near = 3Å, far = 6Å). The final map achieved an estimated resolution of 6.6 A. It was sharpened using DeepEMhancer (*12*) and visualized using UCSF ChimeraX (*13*). Final maps were deposited to the EMDB under accession number EMD-40267 (Fig. S51G-H).

[Data processing details for $Oct_{T=4}$-3]

For both groups of movies, alignment of movie frames was performed separately using Patch Motion with an estimated B-factor of 500 $Å^2$, with an output F-crop factor of 0.5. Defocus and astigmatism values were estimated using Patch CTF with default parameters. Two species were present in this dataset: the intended T=4 species and an off-target T=1 species.

An initial set of 385 particles were picked manually for the T=4 species and sorted into 15 classes using 2D Classification without forcing max poses/shifts. Ten classes containing views down the 4-fold, 2-fold, and 3-fold axes were subsequently used as the input for Template Picking on both groups of movies, which produced 35,828 and 21,472 particles, respectively. These particles were extracted with an 800 px box size and separately sorted into 30 classes. The 24,710 particles comprising 14 classes from the larger set and 4,359 particles comprising 2 classes from the smaller set were selected and combined for C1 Ab-initio 3D reconstruction using 3 classes. The map of the class that contained the octahedral particle was used as input for non-uniform refinement using all particles (29,069). During refinement, we optimized per-particle defocus and used a tight dynamic mask (3 Å near and 6 Å far). The final map reached 6.87 Å GSFSC resolution and was sharpened using DeepEMhancer with the tightTarget model (*12*). Final maps were deposited to the EMDB under accession number EMD-40268 (Fig. S51A-B).

An initial set of 117 particles were picked manually for the T=1 species and sorted into 15 classes using 2D Classification without forcing max poses/shifts. 8 classes containing views down the 4-fold and 2-fold axes were subsequently used as the input for Template Picking on only the larger set of 1,320 exposures, which produced 112,972 particles. These particles were extracted with an 800 px box size and sorted into 30 classes. Many particles were picks of sub-structures of the larger T=4 particle due to their visual similarity. These were selected against and the remaining 25,302 particles were sorted again into 50 classes. 8 classes representing views down the 4-fold and 2-fold axes were selected (15,852 particles) and used for C1 Ab-initio 3D reconstruction using 1 class. The particles were downsampled to 300 px by fourier cropping and then run through Non-uniform refinement. During refinement, we optimized per-particle defocus and used a tight dynamic mask (3 Å near and 6 Å far). The final map reached 6.87 Å GSFSC resolution and was sharpened using DeepEMhancer with the tightTarget model (*12*). Final maps were deposited to the EMDB under accession number EMD-40269 (Fig. S51C-D).

*Small angle X-ray scattering (SAXS)*

TEVp-cleaved (and optionally AEC purified) samples were re-purified by SEC in SAXS buffer and concentrated using thoroughly washed 10k MWCO small spin concentrators; the flowthrough of concentration was used as blanks for buffer subtraction. Scattering measurements were performed at the SIBYLS 12.3.1 beamline at the Advanced Light Source as part of the HT-SAXS program. The X-ray wavelength ($\lambda$) was 1.27 Å, and the sample--to--detector distance was 1.5 m, corresponding to a scattering vector q ($q = 4\pi \sin \theta/\lambda$, where $2\theta$ is the scattering angle) range of 0.01 to 0.3 $Å^{-1}$. A series of exposures, in equal sub-second time slices, were taken of each well: 0.3 second exposures for 10 seconds resulting in 32 frames per sample. For each sample, data was collected for two different concentrations to test for concentration -dependent effects; "low" concentration samples were ~2.5 mg/mL and "high" concentration samples were ~ 5- mg/mL. Data was processed using the SAXS FrameSlice online server (*14*). FoXS (*15*) was used to compare design models to experimental scattering profiles and calculate quality of fit ($\chi$) values. The SAXS Similarity online server was used to compute the similarities of scattering profiles to each other and calculate quality of fit ($\chi$) values.

*X-ray crystallographic*

Crystals of BGL17_A31 were grown using protein purified as described above and TEV cleaved. Protein samples dispensed in 1 uL drops at purification concentrations were mixed with equal volume of a crystallization solution and set in hanging drops with 100mM ammonium citrate tribasic pH 7.0 and 10% w/v polyethylene glycol 3350. Vapor phase equilibration of the resulting drops against a 1 mL reservoir of the same crystallization solution resulting in growth of crystals. The crystals were flash cooled in liquid nitrogen after transfer into a cryoprotective solution of well solution plus 20% ethylene glycol. Diffraction data were collected on a Pilatus areas detector at the Advanced Light Source (ALS) synchrotron facility at beamline 5.0.1. The resulting data set (Table S2) extend to 4.5 Å with one copy of the protein in the asymmetric unit. The greater homotrimer can be generated via application of a crystallographic symmetry axis.

Data was processed using program HKL2000 (*16*). The placement of subunits was determined using the molecular replacement algorithm in program PHENIX (*17*). Local rebuilding was performed using the program COOT (*18*), followed by refinement using the program PHENIX (*17*). There was a shift in the alpha helices from the design to the crystal structure that required a rigid body fit of two helical bundles at a time to build into the density. The final values for Rwork

/ Rfree were 0.301 / 0.344 with good geometry (Table S2). Ramachandran Distribution is (Favored%/Allowed%/Outlier% = 93.82 / 5.88 / 0.29).

**Table S2.** Crystallographic data of BGL17_A31 (PDB ID: 8FLX).

| | BGL17_A31 |
|---|---|
| **Data collection** | |
| Space group | I23 |
| Cell dimensions | |
| *a*, *b*, *c* (Å) | 153.47, 153.47, 153.47 |
| α, β, γ (°) | 90, 90, 90, |
| Resolution (Å) | 4.5 (4.5-4.66) |
| $R_{sym}$ or $R_{merge}$ | 0.087 (0.821) |
| $I / \sigma I$ | 30.25 (2) |
| Completeness (%) | 97.4 % (94.5 %) |
| Redundancy | 18.7 (13.4) |
| **Refinement** | |
| Resolution (Å) | 4.5 |
| No. reflections | 3609 |
| $R_{work}$ / $R_{free}$ | 0.301 / 0.344 |
| No. atoms | |
| Protein | 342 |
| Ligand/ion | 0 |
| Water | 0 |
| *B*-factors | |
| Protein | 170.9 |
| Ligand/ion | - |
| Water | - |
| R.m.s. deviations | |
| Bond lengths (Å) | 0.002 |
| Bond angles (°) | 0.50 |

**Table S3.** Designed sequences of C3 homotrimers (BGL09_A01– BGL19_A39). In fusion type, "AB": c-terminus of BGL to n-terminus of DHR, "BA": n-terminus of BGL to c-terminus of DHR.

| ID | BGL | Arm | Fusion type | Protomer sequence |
|---|---|---|---|---|
| BGL09_A01 | BGL09 | DHR24 | BA | GSHHHHHHGSGENLYFQGWSSEAEELARRAAKEAKELCKRSTDEELCKELKKLAELLKELAERYPDSEAAKLALKAALEAIELCKQSTDEELCEELVKLAQKLIELAKRYPDSEAAKLALKAALLAIELCKEADDEKLCEKLVEDAQKAIELAKRYPDSEEAKTRLRTLEAALQAAKLAFLIKKVVERAIKHARTEEEALRLALKALELLVKAAHIIARSAREAGAEAMLELAARLAEEAARQAEEIARKARYEGNLELALKALQILVNAAYVLAEIARDRGNEELL |

| | | | | |
|---|---|---|---|---|
| | | | | EKAKRLVREAYRQAKEILEQLQKEGNAELALIALEILLEVQRVESELIRER |
| BGL09_A02 | BGL09 | DHR57 | BA | GSHHHHHHGSGENLYFQGWSTEELKKVLERVRELSERAKESTDPEEALKIAKEVIELALKAVKEDPSTDALRAVLEAVRLAAEVAKRVTDPDKALKIAKLVIELALEAVKEDPSEDALKAVLDAIALAFKVARRVTDEDKARKLAKLMQELATEAIKANPSLLANAAQAMALAALIEVVVRRAISHARTEEEALRLALKALELLVKAAHIIAESAREAGAEAALQVAAELAEEAARQAEEIARKARYEGNLELALKALQILVNAAYVLAEIARDRGNEELLEKARRLVREAYRQAQEILEQLQKEGNAELALIALEILLEVQRVESELIRER |
| BGL09_A03 | BGL09 | DHR82 | BA | GSHHHHHHGSGENLYFQGWSDEEVQEAVERAEELREEAEELIKKARKTGDPELLRKALEALEEAVRAVEEAIKRNPDNDEAVETAVRLARELKKVAEELQERAKKTGDPELLKLALRALEVAVRAVALAIKSNPDNDEAVETAVRLARELKKVAEELQERAKKTGDPELLKLALRALEVAVEAVALAILSNPDNEEAVEAAKRLAEELDKVARLLEERAKETGDPELKELARKAAAVAAVAQAIALAALILKVVKRAIKHARTEEEALRLALKALELLVKAAHIIARSAREAGAEAALAVAAKLAEEAARQAEEIARKARYEGNLELALKALQILVNAAYVLAEIARDRGNEELLEKAKRLVREAYRQAKEILEQLQKEGNAELALIALEILLEVQRVESELIRES |
| BGL14_A04 | BGL14 | DHR24 | BA | GSHHHHHHGSGENLYFQGWSSEAEELARRAAKEAKELCKRSTDEELCKELKKLAELLKELAERYPDSEAAKLALKAALEAIELCKQSTDEELCEELVKLAQKLIELAKRYPDSEAAKAALKAALALIEACKKADDEEFCEKAVEIAKALINLARELPDDELAKLLIENIKLIMELLAEVRKNDPELLLSVLEVLVRSVHVIAEVAREAGGEGALQAAAALAEMAAKAAEEVAREARYRGNLELALKALQILVNAAYVLAEIARDRGNEELLQKAHELAREALRQVKEILEQARKEGNLELVIIALRLHTEIMRVLVEIWRHR |
| BGL14_A05 | BGL14 | DHR57 | BA | GSHHHHHHGSGENLYFQGWSTEELKKVLERVRELSERAKESTDPEEALKIAKEVIELALKAVKEDPSTDALRAVLEAVRLASEVAKRVTDPDKALKIAKLVIELAAEAVREDPSTDALRAVLEAVRLASEVAKRVTDPDKALKIAELVLSLAALAVAMNPSEEARRAVEEARRLAKEVAKRVTDPKLSLRLEALALLLENITLIAELLEEVKHNDPELLLSVLEVLVRSVHVIAEVAREAGVEALLEIAAILAELAARQAEEVAREARYRGNLELALKALQILVNAAYVLAEIARDRGNEELLQKADELAREALRQVKEILEQARKEGNLELVIIALRLHTEIMRVLVEIWRHS |
| BGL14_A06 | BGL14 | DHR82 | BA | GSHHHHHHGSGENLYFQGWSDEEVQEAVERAEELREEAEELIKKARKTGDPELLRKALEALEEAVRAVEEAIKRNPDNDEAVETAVRLARELKKVAEELQERAKKTGDPELLKLALRALEVAVRAVELAIKSNPDNDEAVETAVRLAQELVKVAALLAELALRTGDEELLKLAKRALEVAERAVELAIKSNPDNEEARRAAAIIRLAELLIENIKLIQELLEEVKHNDPELLLSVLEVLVRSVHVIAEVAREAGLEAFLEMAALLAEEAARQAEEVAREARYRGNLELALKALQILVNAAYVLAEIARDRGNEELLQKAHELAREALRQVKEILEQARKEGNLELVIIALRLHTEIMRVLVEIWRHR |
| BGL15_A07 | BGL15 | DHR57 | BA | GSHHHHHHGSGENLYFQGWSTEELKKVLERVRELSERAKESTDPEEALKIAKEVIELALKAVKEDPSTDALRAVLEAVRLASEVAKRVTDPDKALKIAKLVIELAAEAVREDPSTDALRAVLEAVRLASEVAKRVTDPDKAAKIATLVLSLAALAVAANPSEEAKRAVEEARRLAEEVAKRVTDPKKSARIKILVLVMELIAHTAELRELLEKLVKHGGASEEYLLELLENLVRLAHVIAEVAREAGVEMMLEIAALLAEDAARQAEELAREARYEGNLELALKALQILVNAAYVLAEIARDRGNEELLEKAERLAREALRQVREISKRLQKEGNIELALKANRLLIDALRVLVRIMRHG |
| BGL15_A08 | BGL15 | DHR62 | BA | GSHHHHHHGSGENLYFQGWSNDEKRKRAEKALQRAQEAEKKGDVEEAVRAAQEAVRAAKESGDNDVLRKVAEQALRIAKEALKQGNAEVAVKAARVAVEAAKQAGDNDVLRKVAEQALQIAAAALALGNADVAQKAIRVAKEAAKQAGDEKVKLKVKILELVVELYKHTKELRELLEKLVKHGG |

| | | | | |
|---|---|---|---|---|
| | | | | ASEEYLLELLENLVRLAHVMAEVAREAGLEALLEAAARLAEEAARQAEELAREARYEGNLELALKALQILVNAAYVLAEIARDRGNEELLEKAERLAREALRQVREISKRLQKEGNIELALKANRLLIDALRVLVRIMRHG |
| BGL15_A09 | BGL15 | DHR79 | BA | GSHHHHHHGSGENLYFQGWSMSSDEEEARELIERAKEAAERAQEAAERTGDPRVRELARELKRLAQEAAEEVKRDPSSSDVNEALKLIVEAIEAAVRALEAAERTGDPEVRELARELVRLAVEAAEEVQRNPSSSDVNEALKLIVEAIEAAVQALEAAEKSGDPEVRELARRLVKALVRAAEEVQRNPSSRKVNNILRAIALVVKLFVHTLELRLLLEKLVKHGGASEEYLLELLENLVRLAHVIAEVAREAGIEALLEVAAILAEDAARQAEELAREARYEGNLELALKALQILVNAAYVLAEIARDRGNEELLEKAERLAREALRQVEEISKRLQKEGNIELALKANRLLIDALRVLVRIMRHG |
| BGL00_A10 | BGL00 | DHR26 | AB | GSLELALKALQILVNAAYVLAEIARDRGNEELLRKAARLAEEAARAAEEIAREARKEGNLELALKALQILVNAAYVLAEIARDRGNEELLEYAARLAEEAARQAWEIAAEALERGNLELALKALQILVNAAYVLAEIARDRGNEELLEKAARLAELAALAAEVIAELARLKEEACRSNSDECLRLASEVEKAVAKLARLAAKATDEEVRRVALEVARELIKLAQEACRSNDDECLRLASEVVKAVQELVKLAEQATDEEVIRVALEVARELIRLAQEACRSNDEECLREASEVVKEVQELVKEAEKSTDEEEIRELLQRAEERIREAQERCREGDGWSENLYFQGSGHHHHHH |
| BGL00_A11 | BGL00 | DHR53 | AB | GSLELALKALQILVNAAYVLAEIARDRGNEELLRKAARLAEEAARLAEEIARQARKEGNLELALKALQILVNAAYVLAEIARDRGNEELLEYAARLALKAAVMALEIAAQALKEGNLELALKALQILVNAAYVLAEIARDRGNEELLRIAAILAEAAAAFALFIANPGSNEAKKAAELVLRLAEELAKSPDPEALKAAIAMAEAVVALNPGSNLAKKALEIILRAAEELAKLPDPEALKEAVKAAEKVVREQPGSELAKKALEIIERAAEELKKSPDPEAQKEAKKAEQKVREERPGGWSENLYFQGSGHHHHHH |
| BGL00_A12 | BGL00 | DHR79 | AB | GSLELALLALQILVNAAYVLAEIARDRGNEELLRKAARLAEEAARAAEEIARQARKEGNLELALKALQILVNAAYVLAEIARDRGNEELLEYAARLAEEAARQALEIARQALKEGNLELALKALQILVNAAYVLAEIARDRGNEELLEKAARLAEIAAIIAKAIAVAAEMEGVASKTGDPRVRELAREMTRLIAEAASEFLRDPSSSDVREALELIAKAAKAAARALRAAERTGDPEVRELARELVRLAVEAAEEVQRNPSSSDVNEALKLIVEAIEAAVRALEAAERTGDPEVRELARELVRLAVEAAEEVQRNPSSEEVNEALKKIVKAIQEAVESLREAEESGDPEKREKARERVREAVERAEEVQRDPGWSENLYFQGSGHHHHHH |
| BGL01_A13 | BGL01 | DHR49 | AB | GSPDLALKALRLLVKQVKSLAELARKRGDEKLLQEAAELAEEAAELAEKIAREARKKGNLELALQALQIMVEAAHVLAEIARERGNEELLEYAFRLAKEAFRQAKEILEQAVREGNLELALIALEILAEATHVLIEIAKEHNSEEEVRKIQEESIILSLIIILAVLADQSQDSEVLEEAIRQILRVAKKAGSEDALRLAILAVALIAREAQDSEVLEEAIRVILRIAKESGSEEALRAAIEAVAEIAKEAQDPRVLEEAIRVIRQIAEESGSEEARRQAERAEEEIRRRAQGWSENLYFQGSGHHHHHH |
| BGL01_A14 | BGL01 | DHR59 | AB | GSPDLALRALRLLVKQVKSLAELARKRGDEKLLQLAAELAEEAAELAEKIAREARKKGNLELALQALQIMVEAAHVLAEIARERGNEELLEYAFRLAKEAFRQAAEILRQAVEEGNLELALIALEILAEATHVLIEIAKEHNSEEEVRIIQLASIVMSLAILLAIELPSEEAQKVVTRIIEAAEAAIRAAEQGKTEVAELALKVLLEAIKLAEENRSEEALKVVLEIARAALAAAQAAEEGKTEVAKLALKVLEEAIELAKENRSEEALKVVLEIARAALAAAQAAEEGKSDEARDALRRLEEAIEEAKENRSKESLEKVREEAKEAEQQAEDAREGKGWSENLYFQGSGHHHHHH |
| BGL01_A15 | BGL01 | DHR82 | AB | GSPELALIALRLLVKQVKSLAELARKRGDEKLLQEAAELAEKAAELAEEIARLARKAGRLELALQALQIMVEAAHVLAEIARERGNEELLEYAFRLAKEAFRQAKEILEQAFEEGNLELALIALEILAEATHVLIEIAKEHNSEEE |

| | | | | |
|---|---|---|---|---|
| | | | | VRKIQEESILLSLIIILAVLARALIELARRLGDPELLRRALEALEEAVRVA<br>EEAIKKNPDAEFAVKAAVILARLLKEVAEELQERAKKTGDPELLKLAL<br>RALEVAVRAVELAIKSNPDNDEAVETAVRLARELKKVAEELQERAKK<br>TGDPELLKLALRALEVAVRAVELAIKSNPDNEEAVETAKRLAEELRKV<br>AELLEERAKETGDPELQELAKRAKEVADRARELAKKSNPNNGWSEN<br>LYFQGSGHHHHHH |
| BGL02_A16 | BGL02 | DHR62 | AB | GSPDLFLKALELLIKLAESLAETARERGDEKALEEAARIAEKAAELAEK<br>LARKARKEGNLELALQALRAMVEAARVLAEIARERGNEELLKYAWEL<br>AREAARQALEIAAEAAFRGNWELALIALEILVEALKVLSHIAREKGDPK<br>LLEESMLWQRILVALVEALLALLEGDVETAVRAAQEAVRIAKEAGLNE<br>ALRAVAMLAMAIAKAAEKQGNVEVAVKAARVAVEAAKQAGDNDVLR<br>KVAELALRIAKEAEKQGNVEVAVKAARVAVEAAKQAGDQDVLRKVS<br>EQAERISKEAKKQGNSEVSEEARKVADEAKKQTGDGWSENLYFQG<br>SGHHHHHH |
| BGL02_A17 | BGL02 | DHR68 | AB | GSPDLFLRALELLIKLAESLAETARERGDEEALEIAARIAEEAAEAAER<br>LARKARKEGNLELALQALRAMVEAARVLAEIARERGNEELLKYAWEL<br>AEEAARQAMEIALEAFKRGNWELALIALEILVEALKVLSHIAREKGDP<br>ELLDLSMAAQLVLVIAALIVKAAALQEQGNKEEAEKVLRKAREAIREV<br>TKRLEEIAKNARTPEQALRAAELLVILIDLLILIAKLLQEQGNKEEAEKV<br>LREATELIKRVTELLEKIAKNSDTPELALRAAELLVRLIKLLIEIAKLLQE<br>QGNKEEAEKVLREATELIKRVTELLEKIAKNSDTPELAKRAAELLKRLI<br>ELLKEIAKLLEEEGNEDEAEKVKEEAKELEERVRELEERIRKNSDTG<br>WSENLYFQGSGHHHHHH |
| BGL02_A18 | BGL02 | DHR82 | AB | GSPDLFLKALELLIKLAESLAETARERGDEKALEEAARIAERAAELAEK<br>LARKARKEGNLELALQALRAMVEAARVLAEIARERGNEELLKYAWEL<br>AREAARQAKEIAVEALLRGNWELALIALEILVEALKVLSHIAREKGDPK<br>LLEESMEEQRALVVLVKLAEAAVAVADVEEAIKRNPDNDEAVETAVR<br>LARELKKIAEKLQELAKRKGDPRLLAAALVALKLAVRAVELAIKSNPD<br>NDEAVETAVRLARELKKVAEELQERAKKTGDPELLKLALEALEVAVR<br>AVELAIKSNPDNEEAVETAKRLAEELRKVAELLEERAKETGDPELQEL<br>AKRAKEVADRARELAKKSNPNNGWSENLYFQGSGHHHHHH |
| BGL03_A19 | BGL03 | DHR49 | AB | GSLDLVLQNLELLVHIAEVLARLARRTGNEEALEHAARVAEEVARQA<br>EEIAREARKQGNLELALEALRIMVEAARVLAEIARERGNEELLKYAHE<br>LAEKAAKEAWEIARQAAEEGDLELANKALRIALEAIRVLIEHDQSEEA<br>AKHLIEALEALLVASKISMIGARAKESGTEESLRQAIEDVAQLAKKATG<br>SEALQFAIAIILAIAKASGSEEALRQAIRAVAEIAKEAQDSEVLEFAIRVI<br>LLIAKESGSEEALRQAIRAVAEIAKEAQDPRVLEEAIRVIRQIAEESGS<br>EEARRQAERAEEEIRRRAQGWSENLYFQGSGHHHHHH |
| BGL03_A20 | BGL03 | DHR71 | AB | GSDDLVLQNLELLVHIAEVLARLARRTGNEEALEHAARVAEEVARQA<br>EEIAREARKQGNLELALEALRIMVEAARVLAEIARERGNEELLKYAHE<br>LAEKAAKEAEEIAKEAAKQGNLELFNKALRILLEAIRVLIEHDQSEEAA<br>KHLIERLERLLKASEASMVGAKARKSLERAREASERGDEEEFRKAAE<br>KALELAKELVERAKQGRLAAMVLLAAKVALIVAELAAKNGDKEVFKK<br>AAESALEVAKRLVEVASKEGDPEMVLEAAKVALRVAELAAKNGDKE<br>VFKKAAESALEVAKRLVEVASKEGDPELVEEAAKVAEEVRKLAKKQG<br>DEEVYEKARETAREVKEELKRVREEKGWSENLYFQGSGHHHHHH |
| BGL03_A21 | BGL03 | DHR72 | AB | GSLDLVLQNLELLVHIAEVLARLARRTGNEEALEIAARVAEDVARKAE<br>EIAREARKQGNLELALEALRIMVEAARVLAEIARERGNEELLKYAHEL<br>AEKAAKEAMEIARQAAEEGNLELLNKALRILLEAIRVLIEHDQSEEAAR<br>ELITRLLALLAMSIAAAVGDKELVRLAAQAAREGDSEKAKAILLAAKAA<br>LVAKEVGDPELIKLALEAARRGDSEKAKAILLAAEAARVAKEVGDPEL<br>IKLALEAARRGDSEKARAILEAAERAREAKERGDPEQIKKARELAKR<br>GDGWSENLYFQGSGHHHHHH |

| | | | | |
|---|---|---|---|---|
| BGL05_A22 | BGL05 | DHR59 | AB | GSLDLDLKALELFVNLAESLARSARKQGNEEALEKAARIAEEAARQAEEIARKARKEGNLELALEALRIMVEAARVLAEIARERGNKELWEYARELATEAARQAMEILEEALRRGNIELALIAIEILIEVNQVLSEGGLPIEESLAELAAIEAAALGKAEVAREALKVLREARELAKENRSEEALKVVREIARAAAAAAQAAAEGKTEVAKLALKVLEEAIELAKENRSEEALKVVLEIARAALAAAQAAEEGKSDEARDALRRLEEAIEEAKENRSKESLEKVREEAKEAEQQAEDAREGKGWSENLYFQGSGHHHHHH |
| BGL05_A23 | BGL05 | DHR64 | AB | GSLDLDLIALELFVNLAESLARSARKQGNEEALEEAARIAEEAARQAEEIARKARKEGNLELALEALRIMVEAARVLAEIARERGNKELWEYARKLATEAARQAAEIAAEAISQGNIELALIAIEILIEVNQVLSEGGLPIEKSLLELAVILALLRAARAAANGDPETALRAAEDMVRLAEEALRRAKESGDEEALEKALRIAELAARLARAVLALAEEQGDPEVALRAVELVVRVAELLLRIAKESGSEEALERALRVAEEAARLAKRVLELAEKQGDPEVARRAVELVKRVAELLERIARESGSEEAKERAERVREEARELQERVKELREREGGWSENLYFQGSGHHHHHH |
| BGL05_A24 | BGL05 | DHR76 | AB | GSLRLDLIALELFVNLAESLARSARKQGNEEALEEAARIAEEAARQAEEIARKARKEGNLELALEALRIMVEAARVLAEIARERGNKELWEYARKLAKEAARQAFEILAEAAARGNIELALIAIEILIEVNQVLSEGGLPIEESLAELRLILLALLLARAAAKAVEEATKQGNPELVEWVRRAVRVAAEALKVADQAWKEGNKDLFRAASELVRAVIEAIEEAVKQGNPELVEWVARAAKVAAEVIKVAIQAEKEGNRDLFRAALELVRAVIEAIEEAVKQGNPELVERVARLAKKAAELIKRAIRAEKEGNRDERREALERVREVIERIEELVRQGNGWSENLYFQGSGHHHHHH |
| BGL06_A25 | BGL06 | DHR46 | AB | GSDDLLLKLLELLVEQARVSAEFARRQGDEKMLEEVARKAEEVARKAEEIARKARKEGNLELALKALEILVRAAHVLAEIARERGNEELLKYAWKLAKEALRQVLKIAAEAAVRGNLELAIIALHISVRIAEVLLETRPDDREEIREQQKMFEILILVLSSDAEEIREAVRLAEELLRKDPSEEAEAILKAAIAAAVRAPDPEAIREAVRAAEELLRENPSTEAEELLRLAIEAAVRAPDPEAIREAVRAAEELLRENPSEEAKELLRRAIESAKKAPDPEAQREAKRAEEELRKEDPSGWSENLYFQGSGHHHHHH |
| BGL06_A26 | BGL06 | DHR68 | AB | GSADLLLKLLELLVEQARVSAEFARRQGDEKMLEEVARKAEEVARKAEEIARKARKEGNLELALKALEILVRAAHVLAEIARERGNEELLKYAWKLAKEALRQVIKIAALAAMEGNLELAIIALHISVRIAEVLLETRPDDREEIRKQQKIFETLIRLLEAAVKVEEIRELIDKARKLQEQGNKEEAEKVLREAREQIREVTRELEDIAKKADTPAIAAAAAALLVKLIKLLIEIAKLLQEQGNKEEAEKVLREATELIKRVTELLEKIAKNSDTPEVALFAAELLVRLIKLLIEIAKLLQEQGNKEEAEKVLREATELIKRVTELLEKIAKNSDTPELAKRAAELLKRLIELLKEIAKLLEEEGNEDEAEKVKEEAKELEERVRELEERIRKNSDTGWSENLYFQGSGHHHHHH |
| BGL06_A27 | BGL06 | DHR77 | AB | GSADLLLKLLELLVEQARVSAEFARRQGDEKMLEEVARKAEEVARKAEEIARKARKEGNLELALKALEILVRAAHVLAEIARERGNEELLKYAWKLAKEALRQVLKIAYEAIREGNLELAIIALHISVRIAEVLLETRPDDREEIRKQQEEFEILIELLEVMVKLKEQAEEARRKKDSEEAEAVYWASRAVQAAAEAMLQAQEEGDEDARRVAEELARQAREAAEKKNSEEAEAVYWAARAVLAALEALEQAKREGDEDARRVAEELLRQAEEAARKKNPEEARAVYEAARDVLEALQRLEEAKRRGDEEERREAEERLRQAEERARKKNGWSENLYFQGSGHHHHHH |
| BGL08_A28 | BGL08 | DHR71 | AB | GSLDQLLKLLELIVKIVESIARFARRQGNEKALEEAAQWAELAAEVAEKVAREARKKGNLELALKALQIMVEAARVLAEIARERGNEELAKYALKLALEAARQALEIAIEAWQRGNNELALIALEILVEVHNVILEQHPRSEELKAIHEILKALLIVALASLRGDPEKVLEAAKEALRLAELAAKKGDKDTFRLAALVALLIAKILVEVASKEGDPELVLEAAKVALRVAELAAKNGDKEVFKRAAESALEVAKRLVEVASKEGDPELVEEAAKVAEEVRKLAKKQGDEEVYEKARETAREVKEELKRVREEKGWSENLYFQGSGHHHHHH |

| | | | | |
|---|---|---|---|---|
| BGL08_A29 | BGL08 | DHR77 | AB | GSLDQLLKLLELIVKIVESIARFARRQGNEKSLELAARWAEKAAEVAEKVAREARKKGNLELALKALQIMVEAARVLAEIARERGNEELAKYALKLAFEAARQAADIALEALERGNKELALIALEILVEVHNVILEQHPRSEALKIIHLALQAALAALIAGVEDALKVAILLAKQAAEAYEKKDSEEREAVEWAMRAVKAALEALEQAKREGDEDARRVAEELLRQAEEAARKKNSEEAEAVYWAARAVLAALEALEQAKREGDEDARRVAEELLRQAEEAARKKNPEEARAVYEAARDVLEALQRLEEAKRRGDEEERREAEERLRQAEERARKKNGWSENLYFQGSGHHHHHH |
| BGL08_A30 | BGL08 | DHR79 | AB | GSLDQLLKLLELIVKIVESIARFARRQGNEKSLEDAARWAERAAEVAEIVARIARKRGNLELALKALQIMVEAARVLAEIARERGNEELAKYALKLAREAARQAREIASEAMRRGNVELALIALEILVEVHNVILEQHPRSEELKEIHEILKVALEALIAAVEGEAVAEQTGDPRVKELAREMTRLLAEAIALFLKDPSSSDVREALKLIAEAAKAAARALKAAERTGDPEVRELARELVRLAVEAAEEVQRNPSSSDVNEALKLIVEAIEAAVRALEAAERTGDPEVRELARELVRLAVEAAEEVQRNPSSEEVNEALKKIVKAIQEAVESLREAEESGDPEKREKARERVREAVERAEEVQRDPGWSENLYFQGSGHHHHHH |
| BGL17_A31 | BGL17 | DHR59 | AB | GSPELFLQDLRSLVEAARILARLARQRGDEHALERAARWAEQAARQAEKLARQARKEGNLELALKALQILVNAAYVLAEIARDRGNEELLEYAARLAEEAARQAAEIWAEAARRGNQQLRTKAAHILLRAAEVLLEIARDRGNQELLEKAQRIVEAVAAAQQVAALALRLAEELDSEEAKKAVRAIAEAAAAALLAALQGKDEVAKLALKVLKEAIELAKENRSEEALKVVLEIARAAAAARAAEEGKTEVAKLALKVLEEAIELAKENRSEEALKVVLEIARAALAAAQAAEEGKSDEARDALRRLEEAIEEAKENRSKESLEKVREEAKEAEQQAEDAREGKGWSENLYFQGSGHHHHHH |
| BGL17_A32 | BGL17 | DHR62 | AB | GSPELFLQDLRSLVEAARILARLARQRGDEHALERAARWAEQAARQAERLARQARKEGNLELALKALQILVNAAYVLAEIARDRGNEELLEYAARLAEEAARQAIEIWAQAMEEGNQQLRTKAAHIILRAAEVLLEIARDRGNQELLEKAASLVDAVAALQQAAAAILEGDVEKAVRAAQEAVKAAKEAGDNDMLRAVAIAALRIAKEAEKQGNVEVAVKAARVAVEAAKQAGDNDVLRKVAEQALRIAKEAEKQGNVEVAVKAARVAVEAAKQAGDQDVLRKVSEQAERISKEAKKQGNSEVSEEARKVADEAKKQTGDGWSENLYFQGSGHHHHHH |
| BGL17_A33 | BGL17 | DHR76 | AB | GSPELFLQDLRSLVEAARILARLARQRGDEHALEKAARWAEQAARQAEKLARQARKEGNLELALKALQILVNAAYVLAEIARDRGNEELLEYAARLAEEAARQAWEIWAQAAEEGNQQLRTKAAHILLRAAEVLLEIARDRGNQELLEKAQKLVEWVALLQQLAAELLRAVKEAIEEAKKQGNPELVEWVARAAKVALKVLALAAKAALEGNDDLSRAAKELARAVIEAIEEAVKQGNPELVEWVARAAKVAAEVIKVAIQAEKEGNRDLFRAALELVRAVIEAIEEAVKQGNPELVERVARLAKKAAELIKRAIRAEKEGNRDERREALERVREVIERIEELVRQGNGWSENLYFQGSGHHHHHH |
| BGL18_A34 | BGL18 | DHR24 | AB | GSPELVLLALENMVRAAHTLAEIARDNGNEEWLERAARLAEEVARMAEELARQARKEGNLELALKALQILVNAAYVLAEIARDRGNEELLEYAARLAEEAARQAEKIALEAAAQGNFELALEALEILNEAARVLARIAHHRGNQELLEKAWRLTEKSAIMSKMIALMAELRGRYPDSEAAKLALKAAKEAIKLMEQSTDEELAKELAKLAEKLAELAKRYPDSEAAKLALKAALEAIELCKQSTDEELCEELVKLAQKLIELAKRYPDSEEAKRALKEAKELIEQCKESTDEDECRELVKRAEELIREAKENPDGWSENLYFQGSGHHHHHH |
| BGL18_A35 | BGL18 | DHR26 | AB | GSPELVLKALENMVRAAHTLAEIARDNGNEEWLEAAARLAELVAKAAEELAREARKEGNLELALKALQILVNAAYVLAEIARDRGNEELLEYAARLAEEAARQALEIASKAAKKGNFELALEALRILNEAARVLARIAHHRGNQELLKKAWLLTAMSAAMSALIALAAKSTDEEVRKIAARVAEELAQLAEEALRSGSEECIRLAEEVIKAVLLLAALAWKATDEEVIRVALEVARELIR |

| | | | | |
|---|---|---|---|---|
| | | | | LAQEACRSNDDECLRLASEVVKAVQELVKLAEQATDEEVIRVALEVARELIRLAQEACRSNDEECLREASEVVKEVQELVKEAEKSTDEEEIRELLQRAEERIREAQERCREGDGWSENLYFQGSGHHHHHH |
| BGL18_A36 | BGL18 | DHR49 | AB | GSPELVLKALENMVRAAHTLAEIARDNGNEEWLERAARLAELVARLAEKLARKARKEGNLELALKALQILVNAAYVLAEIARDRGNEELLEYAARLAEEAARQAKEIAAKAASKGNFELALEALRILNEAARVLARIAHHRGNQELLEKAWRLTEASALASLLIAIAAEAKDSGTEESLRQAIELAAQIAKKADDKAVLIAAIAIIRVIAEESGSEEALRQAIRAVAEIAKEAQDSEVLEAAIRVILLIAKESGSEEALRQAIRAVAEIAKEAQDPRVLEEAIRVIRQIAEESGSEEARRQAERAEEEIRRRAQGWSENLYFQGSGHHHHHH |
| BGL19_A37 | BGL19 | DHR14 | AB | GSPRWYLQHLETLVRAAEILAELARQRGDEEALEEAARMAEEAARKAEELAREARKEGNLELALKALQILVNAAYVLAEIARDRGNEELLEYAARLAEEAARQAKEIAEEAMKQGDIQLAIKALRILLEAAHVLMQIARDRGNEELLEKAKRLVREAAELSKLAMEALRLAKKAKEATDKEEVIEIVKELAELAKQATDKALVALIVKQLAEVAKEATDKELVIYIVKILAELAKQSTDSELVNTIVKQLAEVAKEATDKELVIYIVKILAELAKQSTDSELVNEIVKQLEEVAKEATDKELVEHIEKILEELKKQSTDGWSENLYFQGSGHHHHHH |
| BGL19_A38 | BGL19 | DHR71 | AB | GSPRWYLQHLETLVRAAEILAELARQRGDEEALEEAARMAEEAARAAEKLAREARKEGNLELALKALQILVNAAYVLAEIARDRGNEELLEYAARLAEEAARQAAEIAARALQEGDVQLAIKALRILLEAAHVLMQIARDRGNEELLEKAKILVKAAAMASKAYMLAQQAKEEGDPELVLEAAKLALELAERAAKKGDKAALRMAAALADALARLLVEVASKEGDPELVLEAAKVALRVAELAAKNGDKEVFKKAAESALEVAKRLVEVASKEGDPELVEEAAKVAEEVRKLAKKQGDEEVYEKARETAREVKEELKRVREEKGWSENLYFQGSGHHHHHH |
| BGL19_A39 | BGL19 | DHR77 | AB | GSPRWYLQHLETLVRAAEILAELARQRGDEEALEEAARMAEHAARAAEKLAREARKEGNLELALKALQILVNAAYVLAEIARDRGNEELLEYAARLAEEAARQALEIAAQALEEGDMQLAIKALRILLEAAHVLMQIARDRGNEELLEKAKRLVRLAARISEMVMELQRAKEEARRKKDSEEAEAAYWAQKAIAAALAALAQALEEGDEDARRVAEELLRQAEEAARKKNSEEAEAVYWAAEAVLAALEALEQAKREGDEDARRVAEELLRQAEEAARKKNPEEARAVYEAARDVLEALQRLEEAKRRGDEEERREAEERLRQAEERARKKNGWSENLYFQGSGHHHHHH |

**Table S4.** Designed sequences of interface transplanted C3 homotrimers.

| ID | Host | Guest | Protomer seq |
|---|---|---|---|
| BGL17_A11 | BGL00_A11 | BGL17 | MGPELFLQDLRSLVEAARILARLARQRGDEHALRRAARWAEQAARLAEELARQARKEGNLELALKALQILVNAAYVLAEIARDRGNEELLEYAARLALKAAVMALEIWAQALKEGNQQLRTKAAHILLRAAEVLLEIARDRGNQELLRIAQILVDAVAAFQLFAANPGSNEAKKAAELVLRLAEELAKSPDPEALKAAIAMAEAVVALNPGSNLAKKALEIILRAAEELAKLPDPEALKEAVKAAEKVVREQPGSELAKKALEIIERAAEELKKSPDPEAQKEAKKAEQKVREERPGGSENLYFQGSHHHHHH |
| BGL17_A35 | BGL18_A35 | BGL17 | MGPELFLKDLRSLVEAARILARLARQRGDEHALEAAARWAELAAKAAEELARQARKEGNLELALKALQILVNAAYVLAEIARDRGNEELLEYAARLAEEAARQALEIWSKAAKKGNQQLRTKAARILLRAAEVLLEIARDRGNQELLKKAQLLVAMVAAMQALAALAIKSTDKEVRKIAARVAEELAQLAEEALRSGSEECIRLAEEVIKAVLLLAALAWKATDEEVIRVALEVARELIRLAQEACRSNDDECLRLASEVVKAVQELVKLAEQATDEEVIRVALEVARELIRLAQEACRSNDEECLREASEVVKEVQELVKEAEKSTDEEEIRELLQRAEERIREAQERCREGDGSENLYFQGSHHHHHH |
| BGL17_A39 | BGL19_A39 | BGL17 | MGPELFLQDLRSLVEAARILARLARQRGDEHALERAARWAEQAARAAEKLARQARKEGNLELALKALQILVNAAYVLAEIARDRGNEELLEYAARLAEEAARQALEIAAQALEEGNMQLRTKAAHILLRAAEVLLEIARDRGNQELLEKAQRLVDLVARI |

| | | | |
|---|---|---|---|
| | | | QQMVARLIRAKEEARRKKDSEEAEAAYWAQKAIAAALAALAQALEEGDEDAR<br>RVAEELLRQAEEAARKKNSEEAEAVYWAAEAVLAALEALEQAKREGDEDARR<br>VAEELLRQAEEAARKKNPEEARAVYEAARDVLEALQRLEEAKRRGDEEERRE<br>AEERLRQAEERARKKNGSENLYFQGSHHHHHH |
| BGL18<br>_A10 | BGL00<br>_A10 | BGL18 | MGPRLVLRALENMVRAAHTLAEIARDNGNEEWLRRAARLAEEVARAAEELAR<br>EARKEGNLELALKALQILVNAAYVLAEIARDRGNEELLEYAARLAEEAARQAW<br>EIAAEALERGNFELALEALEILNEAARVLARIAHHRGNQELLEKAWRLTHLSALA<br>SRVIAELARLKEEACRSNSDECLRLASEVEKAVAKLARLAAKATDEEVRRVAL<br>EVARELIKLAQEACRSNDDECLRLASEVVKAVQELVKLAEQATDEEVIRVALEV<br>ARELIRLAQEACRSNDEECLREASEVVKEVQELVKEAEKSTDEEEIRELLQRA<br>EERIREAQERCREGDGSENLYFQGSHHHHHH |
| BGL18<br>_A11 | BGL00<br>_A11 | BGL18 | MGPRLVLRALENMVRAAHTLAEIARDNGNEEWLRRAARLAEEVARLAEELAR<br>EARKEGNLELALKALQILVNAAYVLAEIARDRGNEELLEYAARLALKAAVMALEI<br>AAQALKEGNFELALEALEILNEAARVLARIAHHRGNQELLRIAWILTHASAAFSL<br>FIANPGSNEAKKAAELVLRLAEELAKSPDPEALKAAIAMAEAVVALNPGSNLAK<br>KALEIILRAAEELAKLPDPEALKEAVKAAEKVVREQPGSELAKKALEIIERAAEEL<br>KKSPDPEAQKEAKKAEQKVREERPGGSENLYFQGSHHHHHH |
| BGL18<br>_A31 | BGL17<br>_A31 | BGL18 | MGPRLVLRALENMVRAAHTLAEIARDNGNEEWLERAARLAEEVARRAEKLAR<br>EARKEGNLELALKALQILVNAAYVLAEIARDRGNEELLEYAARLAEEAARQAAE<br>IAAEAARRGNFELALEALEILNEAARVLARIAHHRGNQELLEKAWRITEASAAA<br>SRVIALALRLAEELDSEEAKKAVRAIAEAAAAALLAALQGKDEVAKLALKVLKEA<br>IELAKENRSEEALKVVLEIARAAAAAARAAEEGKTEVAKLALKVLEEAIELAKEN<br>RSEEALKVVLEIARAALAAAQAAEEGKSDEARDALRRLEEAIEEAKENRSKESL<br>EKVREEAKEAEQQAEDAREGKGSENLYFQGSHHHHHH |
| BGL18<br>_A32 | BGL17<br>_A32 | BGL18 | MGPRLVLRALENMVRAAHTLAEIARDNGNEEWLERAARLAEEVARRAERLAR<br>EARKEGNLELALKALQILVNAAYVLAEIARDRGNEELLEYAARLAEEAARQAIEI<br>AAQAMEEGNFELALEALEIINEAARVLARIAHHRGNQELLEKAASLTHASAALS<br>RAIAAILEGDVEKAVRAAQEAVKAAKEAGDNDMLRAVAIAALRIAKEAEKQGNV<br>EVAVKAARVAVEAAKQAGDNDVLRKVAEQALRIAKEAEKQGNVEVAVKAARV<br>AVEAAKQAGDQDVLRKVSEQAERISKEAKKQGNSEVSEEARKVADEAKKQTG<br>DGSENLYFQGSHHHHHH |
| BGL18<br>_A38 | BGL19<br>_A38 | BGL18 | MGPRLVLRALENMVRAAHTLAEIARDNGNEEWLERAARLAEEVARAAEKLAR<br>EARKEGNLELALKALQILVNAAYVLAEIARDRGNEELLEYAARLAEEAARQAAE<br>IAARALQEGNVELALEALEILNEAARVLARIAHHRGNQELLEKAWILTKASAMA<br>SKAYALAAQAKEEGDPELVLEAAKLALELAERAAKKGDKAALRMAAALADALA<br>RLLVEVASKEGDPELVLEAAKVALRVAELAAKNGDKEVFKKAAESALEVAKRL<br>VEVASKEGDPELVEEAAKVAEEVRKLAKKQGDEEVYEKARETAREVKEELKR<br>VREEKGSENLYFQGSHHHHHH |
| BGL18<br>_A39 | BGL19<br>_A39 | BGL18 | MGPRLVLRALENMVRAAHTLAEIARDNGNEEWLERAARLAEEVARAAEKLAR<br>EARKEGNLELALKALQILVNAAYVLAEIARDRGNEELLEYAARLAEEAARQALEI<br>AAQALEEGNMELALEALEILNEAARVLARIAHHRGNQELLEKAWRLTHLSAKIS<br>RMVAELARAKEEARRKKDSEEAEAAYWAQKAIAAALAALAQALEEGDEDARR<br>VAEELLRQAEEAARKKNSEEAEAVYWAAEAVLAALEALEQAKREGDEDARRV<br>AEELLRQAEEAARKKNPEEARAVYEAARDVLEALQRLEEAKRRGDEEERREA<br>EERLRQAEERARKKNGSENLYFQGSHHHHHH |
| BGL19<br>_A10 | BGL00<br>_A10 | BGL19 | MGPRWYLQHLETLVRAAEILAELARQRGDEEALREAARMAEHAARAAEELAR<br>EARKEGNLELALKALQILVNAAYVLAEIARDRGNEELLEYAARLAEEAARQAW<br>EIAAEALERGDNQLAIKALRILLEAAHVLMQIARDRGNEELLEKAKRLVRLAALA<br>SEVVMELQRLKEEACRSNSDECLRLASEVEKAVAKLARLAAKATDEEVRRVAL<br>EVARELIKLAQEACRSNDDECLRLASEVVKAVQELVKLAEQATDEEVIRVALEV<br>ARELIRLAQEACRSNDEECLREASEVVKEVQELVKEAEKSTDEEEIRELLQRA<br>EERIREAQERCREGDGSENLYFQGSHHHHHH |

| | | | |
|---|---|---|---|
| BGL19_A11 | BGL00_A11 | BGL19 | MGPRWYLQHLETLVRAAEILAELARQRGDEEALREAARMAEHAARLAEELAREARKEGNLELALKALQILVNAAYVLAEIARDRGNEELLEYAARLALKAAVMALEIAAQALREGDNQLAIKALRILLEAAHVLMQIARDRGNEELLRIAKILVRAAAAFSLFVMNPGSNEAKKAAELVLRLAEELAKSPDPEALKAAIAMAEAVVALNPGSNLAKKALEIILRAAEELAKLPDPEALKEAVKAAEKVVREQPGSELAKKALEIIERAAEELKKSPDPEAQKEAKKAEQKVREERPGGSENLYFQGSHHHHHH |
| BGL19_A31 | BGL17_A31 | BGL19 | MGPRWYLQHLETLVRAAEILAELARQRGDEEALEEAARMAEHAARQAEKLAREARKEGNLELALKALQILVNAAYVLAEIARDRGNEELLEYAARLAEEAARQAAEIAAEAARRGDNQLAIKALRILLEAAHVLMQIARDRGNEELLEKAKRIVEAAAAASEVVMLALRLAEELDSEEAKKAVRAIAEAAAAALLAALQGKDEVAKLALKVLKEAIELAKENRSEEALKVVLEIARAAAAAARAAEEGKTEVAKLALKVLEEAIELAKENRSEEALKVVLEIARAALAAAQAAEEGKSDEARDALRRLEEAIEEAKENRSKESLEKVREEAKEAEQQAEDAREGKGSENLYFQGSHHHHHH |
| BGL19_A32 | BGL17_A32 | BGL19 | MGPRWYLQHLETLVRAAEILAELARQRGDEEALEEAARMAEHAARQAERLAREARKEGNLELALKALQILVNAAYVLAEIARDRGNEELLEYAARLAEEAARQAIEIAAQAMEEGDNQLAIKALRIILEAAHVLMQIARDRGNEELLEKAASLVRAAAALSEAVMAILEGDVEKAVRAAQEAVKAAKEAGDNDMLRAVAIAALRIAKEAEKQGNVEVAVKAARVAVEAAKQAGDNDVLRKVAEQALRIAKEAEKQGNVEVAVKAARVAVEAAKQAGDQDVLRKVSEQAERISKEAKKQGNSEVSEEARKVADEAKKQTGDGSENLYFQGSHHHHHH |
| BGL19_A35 | BGL18_A35 | BGL19 | MGPEWYLKHLETLVRAAEILAELARQRGDEEALEAAARMAELAAKAAEELAREARKEGNLELALKALQILVNAAYVLAEIARDRGNEELLEYAARLAEEAARQALEIASKAARKGDNQLAIKALRILLEAAHVLMQIARDRGNEELLKKAKLLVAMAAAMSALVMLAQKSTDKEVRKIAARVAEELAQLAEEALRSGSEECIRLAEEVIKAVLLLAALAWKATDEEVIRVALEVARELIRLAQEACRSNDDECLRLASEVVKAVQELVKLAEQATDEEVIRVALEVARELIRLAQEACRSNDEECLREASEVVKEVQELVKEAEKSTDEEEIRELLQRAEERIREAQERCREGDGSENLYFQGSHHHHHH |
| BGL00_A32 | BGL17_A32 | BGL00 | MGLELALKALQILVNAAYVLAEIARDRGNEELLEKAARLAEEAARQAERIARQARKEGNLELALKALQILVNAAYVLAEIARDRGNEEELEYAARLAEEAARQAIEIAAQAMEEGNLELALKALQIIVNAAYVLAEIARDRGNEELLEKAASLAEAAAALAEAIAAILEGDVEKAVRAAQEAVKAAKEAGDNDMLRAVAIAALRIAKEAEKQGNVEVAVKAARVAVEAAKQAGDNDVLRKVAEQALRIAKEAEKQGNVEVAVKAARVAVEAAKQAGDQDVLRKVSEQAERISKEAKKQGNSEVSEEARKVADEAKKQTGDGSENLYFQGSHHHHHH |
| BGL00_A35 | BGL18_A35 | BGL00 | MGLELALKALQILVNAAYVLAEIARDRGNEELLEAAARLAELAAKAAEEIARQARKEGNLELALKALQILVNAAYVLAEIARDRGNEEELEYAARLAEEAARQALEIASKAAKKGNLELALKALRILVNAAYVLAEIARDRGNEELLKKAALLAAMAAAMAALIALAAKSTDNEVRKIAARVAEELAQLAEEALRSGSEECIRLAEEVIKAVLLLAALAWKATDEEVIRVALEVARELIRLAQEACRSNDDECLRLASEVVKAVQELVKLAEQATDEEVIRVALEVARELIRLAQEACRSNDEECLREASEVVKEVQELVKEAEKSTDEEEIRELLQRAEERIREAQERCREGDGSENLYFQGSHHHHHH |
| BGL00_A38 | BGL19_A38 | BGL00 | MGLELALKALQILVNAAYVLAEIARDRGNEELLEKAARLAEEAARAAEKIARQARKEGNLELALKALQILVNAAYVLAEIARDRGNEEELEYAARLAEEAARQAAEIAARALQEGNVELALKALQILVNAAYVLAEIARDRGNEELLEKAAILAKAAAMAAKAYALAAQAKEEGDPELVLEAAKLALELAERAAKKGDKAALRMAAALADALARLLVEVASKEGDPELVLEAAKVALRVAELAAKNGDKEVFKKAAESALEVAKRLVEVASKEGDPELVEEAAKVAEEVRKLAKKQGDEEVYEKARETAREVKEELKRVREEKGSENLYFQGSHHHHHH |
| BGL00_A39 | BGL19_A39 | BGL00 | MGLELALKALQILVNAAYVLAEIARDRGNEELLEKAARLAEEAARAAEKIARQARKEGNLELALKALQILVNAAYVLAEIARDRGNEEELEYAARLAEEAARQALEIAAQALEEGNMELALKALQILVNAAYVLAEIARDRGNEELLEKAARLAELAARIAEMVARLARAKEEARRKKDSEEAEAAYWAQKAIAAALAALAQALEEGDEDARRVAEELLRQAEEAARKKNSEEAEAVYWAAEAVLAALEALEQAKREGDEDARR |

| | | | |
|---|---|---|---|
| | | | VAEELLRQAEEAARKKNPEEARAVYEAARDVLEALQRLEEAKRRGDEEERR<br>EAEERLRQAEERARKKNGSENLYFQGSHHHHHH |

**Table S5.** Designed sequences of pseudo-C3 heterotrimers (host: BGL17_A32).

| ID | Guest | Chain | Protomer sequence |
|---|---|---|---|
| hetBGL00-17-18_A32 | BGL00<br>BGL17<br>BGL18 | A | MGLELALKALQILVNAAYVLAEIARDRGNEELLEKAARLAEEAARQAERI<br>ARQARKEGNLELALKALQILVNAAYVLAEIARDRGNEELLEYAARLAEEA<br>ARQAIEIAAQAMEEGNFELALEALEIINEAARVLARIAHHRGNQELLEKAA<br>SLTHASAALSRAIAAILEGDVEKAVRAAQEAVKAAKEAGDNDMLRAVAIA<br>ALRIAKEAEKQGNVEVAVKAARVAVEAAKQAGDQDVLRKVSEQAERISK<br>EAKKQGNSEVSEEARKVADEAKKQTGDG |
| | | B | MGPELFLQDLRSLVEAARILARLARQRGDEHALERAARWAEQAARQAE<br>RLARQARKEGNLELALKALQILVNAAYVLAEIARDRGNEEELEYAARLAE<br>EAARQAIEIAAQAMEEGNLELALKALQIIVNAAYVLAEIARDRGNEELLEK<br>AASLAEAAAALAEAIAAILEGDVEKAVRAAQEAVKAAKEAGDNDMLRAV<br>AIAALRIAKEAEKQGNVEVAVKAARVAVEAAKQAGDNDVLRKVAEQALR<br>IAKEAEKQGNVEVAVKAARVAVEAAKQAGDQDVLRKVSEQAERISKEA<br>KKQGNSEVSEEARKVADEAKKQTGDG |
| | | C | MGPRLVLRALENMVRAAHTLAEIARDNGNEEWLERAARLAEEVARRAE<br>RLAREARKEGNLELALKALQILVNAAYVLAEIARDRGNEELLEYAARLAE<br>EAARQAIEIWAQAMEEGNQQLRTKAAHIILRAAEVLLEIARDRGNQELLE<br>KAASLVDAVAALQQAAAAILEGDVEKAVRAAQEAVKAAKEAGDNDMLR<br>AVAIAALRIAKEAEKQGNVEVAVKAARVAVEAAKQAGDNDVLRKVAEQA<br>LRIAKEAEKQGNVEVAVKAARVAVEAAKQAGDQDVLRKVAEQALRIAKE<br>AEKQGNVEVAVEAARVAVEAAKQAGDQDVLRKVSEQAERISKEAKKQG<br>NSEVSEEARKVADEAKKQTGDGSENLYFQGSHHHHHHG |
| hetBGL0-17-19_A32 | BGL00<br>BGL17<br>BGL19 | A | MGLELALKALQILVNAAYVLAEIARDAGNEELLEKAARLAEEAARQAERI<br>ARQARKEGNLELALKALQILVNAAYVLAEIARDRGNEELLEYAARLAEEA<br>ARQAIEIEAQAMEEGDNQLAIKALRIILEAAHVLMQIARDRGNEELLEKAA<br>SLVRAAAALSEARMAILEGDVEKAVRAAQEAVKAAKEAGDNDMLRAVAI<br>AALRIAKEAEKQGNVEVAVKAARVAVEAAKQAGDQDVLRKVSEQAERIS<br>KEAKKQGNSEVSEEARKVADEAKKQTGDG |
| | | B | MGPELFLQDLRSLVEAARILARLARQRGDEHALERAARWAEQAARQAE<br>RLARQARKEGNLELALKALQILVNAAYVLAEIARDRGNEEELEYAARLAE<br>EAARQAIEIAAQAMEEGNLELALKALQIIVNAAYVLAEIAADRGNEELLEK<br>AASLAEAAAALAEAIAAILEGDVEKAVRAAQEAVKAAKEAGDNDMLRAV<br>AIAALRIAKEAEKQGNVEVAVKAARVAVEAAKQAGDNDVLRKVAEQALR<br>IAKEAEKQGNVEVAVKAARVAVEAAKQAGDQDVLRKVSEQAERISKEA<br>KKQGNSEVSEEARKVADEAKKQTGDG |
| | | C | MGPRWYLQHLETLVRAAEILAELARQRGDEEALEEAARMAEHAARQAE<br>RLAREARKEGNLELALKALQILVNAAYVLAEIARDRGNEELLEYAARLAE<br>EAARQAIEIWAQAMEEGNQQLRTKAAHIILRAAEVLLEIARDRGNQELLE<br>KAASLVDAVAALQQAAAAILEGDVEKAVRAAQEAVKAAKEAGDNDMLR<br>AVAIAALRIAKEAEKQGNVEVAVKAARVAVEAAKQAGDNDVLRKVAEQA<br>LRIAKEAEKQGNVEVAVKAARVAVEAAKQAGDQDVLRKVAEQALRIAKE<br>AEKQGNVEVAVEAARVAVEAAKQAGDQDVLRKVSEQAERISKEAKKQG<br>NSEVSEEARKVADEAKKQTGDGSENLYFQGSHHHHHHG |
| hetBGL0-18-17_A32 | BGL00<br>BGL18<br>BGL17 | A | MGLELALKALQILVNAAYVLAEIARDRGNEELLEKAARLAEEAARQAERI<br>ARQARKEGNLELALKALQILVNAAYVLAEIARDRGNEELLEYAARLAEEA<br>ARQAIEIWAQAMEEGNQQLRTKAAHIILRAAEVLLEIARDRGNQELLEKA<br>ASLVDAVAALQQAAAAILEGDVEKAVRAAQEAVKAAKEAGDNDMLRAV |

| | | | |
|---|---|---|---|
| | | | AIAALRIAKEAEKQGNVEVAVKAARVAVEAAKQAGDQDVLRKVSEQAER<br>ISKEAKKQGNSEVSEEARKVADEAKKQTGDG |
| | | B | MGPRLVLRALENMVRAAHTLAEIARDNGNEEWLERAARLAEEVARRAE<br>RLAREARKEGNLELALKALQILVNAAYVLAEIARDRGNEEELEYAARLAE<br>EAARQAIEIAAQAMEEGNLELALKALQIIVNAAYVLAEIARDRGNEELLEK<br>AASLAEAAAALAEAIAAILEGDVEKAVRAAQEAVKAAKEAGDNDMLRAV<br>AIAALRIAKEAEKQGNVEVAVKAARVAVEAAKQAGDNDVLRKVAEQALR<br>IAKEAEKQGNVEVAVKAARVAVEAAKQAGDQDVLRKVSEQAERISKEA<br>KKQGNSEVSEEARKVADEAKKQTGDG |
| | | C | MGPELFLQDLRSLVEAARILARLARQRGDEHALERAARWAEQAARQAE<br>RLARQARKEGNLELALKALQILVNAAYVLAEIARDRGNEELLEYAARLAE<br>EAARQAIEIAAQAMEEGNFELALEALEIINEAARVLARIAHHRGNQELLEK<br>AASLTHASAALSRAIAAILEGDVEKAVRAAQEAVKAAKEAGDNDMLRAV<br>AIAALRIAKEAEKQGNVEVAVKAARVAVEAAKQAGDNDVLRKVAEQALR<br>IAKEAEKQGNVEVAVKAARVAVEAAKQAGDQDVLRKVAEQALRIAKEAE<br>KQGNVEVAVEAARVAVEAAKQAGDQDVLRKVSEQAERISKEAKKQGN<br>SEVSEEARKVADEAKKQTGDGSENLYFQGSHHHHHHG |
| hetBGL0-19-17_A32 | BGL00<br>BGL19<br>BGL17 | A | MGLELALKALQILVNAAYVLAEIARDRGNEELLEKAARLAEEAARQAERI<br>ARQARKEGNLELALKALQILVNAAYVLAEIARDRGNEELLEYAARLAEEA<br>ARQAIEIWAQAMEEGNQQLRTKAAHIILRAAEVLLEIARDRGNQELLEKA<br>ASLVDAVAALQQAAAAILEGDVEKAVRAAQEAVKAAKEAGDNDMLRAV<br>AIAALRIAKEAEKQGNVEVAVKAARVAVEAAKQAGDQDVLRKVSEQAER<br>ISKEAKKQGNSEVSEEARKVADEAKKQTGDG |
| | | B | MGPRWYLQHLETLVRAAEILAELARQRGDEEALEEAARMAEHAARQAE<br>RLAREARKEGNLELALKALQILVNAAYVLAEIARDRGNEEELEYAARLAE<br>EAARQAIEIAAQAMEEGNLELALKALQIIVNAAYVLAEIARDRGNEELLEK<br>AASLAEAAAALAEAIAAILEGDVEKAVRAAQEAVKAAKEAGDNDMLRAV<br>AIAALRIAKEAEKQGNVEVAVKAARVAVEAAKQAGDNDVLRKVAEQALR<br>IAKEAEKQGNVEVAVKAARVAVEAAKQAGDQDVLRKVSEQAERISKEA<br>KKQGNSEVSEEARKVADEAKKQTGDG |
| | | C | MGPELFLQDLRSLVEAARILARLARQRGDEHALERAARWAEQAARQAE<br>RLARQARKEGNLELALKALQILVNAAYVLAEIARDRGNEELLEYAARLAE<br>EAARQAIEIEAQAMEEGDNQLAIKALRIILEAAHVLMQIARDRGNEELLEK<br>AASLVRAAAALSEARMAIQEGDVEKAVRAAQEAVKAAKEAGDNDMLRA<br>VAIAALRIAKEAEKQGNVEVAVKAARVAVEAAKQAGDNDVLRKVAEQAL<br>RIAKEAEKQGNVEVAVKAARVAVEAAKQAGDQDVLRKVAEQALRIAKEA<br>EKQGNVEVAVEAARVAVEAAKQAGDQDVLRKVSEQAERISKEAKKQG<br>NSEVSEEARKVADEAKKQTGDGSENLYFQGSHHHHHHG |
| hetBGL17-18-19_A32 | BGL17<br>BGL18<br>BGL19 | A | MGPELFLQDLRSLVEAARILARLARQRGDEHALERAARWAEQAARQAE<br>RLARQARKEGNLELALKALQILVNAAYVLAEIARDRGNEELLEYAARLAE<br>EAARQAIEIMAQAMEEGDNQLAIKALRIILEAAHVLMQIARDRGNEELLEK<br>AASLVRAAAALSEAAMAILEGDVEKAVRAAQEAVKAAKEAGDNDMLRA<br>VAIAALRIAKEAEKQGNVEVAVKAARVAVEAAKQAGDQDVLRKVSEQAE<br>RISKEAKKQGNSEVSEEARKVADEAKKQTGDG |
| | | B | MGPRLVLRALENMVRAAHTLAEIARDNGNEEWLERAARLAEEVARRAE<br>RLAREARKEGNLELALKALQILVNAAYVLAEIARDRGNEELLEYAARLAE<br>EAARQAIEIWAQAMEEGNQQLRTKAAHIILRAAEVLLEIARDRGNQELLE<br>KAASLVDAVAALQQAAAAILEGDVEKAVRAAQEAVKAAKEAGDNDMLR<br>AVAIAALRIAKEAEKQGNVEVAVKAARVAVEAAKQAGDNDVLRKVAEQA<br>LRIAKEAEKQGNVEVAVKAARVAVEAAKQAGDQDVLRKVSEQAERISKE<br>AKKQGNSEVSEEARKVADEAKKQTGDG |
| | | C | MGPRWYLQHLETLVRAAEILAELARQRGDEEALEEAARMAEHAARQAE<br>RLAREARKEGNLELALKALQILVNAAYVLAEIARDRGNEELLEYAARLAE |

| | | | |
|---|---|---|---|
| | | | EAARQAIEIAAQAMEEGNFELALEALEIINEAARVLARIAHHRGNQELLEKAASLTHASAALSRAIAAILEGDVEKAVRAAQEAVKAAKEAGDNDMLRAVAIAALRIAKEAEKQGNVEVAVKAARVAVEAAKQAGDNDVLRKVAEQALRIAKEAEKQGNVEVAVKAARVAVEAAKQAGDQDVLRKVAEQALRIAKEAEKQGNVEVAVEAARVAVEAAKQAGDQDVLRKVSEQAERISKEAKKQGNSEVSEEARKVADEAKKQTGDGSENLYFQGSHHHHHHG |

**Table S6.** Designed sequences of T=1 tetrahedral cages (symmetry: T3).

| ID | Protomer sequence |
|---|---|
| Tet$_{T=1}$-1 | MGSPELFLQDLRSLVEAARILARLARQRGDEHALERAARWAEQAARQAERLARQARKEGNLELALKALQILVNAAYVLAEIARDRGNEELLEYAARLAEEAARQAIEIWATAMVEGNQQLRTKAAHIILRAAEVLLEIARDRGNQELLEKAASLVDAVAALQQAAAAILEGDVEKAVRAAQEAVKAAKEAGDNDMLRAVAAAAVRIALEALKQGNHEVALKALRVAEEALKQAGGSGGSHHHHHH |
| Tet$_{T=1}$-2 | MGSPELFLQDLRSLVEAARILARLARQRGDEHALERAARWAEQAARQAERLARQARKEGNLELALKALQILVNAAYVLAEIARDRGNEELLEYAARLAEEAARQAIEIWADAMVEGNQQLRTKAAHIILRAAEVLLEIARDRGNQELLEKAASLVDAVAALQQAAAAILEGDVEKAVRAAQEAVKAAKEAGDNDMLRAVAAAATRIALEAFRQGNDEVALKALRVAQEALKQAGGSGGSHHHHHH |
| Tet$_{T=1}$-3 | MGSPELFLQDLRSLVEAARILARLARQRGDEHALERAARWAEQAARQAERLARQARKEGNLELALKALQILVNAAYVLAEIARDRGNEELLEYAARLAEEAARQAIEIWATAMVEGNQQLRTKAAHIILRAAEVLLEIARDRGNQELLEKAASLVDAVAALQQAAAAILEGDVEKAVRAAQEAVKAAKEAGDNDMLRAVAAAAIRIAAEAIKQGNFEVALKALRVAIEALKQAGGSGGSHHHHHH |
| Tet$_{T=1}$-4 | MGSPELFLQDLRSLVEAARILARLARQRGDEHALERAARWAEQAARQAERLARQARKEGNLELALKALQILVNAAYVLAEIARDRGNEELLEYAARLAEEAARQAIEIWANAMIEGNQQLRTKAAHIILRAAEVLLEIARDRGNQELLEKAASLVDAVAALQQAAAAILEGDVEKAVRAAQEAVKAAKEAGDNDMLNAVASAAARIALEAMQQGNLEVALKALRVAEEALKQAGGSGGSHHHHHH |
| Tet$_{T=1}$-5 | MGSPELFLQDLRSLVEAARILARLARQRGDEHALERAARWAEQAARQAERLARQARKEGNLELALKALQILVNAAYVLAEIARDRGNEELLEYAARLAEEAARQAIEIWATAMVEGNQQLRTKAAHIILRAAEVLLEIARDRGNQELLEKAASLVDAVAALQQAAAAILEGDVEKAVRAAQEAVKAAKEAGDNDMLRAVASAAARIALEALAQGNIEVALKALRVAEEALKQAGGSGGSHHHHHH |
| Tet$_{T=1}$-6 | MGSPELFLQDLRSLVEAARILARLARQRGDEHALERAARWAEQAARQAERLARQARKEGNLELALKALQILVNAAYVLAEIARDRGNEELLEYAARLAEEAARQAIEIWATAMEEGNQQLRTKAAHIILRAAEVLLEIARDRGNQELLEKAASLVDAVAALQQAAAAILEGDVEKAVRAAQEAVKAAKEAGDNDMLRAVAAAAIRIALEAMRQGNAEVALKALRVALEALKQAGGSGGSHHHHHH |
| Tet$_{T=1}$-7 | MGSPELFLQDLRSLVEAARILARLARQRGDEHALERAARWAEQAARQAERLARQARKEGNLELALKALQILVNAAYVLAEIARDRGNEELLEYAARLAEEAARQAIEIWADAMVEGNQQLRTKAAHIILRAAEVLLEIARDRGNQELLEKAASLVDAVAALQQAAAAILEGDVEKAVRAAQEAVKAAKEAGDNDMLRAVAAAAARIAIEALRQGNSEVALKALRVAEEALKQAGGSGGSHHHHHH |

**Table S7.** Designed sequences of T=1 octahedral cages (symmetry: O3).

| ID | Protomer sequence |
|---|---|
| Oct$_{T=1}$-1 | MGSPELFLQDLRSLVEAARILARLARQRGDEHALERAARWAEQAARQAERLARQARKEGNLELALKALQILVNAAYVLAEIARDRGNEELLEYAARLAEEAARQAIEIWAQAMEEGNQQLRTKAAHIILRAAEVLLEIARDRGNQELLEKAASLVDAVAALQQAAAAILEGDVEKAVRAAQEAVKAAKEAGDNDMLRAVAIAALRIAKEAEKQGNVEVAVKAARVAVEAAKQAGDQDVLAKVAIQAARIMTEALKQGNLEVALEAAKVASEALKQTGGSGGSHHHHHH |
| Oct$_{T=1}$-2 | MGSPELFLQDLRSLVEAARILARLARQRGDEHALERAARWAEQAARQAERLARQARKEGNLELALKALQILVNAAYVLAEIARDRGNEELLEYAARLAEEAARQAIEIWAQAMEEGNQQLRTKAAHIILRAAEVLLEIARDRGNQELLEKAASLVDAVAALQQAAAAILEGDVEKAVRAAQEAVKAAKEAGDNDMLRAVAIAALRIAKEAEKQGNVEVAVKAARVAVEAAKQAGDQDVLAKVADQAYRIAHEALKQGNLEVALEASKVAMEALKQTGGSGGSHHHHHH |

| | |
|---|---|
| Oct$_{T=1}$-3 | MGSPELFLQDLRSLVEAARILARLARQRGDEHALERAARWAEQAARQAERLARQARKEGNL<br>ELALKALQILVNAAYVLAEIARDRGNEELLEYAARLAEEAARQAIEIWAQAMEEGNQQLRTKAA<br>HIILRAAEVLLEIARDRGNQELLEKAASLVDAVAALQQAAAAILEGDVEKAVRAAQEAVKAAKE<br>AGDNDMLRAVAIAALRIAKEAEKQGNVEVAVKAARVAVEAAKQAGDQDVLKKVAAQAFRIAL<br>EALKQGNVEVALEALKVADEALKQTGGSGGSHHHHHH |
| Oct$_{T=1}$-4 | MGSPELFLQDLRSLVEAARILARLARQRGDEHALERAARWAEQAARQAERLARQARKEGNL<br>ELALKALQILVNAAYVLAEIARDRGNEELLEYAARLAEEAARQAIEIWAQAMEEGNQQLRTKAA<br>HIILRAAEVLLEIARDRGNQELLEKAASLVDAVAALQQAAAAILEGDVEKAVRAAQEAVKAAKE<br>AGDNDMLRAVAIAALRIAKEAEKQGNVEVAVKAARVAVEAAKQAGDQDVLEKVATQALRIAK<br>EANKQGNVEVALEAIKVAQEALKQTGGSGGSHHHHHH |
| Oct$_{T=1}$-5 | MGSPELFLQDLRSLVEAARILARLARQRGDEHALERAARWAEQAARQAERLARQARKEGNL<br>ELALKALQILVNAAYVLAEIARDRGNEELLEYAARLAEEAARQAIEIWAQAMEEGNQQLRTKAA<br>HIILRAAEVLLEIARDRGNQELLEKAASLVDAVAALQQAAAAILEGDVEKAVRAAQEAVKAAKE<br>AGDNDMLRAVAIAALRIAKEAEKQGNVEVAVKAARVAVEAAKQAGDQDVLAKVAEQALRIAIE<br>AIKQGNTEVAAEAIKVADEALKQTGGSGGSHHHHHH |
| Oct$_{T=1}$-6 | MGSPELFLQDLRSLVEAARILARLARQRGDEHALERAARWAEQAARQAERLARQARKEGNL<br>ELALKALQILVNAAYVLAEIARDRGNEELLEYAARLAEEAARQAIEIWAQAMEEGNQQLRTKAA<br>HIILRAAEVLLEIARDRGNQELLEKAASLVDAVAALQQAAAAILEGDVEKAVRAAQEAVKAAKE<br>AGDNDMLRAVAIAALRIAKEAEKQGNVEVAVKAARVAVEAAKQAGDQDVLTKVANQALRIAR<br>EAIKQGNVEVAAEAIKVALEALKQTGGSGGSHHHHHH |
| Oct$_{T=1}$-7 | MGSPELFLQDLRSLVEAARILARLARQRGDEHALERAARWAEQAARQAERLARQARKEGNL<br>ELALKALQILVNAAYVLAEIARDRGNEELLEYAARLAEEAARQAIEIWAQAMEEGNQQLRTKAA<br>HIILRAAEVLLEIARDRGNQELLEKAASLVDAVAALQQAAAAILEGDVEKAVRAAQEAVKAAKE<br>AGDNDMLRAVAIAALRIAKEAEKQGNVEVAVKAARVAVEAAKQAGDQDVLRKVAEQAMRIAR<br>EALKQGNTEVALEAVKVAREALKQTGGSGGSHHHHHH |
| Oct$_{T=1}$-8 | MGSPELFLQDLRSLVEAARILARLARQRGDEHALERAARWAEQAARQAERLARQARKEGNL<br>ELALKALQILVNAAYVLAEIARDRGNEELLEYAARLAEEAARQAIEIWAQAMEEGNQQLRTKAA<br>HIILRAAEVLLEIARDRGNQELLEKAASLVDAVAALQQAAAAILEGDVEKAVRAAQEAVKAAKE<br>AGDNDMLRAVAIAALRIAKEAEKQGNVEVAVKAARVAVEAAKQAGDQDVLVKVAKQAARIML<br>EAIKQGNTEVALEAVKVAAEALKQTGGSGGSHHHHHH |

**Table S8.** Designed sequences of T=1 icosahedral cages (symmetry: I3).

| ID | Protomer sequence |
|---|---|
| Ico$_{T=1}$-1 | GSPELFLQDLRSLVEAARILARLARQRGDEHALERAARWAEQAARQAERLARQARKEGNLEL<br>ALKALQILVNAAYVLAEIARDRGNEELLEYAARLAEEAARQAIEIWAQAMEEGNQQLRTKAAHII<br>LRAAEVLLEIARDRGNQELLEKAASLVDAVAALQQAAAAILEGDVEKAVRAAQEAVKAAKEAG<br>DNDMLRAVAIAALRIAKEAEKQGNVEVAVKAARVAVEAAKQAGDNDVLRKVAEQALRIAKEAE<br>KQGNVEVAVKAARVAVEAAKQAGDNDVLRKVARQAFQIAGKALEQGDIGVAKKAFDVAVEAA<br>SQGGGSGGSHHHHHH |
| Ico$_{T=1}$-2 | GSPELFLQDLRSLVEAARILARLARQRGDEHALERAARWAEQAARQAERLARQARKEGNLEL<br>ALKALQILVNAAYVLAEIARDRGNEELLEYAARLAEEAARQAIEIWAQAMEEGNQQLRTKAAHII<br>LRAAEVLLEIARDRGNQELLEKAASLVDAVAALQQAAAAILEGDVEKAVRAAQEAVKAAKEAG<br>DNDMLRAVAIAALRIAKEAEKQGNVEVAVKAARVAVEAAKQAGDNDVLRKVAEQALRIAKEAE<br>KQGNVEVAVKAARVAVEAAKQAGDNDVLAKVATQALQIAAKAVGQGDTYVAGKAMHVALSAS<br>TQAGGSGGSHHHHHH |
| Ico$_{T=1}$-3 | GSPELFLQDLRSLVEAARILARLARQRGDEHALERAARWAEQAARQAERLARQARKEGNLEL<br>ALKALQILVNAAYVLAEIARDRGNEELLEYAARLAEEAARQAIEIWAQAMEEGNQQLRTKAAHII<br>LRAAEVLLEIARDRGNQELLEKAASLVDAVAALQQAAAAILEGDVEKAVRAAQEAVKAAKEAG<br>DNDMLRAVAIAALRIAKEAEKQGNVEVAVKAARVAVEAAKQAGDNDVLRKVAEQALRIAKEAE<br>KQGNVEVAVKAARVAVEAAKQAGDNDVLEKVANQAMQIALAALQQGDTKVAQKAFDVAERA<br>HSQGGGSGGSHHHHHH |

| | |
|---|---|
| $Ico_{T=1}$-4 | GSPELFLQDLRSLVEAARILARLARQRGDEHALERAARWAEQAARQAERLARQARKEGNLEL<br>ALKALQILVNAAYVLAEIARDRGNEELLEYAARLAEEAARQAIEIWAQAMEEGNQQLRTKAAHII<br>LRAAEVLLEIARDRGNQELLEKAASLVDAVAALQQAAAAILEGDVEKAVRAAQEAVKAAKEAG<br>DNDMLRAVAIAALRIAKEAEKQGNVEVAVKAARVAVEAAKQAGDNDVLRKVAEQALRIAKEAE<br>KQGNVEVAVKAARVAVEAAKQAGDNDVLTKVALQAFEIARKALEQGDQEVAQKALDVAVSAF<br>TQAGGSGGSHHHHHH |

**Table S9.** Designed sequences of C3 crowns. Chain C: hetBGL0-18-17_A32_chC.

| ID | Chain | Protomer sequence |
|---|---|---|
| $Crown_{C3}$-1 | A | MGSLELALKALQILVNAAYVLAEIARDRGNEELLEKAARLAEEAARQAERIARQARKE<br>GNLELALKALQILVNAAYVLAEIARDRGNEELLEYAARLAEEAARQAIEIWAQAMIEGN<br>QQLRTKAAHIILRAAEVLLEIARDRGNQELLEKAASLVDAVAALQQAAAAILEGDVEKA<br>VRAAQEAVKAAKEAGDNDMLLAVAAAAARIALEALLQGNLEVALKALRVALEALKQA<br>GGSGGSHHHHHH |
| | B | MGSPRLVLRALENMVRAAHTLAEIARDNGNEEWLERAARLAEEVARRAERLAREAR<br>KEGNLELALKALQILVNAAYVLAEIARDRGNEEELEYAARLAEEAARQAIEIAADAMIE<br>GNLELALKALQIIVNAAYVLAEIARDRGNEELLEKAASLAEAAAALAEAIAAILEGDVEK<br>AVRAAQEAVKAAKEAGDNDMLRAVAAAAARIALEALAQGNAEVAAKALRVALEALKQ<br>AGGSGGSHHHHHH |
| $Crown_{C3}$-2 | A | MGSLELALKALQILVNAAYVLAEIARDRGNEELLEKAARLAEEAARQAERIARQARKE<br>GNLELALKALQILVNAAYVLAEIARDRGNEELLEYAARLAEEAARQAIEIWARAMVEG<br>NQQLRTKAAHIILRAAEVLLEIARDRGNQELLEKAASLVDAVAALQQAAAAILEGDVEK<br>AVRAAQEAVKAAKEAGDNDMLRAVAAAAARIALEALAQGNYEVAVKALRVALEALKQ<br>AGGSGGSHHHHHH |
| | B | MGSPRLVLRALENMVRAAHTLAEIARDNGNEEWLERAARLAEEVARRAERLAREAR<br>KEGNLELALKALQILVNAAYVLAEIARDRGNEEELEYAARLAEEAARQAIEIAATAMVE<br>GNLELALKALQIIVNAAYVLAEIARDRGNEELLEKAASLAEAAAALAEAIAAILEGDVEK<br>AVRAAQEAVKAAKEAGDNDMLKAVAAAAARIALEAFRQGNYEVALKALRVALEALKQ<br>AGGSGGSHHHHHH |
| $Crown_{C3}$-3 | A | MGSLELALKALQILVNAAYVLAEIARDRGNEELLEKAARLAEEAARQAERIARQARKE<br>GNLELALKALQILVNAAYVLAEIARDRGNEELLEYAARLAEEAARQAIEIWAQAMVEG<br>NQQLRTKAAHIILRAAEVLLEIARDRGNQELLEKAASLVDAVAALQQAAAAILEGDVEK<br>AVRAAQEAVKAAKEAGDNDMLIAVAAAAARIAVEAMRQGNVEVALKALRVASEALKQ<br>AGGSGGSHHHHHH |
| | B | MGSPRLVLRALENMVRAAHTLAEIARDNGNEEWLERAARLAEEVARRAERLAREAR<br>KEGNLELALKALQILVNAAYVLAEIARDRGNEEELEYAARLAEEAARQAIEIAATAMVE<br>GNLELALKALQIIVNAAYVLAEIARDRGNEELLEKAASLAEAAAALAEAIAAILEGDVEK<br>AVRAAQEAVKAAKEAGDNDMLTAVAAAAARIAVEALLQGNTEVALKALRVASEALKQ<br>AGGSGGSHHHHHH |
| $Crown_{C3}$-4 | A | MGSLELALKALQILVNAAYVLAEIARDRGNEELLEKAARLAEEAARQAERIARQARKE<br>GNLELALKALQILVNAAYVLAEIARDRGNEELLEYAARLAEEAARQAIEIWAQAMVEG<br>NQQLRTKAAHIILRAAEVLLEIARDRGNQELLEKAASLVDAVAALQQAAAAILEGDVEK<br>AVRAAQEAVKAAKEAGDNDMLRAVATAAARIAVEALAQGNTEVALKALRVAQEALK<br>QAGGSGGSHHHHHH |
| | B | MGSPRLVLRALENMVRAAHTLAEIARDNGNEEWLERAARLAEEVARRAERLAREAR<br>KEGNLELALKALQILVNAAYVLAEIARDRGNEEELEYAARLAEEAARQAIEIAAEAMVE<br>GNLELALKALQIIVNAAYVLAEIARDRGNEELLEKAASLAEAAAALAEAIAAILEGDVEK<br>AVRAAQEAVKAAKEAGDNDMLRAVAAAAARIAVEAGLQGNFEVALKALRVAQEALK<br>QAGGSGGSHHHHHH |
| $Crown_{C3}$-5 | A | MGSLELALKALQILVNAAYVLAEIARDRGNEELLEKAARLAEEAARQAERIARQARKE<br>GNLELALKALQILVNAAYVLAEIARDRGNEELLEYAARLAEEAARQAIEIWAVAMVEG |

| | | |
|---|---|---|
| | | NQQLRTKAAHIILRAAEVLLEIARDRGNQELLEKAASLVDAVAALQQAAAAILEGDVEK<br>AVRAAQEAVKAAKEAGDNDMLKAVAAAAHRIALEALRQGNTEVALKALRVAREAVK<br>QAGGSGGSHHHHHH |
| | B | MGSPRLVLRALENMVRAAHTLAEIARDNGNEEWLERAARLAEEVARRAERLAREAR<br>KEGNLELALKALQILVNAAYVLAEIARDRGNEEELEYAARLAEEAARQAIEIAALAMVE<br>GNLELALKALQIIVNAAYVLAEIARDRGNEELLEKAASLAEAAAALAEAIAAILEGDVEK<br>AVRAAQEAVKAAKEAGDNDMLFAVANAATRIALEAFRQGNAEVALKALRVADEALK<br>QAGGSGGSHHHHHH |

**Table S10.** Designed sequences of C4 crowns. Chain C: hetBGL0-18-17_A32_chC.

| ID | Chain | Protomer sequence |
|---|---|---|
| Crown$_{C4}$-1 | A | MGSLELALKALQILVNAAYVLAEIARDRGNEELLEKAARLAEEAARQAERIARQARKE<br>GNLELALKALQILVNAAYVLAEIARDRGNEELLEYAARLAEEAARQAIEIWAQAMEEG<br>NQQLRTKAAHIILRAAEVLLEIARDRGNQELLEKAASLVDAVAALQQAAAAILEGDVEK<br>AVRAAQEAVKAAKEAGDNDMLRAVAIAALRIAKEAEKQGNVEVAVKAARVAVEAAK<br>QAGDQDVLEKVARQALRIAEEAIKQGNLEVALEALKVAEEALKQTGGSGGSHHHHH<br>H |
| | B | MGSPRLVLRALENMVRAAHTLAEIARDNGNEEWLERAARLAEEVARRAERLAREAR<br>KEGNLELALKALQILVNAAYVLAEIARDRGNEEELEYAARLAEEAARQAIEIAAQAMEE<br>GNLELALKALQIIVNAAYVLAEIARDRGNEELLEKAASLAEAAAALAEAIAAILEGDVEK<br>AVRAAQEAVKAAKEAGDNDMLRAVAIAALRIAKEAEKQGNVEVAVKAARVAVEAAK<br>QAGDQDVLIKVARQALRIAEEALKQGNTEVAAEALKVAEEALKQTGGSGGSHHHHH<br>H |
| | C | Same with hetBGL0-18-17_A32_chC. |
| Crown$_{C4}$-2 | A | MGSLELALKALQILVNAAYVLAEIARDRGNEELLEKAARLAEEAARQAERIARQARKE<br>GNLELALKALQILVNAAYVLAEIARDRGNEELLEYAARLAEEAARQAIEIWAQAMEEG<br>NQQLRTKAAHIILRAAEVLLEIARDRGNQELLEKAASLVDAVAALQQAAAAILEGDVEK<br>AVRAAQEAVKAAKEAGDNDMLRAVAIAALRIAKEAEKQGNVEVAVKAARVAVEAAK<br>QAGDQDVLLKVATQAARIALEAIKQGNLEVKLEALKVAQEALKQTGGSGGSHHHHH<br>H |
| | B | MGSPRLVLRALENMVRAAHTLAEIARDNGNEEWLERAARLAEEVARRAERLAREAR<br>KEGNLELALKALQILVNAAYVLAEIARDRGNEEELEYAARLAEEAARQAIEIAAQAMEE<br>GNLELALKALQIIVNAAYVLAEIARDRGNEELLEKAASLAEAAAALAEAIAAILEGDVEK<br>AVRAAQEAVKAAKEAGDNDMLRAVAIAALRIAKEAEKQGNVEVAVKAARVAVEAAK<br>QAGDQDVLKKVAAQAARIMLEAIKQGNTEVALEALKVAQEALKQTGGSGGSHHHHH<br>H |
| | C | Same with hetBGL0-18-17_A32_chC. |
| Crown$_{C4}$-3 | A | MGSLELALKALQILVNAAYVLAEIARDRGNEELLEKAARLAEEAARQAERIARQARKE<br>GNLELALKALQILVNAAYVLAEIARDRGNEELLEYAARLAEEAARQAIEIWAQAMEEG<br>NQQLRTKAAHIILRAAEVLLEIARDRGNQELLEKAASLVDAVAALQQAAAAILEGDVEK<br>AVRAAQEAVKAAKEAGDNDMLRAVAIAALRIAKEAEKQGNVEVAVKAARVAVEAAK<br>QAGDQDVLAKVAKQAVRIALEAIKQGNTEVALEAAKVATEALKQTGGSGGSHHHHH<br>H |
| | B | MGSPRLVLRALENMVRAAHTLAEIARDNGNEEWLERAARLAEEVARRAERLAREAR<br>KEGNLELALKALQILVNAAYVLAEIARDRGNEEELEYAARLAEEAARQAIEIAAQAMEE<br>GNLELALKALQIIVNAAYVLAEIARDRGNEELLEKAASLAEAAAALAEAIAAILEGDVEK<br>AVRAAQEAVKAAKEAGDNDMLRAVAIAALRIAKEAEKQGNVEVAVKAARVAVEAAK<br>QAGDQDVLKKVAEQAVRIMLEALKQGNTEVALEAAKVATEALKQTGGSGGSHHHHH<br>H |
| | C | Same with hetBGL0-18-17_A32_chC. |

| | | |
|---|---|---|
| Crown$_{C4}$-4 | A | MGSLELALKALQILVNAAYVLAEIARDRGNEELLEKAARLAEEAARQAERIARQARKEGNLELALKALQILVNAAYVLAEIARDRGNEELLEYAARLAEEAARQAIEIWAQAMEEGNQQLRTKAAHIILRAAEVLLEIARDRGNQELLEKAASLVDAVAALQQAAAAILEGDVEKAVRAAQEAVKAAKEAGDNDMLRAVAIAALRIAKEAEKQGNVEVAVKAARVAVEAAKQAGDQDVLQKVALQAIRIALEALKQGNVEVALEALKVASEALKQTGGSGGSHHHHHH |
| | B | MGSPRLVLRALENMVRAAHTLAEIARDNGNEEWLERAARLAEEVARRAERLAREARKEGNLELALKALQILVNAAYVLAEIARDRGNEEELEYAARLAEEAARQAIEIAAQAMEEGNLELALKALQIIVNAAYVLAEIARDRGNEELLEKAASLAEAAAALAEAIAAILEGDVEKAVRAAQEAVKAAKEAGDNDMLRAVAIAALRIAKEAEKQGNVEVAVKAARVAVEAAKQAGDQDVLKKVAEQAIRIALEALKQGNAEVALEAIKVASEAMKQTGGSGGSHHHHHH |
| | C | Same with hetBGL0-18-17_A32_chC. |
| Crown$_{C4}$-5 | A | MGSLELALKALQILVNAAYVLAEIARDRGNEELLEKAARLAEEAARQAERIARQARKEGNLELALKALQILVNAAYVLAEIARDRGNEELLEYAARLAEEAARQAIEIWAQAMEEGNQQLRTKAAHIILRAAEVLLEIARDRGNQELLEKAASLVDAVAALQQAAAAILEGDVEKAVRAAQEAVKAAKEAGDNDMLRAVAIAALRIAKEAEKQGNVEVAVKAARVAVEAAKQAGDQDVLAKVAEQALRIALEAIKQGNLEVADEAMKVAKEALKQTGGSGGSHHHHHH |
| | B | MGSPRLVLRALENMVRAAHTLAEIARDNGNEEWLERAARLAEEVARRAERLAREARKEGNLELALKALQILVNAAYVLAEIARDRGNEEELEYAARLAEEAARQAIEIAAQAMEEGNLELALKALQIIVNAAYVLAEIARDRGNEELLEKAASLAEAAAALAEAIAAILEGDVEKAVRAAQEAVKAAKEAGDNDMLRAVAIAALRIAKEAEKQGNVEVAVKAARVAVEAAKQAGDQDVLIKVAEQAQRIALEALKQGNLEVALEAMKVAEEALKQTGGSGGSHHHHHH |
| | C | Same with hetBGL0-18-17_A32_chC. |
| Crown$_{C4}$-6 | A | MGSLELALKALQILVNAAYVLAEIARDRGNEELLEKAARLAEEAARQAERIARQARKEGNLELALKALQILVNAAYVLAEIARDRGNEELLEYAARLAEEAARQAIEIWAQAMEEGNQQLRTKAAHIILRAAEVLLEIARDRGNQELLEKAASLVDAVAALQQAAAAILEGDVEKAVRAAQEAVKAAKEAGDNDMLRAVAIAALRIAKEAEKQGNVEVAVKAARVAVEAAKQAGDQDVLAKVAKQALRIALEALKQGNTEVALEAVKVALEALKQTGGSGGSHHHHHH |
| | B | MGSPRLVLRALENMVRAAHTLAEIARDNGNEEWLERAARLAEEVARRAERLAREARKEGNLELALKALQILVNAAYVLAEIARDRGNEEELEYAARLAEEAARQAIEIAAQAMEEGNLELALKALQIIVNAAYVLAEIARDRGNEELLEKAASLAEAAAALAEAIAAILEGDVEKAVRAAQEAVKAAKEAGDNDMLRAVAIAALRIAKEAEKQGNVEVAVKAARVAVEAAKQAGDQDVLRKVSEQAARIALEALKQGNMEVAREAVKVALEALKQTGGSGGSHHHHHH |
| | C | Same with hetBGL0-18-17_A32_chC. |
| Crown$_{C4}$-7 | A | MGSLELALKALQILVNAAYVLAEIARDRGNEELLEKAARLAEEAARQAERIARQARKEGNLELALKALQILVNAAYVLAEIARDRGNEELLEYAARLAEEAARQAIEIWAQAMEEGNQQLRTKAAHIILRAAEVLLEIARDRGNQELLEKAASLVDAVAALQQAAAAILEGDVEKAVRAAQEAVKAAKEAGDNDMLRAVAIAALRIAKEAEKQGNVEVAVKAARVAVEAAKQAGDQDVLKKVALQAARIAKEALKQGNNEVAFEALKVALEALKQTGGSGGSHHHHHH |
| | B | MGSPRLVLRALENMVRAAHTLAEIARDNGNEEWLERAARLAEEVARRAERLAREARKEGNLELALKALQILVNAAYVLAEIARDRGNEEELEYAARLAEEAARQAIEIAAQAMEEGNLELALKALQIIVNAAYVLAEIARDRGNEELLEKAASLAEAAAALAEAIAAILEGDVEKAVRAAQEAVKAAKEAGDNDMLRAVAIAALRIAKEAEKQGNVEVAVKAARVAVEAAKQAGDQDVLAKVAIQAARIMEEALKQGNWEVAGEAAKVALEALKQTGGSGGSHHHHHH |

| | | |
|---|---|---|
| | C | Same with hetBGL0-18-17_A32_chC. |

**Table S11.** Designed sequences of C5 crowns. Chain C: hetBGL0-18-17_A32_chC.

| ID | Chain | Protomer sequence |
|---|---|---|
| Crown$_{C5}$-1 | A | MGSLELALKALQILVNAAYVLAEIARDRGNEELLEKAARLAEEAARQAERIARQARKE<br>GNLELALKALQILVNAAYVLAEIARDRGNEELLEYAARLAEEAARQAIEIWAQAMEEG<br>NQQLRTKAAHIILRAAEVLLEIARDRGNQELLEKAASLVDAVAALQQAAAAILEGDVEK<br>AVRAAQEAVKAAKEAGDNDMLRAVAIAALRIAKEAEKQGNVEVAVKAARVAVEAAK<br>QAGDNDVLRKVAEQALRIAKEAEKQGNVEVAVKAARVAVEAAKQAGDNDVLRKVAD<br>QALEIAKAALEQGDIDVAQKAMDVAVEALTQAGGSGGSHHHHHH |
| | B | MGSPRLVLRALENMVRAAHTLAEIARDNGNEEWLERAARLAEEVARRAERLAREAR<br>KEGNLELALKALQILVNAAYVLAEIARDRGNEEELEYAARLAEEAARQAIEIAAQAMEE<br>GNLELALKALQIIVNAAYVLAEIARDRGNEELLEKAASLAEAAAALAEAIAAILEGDVEK<br>AVRAAQEAVKAAKEAGDNDMLRAVAIAALRIAKEAEKQGNVEVAVKAARVAVEAAK<br>QAGDNDVLRKVAEQALRIAKEAEKQGNVEVAVKAARVAVEAAKQAGDNDVLRKVAE<br>QALEIAKKAAEQGDVGVMQKAMDVALRAAGQAGGSGGSHHHHHH |
| | C | Same with hetBGL0-18-17_A32_chC. |
| Crown$_{C5}$-2 | A | MGSLELALKALQILVNAAYVLAEIARDRGNEELLEKAARLAEEAARQAERIARQARKE<br>GNLELALKALQILVNAAYVLAEIARDRGNEELLEYAARLAEEAARQAIEIWAQAMEEG<br>NQQLRTKAAHIILRAAEVLLEIARDRGNQELLEKAASLVDAVAALQQAAAAILEGDVEK<br>AVRAAQEAVKAAKEAGDNDMLRAVAIAALRIAKEAEKQGNVEVAVKAARVAVEAAK<br>QAGDNDVLRKVAEQALRIAKEAEKQGNVEVAVKAARVAVEAAKQAGDNDVLKKVAK<br>QAMQIAEAALKQGDVDVAWKAMDVAFEALSQAGGSGGSHHHHHH |
| | B | MGSPRLVLRALENMVRAAHTLAEIARDNGNEEWLERAARLAEEVARRAERLAREAR<br>KEGNLELALKALQILVNAAYVLAEIARDRGNEEELEYAARLAEEAARQAIEIAAQAMEE<br>GNLELALKALQIIVNAAYVLAEIARDRGNEELLEKAASLAEAAAALAEAIAAILEGDVEK<br>AVRAAQEAVKAAKEAGDNDMLRAVAIAALRIAKEAEKQGNVEVAVKAARVAVEAAK<br>QAGDNDVLRKVAEQALRIAKEAEKQGNVEVAVKAARVAVEAAKQAGDNDVLNKVAD<br>QALQIAEKALDQGDIGVAWKAMDVALYAATQGGGSGGSHHHHHH |
| | C | Same with hetBGL0-18-17_A32_chC. |
| Crown$_{C5}$-3 | A | MGSLELALKALQILVNAAYVLAEIARDRGNEELLEKAARLAEEAARQAERIARQARKE<br>GNLELALKALQILVNAAYVLAEIARDRGNEELLEYAARLAEEAARQAIEIWAQAMEEG<br>NQQLRTKAAHIILRAAEVLLEIARDRGNQELLEKAASLVDAVAALQQAAAAILEGDVEK<br>AVRAAQEAVKAAKEAGDNDMLRAVAIAALRIAKEAEKQGNVEVAVKAARVAVEAAK<br>QAGDNDVLRKVAEQALRIAKEAEKQGNVEVAVKAARVAVEAAKQAGDNDVLGKVAK<br>QAINIATAALNQGDIKVAGKAFSVAQEALEQGGGSGGSHHHHHH |
| | B | MGSPRLVLRALENMVRAAHTLAEIARDNGNEEWLERAARLAEEVARRAERLAREAR<br>KEGNLELALKALQILVNAAYVLAEIARDRGNEEELEYAARLAEEAARQAIEIAAQAMEE<br>GNLELALKALQIIVNAAYVLAEIARDRGNEELLEKAASLAEAAAALAEAIAAILEGDVEK<br>AVRAAQEAVKAAKEAGDNDMLRAVAIAALRIAKEAEKQGNVEVAVKAARVAVEAAK<br>QAGDNDVLRKVAEQALRIAKEAEKQGNVEVAVKAARVAVEAAKQAGDNDVLRKVAR<br>QAINIATKAVEQGNVKVADKAFSVAESAAQQAGGSGGSHHHHHH |
| | C | Same with hetBGL0-18-17_A32_chC. |
| Crown$_{C5}$-4 | A | MGSLELALKALQILVNAAYVLAEIARDRGNEELLEKAARLAEEAARQAERIARQARKE<br>GNLELALKALQILVNAAYVLAEIARDRGNEELLEYAARLAEEAARQAIEIWAQAMEEG<br>NQQLRTKAAHIILRAAEVLLEIARDRGNQELLEKAASLVDAVAALQQAAAAILEGDVEK<br>AVRAAQEAVKAAKEAGDNDMLRAVAIAALRIAKEAEKQGNVEVAVKAARVAVEAAK<br>QAGDNDVLRKVAEQALRIAKEAEKQGNVEVAVKAARVAVEAAKQAGDNDVLTKVAA<br>QAMEIAKAALKQGDVAVARKAFDVADHALAQGGGSGGSHHHHHH |

| | | |
|---|---|---|
| | B | MGSPRLVLRALENMVRAAHTLAEIARDNGNEEWLERAARLAEEVARRAERLAREAR<br>KEGNLELALKALQILVNAAYVLAEIARDRGNEEELEYAARLAEEAARQAIEIAAQAMEE<br>GNLELALKALQIIVNAAYVLAEIARDRGNEELLEKAASLAEAAAALAEAIAAILEGDVEK<br>AVRAAQEAVKAAKEAGDNDMLRAVAIAALRIAKEAEKQGNVEVAVKAARVAVEAAK<br>QAGDNDVLRKVAEQALRIAKEAEKQGNVEVAVKAARVAVEAAKQAGDNDVLRKVAE<br>QAMEIAKKAFAQGNIEVARKAFDVAVHALTQAGGSGGSHHHHHH |
| | C | Same with hetBGL0-18-17_A32_chC. |
| Crown$_{C5}$-5 | A | MGSLELALKALQILVNAAYVLAEIARDRGNEELLEKAARLAEEAARQAERIARQARKE<br>GNLELALKALQILVNAAYVLAEIARDRGNEELLEYAARLAEEAARQAIEIWAQAMEEG<br>NQQLRTKAAHIILRAAEVLLEIARDRGNQELLEKAASLVDAVAALQQAAAAILEGDVEK<br>AVRAAQEAVKAAKEAGDNDMLRAVAIAALRIAKEAEKQGNVEVAVKAARVAVEAAK<br>QAGDNDVLRKVAEQALRIAKEAEKQGNVEVAVKAARVAVEAAKQAGDNDVLKKVAQ<br>QALDIAAAALRQGDVNVMRKAMDVALQAQTQAGGSGGSHHHHHH |
| | B | MGSPRLVLRALENMVRAAHTLAEIARDNGNEEWLERAARLAEEVARRAERLAREAR<br>KEGNLELALKALQILVNAAYVLAEIARDRGNEEELEYAARLAEEAARQAIEIAAQAMEE<br>GNLELALKALQIIVNAAYVLAEIARDRGNEELLEKAASLAEAAAALAEAIAAILEGDVEK<br>AVRAAQEAVKAAKEAGDNDMLRAVAIAALRIAKEAEKQGNVEVAVKAARVAVEAAK<br>QAGDNDVLRKVAEQALRIAKEAEKQGNVEVAVKAARVAVEAAKQAGDNDVLWKVA<br>GQALDIAARALQQGNLDVARKAMDVAIRAAGQGGGSGGSHHHHHH |
| | C | Same with hetBGL0-18-17_A32_chC. |
| Crown$_{C5}$-6 | A | MGSLELALKALQILVNAAYVLAEIARDRGNEELLEKAARLAEEAARQAERIARQARKE<br>GNLELALKALQILVNAAYVLAEIARDRGNEELLEYAARLAEEAARQAIEIWAQAMEEG<br>NQQLRTKAAHIILRAAEVLLEIARDRGNQELLEKAASLVDAVAALQQAAAAILEGDVEK<br>AVRAAQEAVKAAKEAGDNDMLRAVAIAALRIAKEAEKQGNVEVAVKAARVAVEAAK<br>QAGDNDVLRKVAEQALRIAKEAEKQGNVEVAVKAARVAVEAAKQAGDNDVLKKVFD<br>QATAIAVAAMEQGDADVAKKAFDVANHAWSQAGGSGGSHHHHHH |
| | B | MGSPRLVLRALENMVRAAHTLAEIARDNGNEEWLERAARLAEEVARRAERLAREAR<br>KEGNLELALKALQILVNAAYVLAEIARDRGNEEELEYAARLAEEAARQAIEIAAQAMEE<br>GNLELALKALQIIVNAAYVLAEIARDRGNEELLEKAASLAEAAAALAEAIAAILEGDVEK<br>AVRAAQEAVKAAKEAGDNDMLRAVAIAALRIAKEAEKQGNVEVAVKAARVAVEAAK<br>QAGDNDVLRKVAEQALRIAKEAEKQGNVEVAVKAARVAVEAAKQAGDNDVLLKVAR<br>QATAIAVAAGQQGNLDVMKKAFDVAFEALTQAGGSGGSHHHHHH |
| | C | Same with hetBGL0-18-17_A32_chC. |
| Crown$_{C5}$-7 | A | MGSLELALKALQILVNAAYVLAEIARDRGNEELLEKAARLAEEAARQAERIARQARKE<br>GNLELALKALQILVNAAYVLAEIARDRGNEELLEYAARLAEEAARQAIEIWAQAMEEG<br>NQQLRTKAAHIILRAAEVLLEIARDRGNQELLEKAASLVDAVAALQQAAAAILEGDVEK<br>AVRAAQEAVKAAKEAGDNDMLRAVAIAALRIAKEAEKQGNVEVAVKAARVAVEAAK<br>QAGDNDVLRKVAEQALRIAKEAEKQGNVEVAVKAARVAVEAAKQAGDNDVLRKVAH<br>QALRIADKALDQGDIAVAEKAYEVARRALTQAGGSGGSHHHHHH |
| | B | MGSPRLVLRALENMVRAAHTLAEIARDNGNEEWLERAARLAEEVARRAERLAREAR<br>KEGNLELALKALQILVNAAYVLAEIARDRGNEEELEYAARLAEEAARQAIEIAAQAMEE<br>GNLELALKALQIIVNAAYVLAEIARDRGNEELLEKAASLAEAAAALAEAIAAILEGDVEK<br>AVRAAQEAVKAAKEAGDNDMLRAVAIAALRIAKEAEKQGNVEVAVKAARVAVEAAK<br>QAGDNDVLRKVAEQALRIAKEAEKQGNVEVAVKAARVAVEAAKQAGDNDVLRKVAH<br>QALRIADYALEQGNVGVAEKAYEVAKAAATQAGGSGGSHHHHHH |
| | C | Same with hetBGL0-18-17_A32_chC. |

**Table S12.** Designed sequences of T=4 tetrahedral cages (Symmetry: T33). Chain A: Crown$_{C3}$-3_chA. Chain B: Crown$_{C3}$-3_chB. Chain 'ho' indicates the protomer that forms homotrimer.

| ID | Chain | Protomer sequence |
|---|---|---|
| Tet$_{T=4}$-1 | A | Same with Crown$_{C3}$-3_chA |
| | B | Same with Crown$_{C3}$-3_chB |
| | C | MGSPELFLQDLRSLVEAARILARLARQRGDEHALERAARWAEQAARQAERLARQARKE<br>GNLELALKALQILVNAAYVLAEIARDRGNEELLEYAARLAEEAARQAIEIAAQAMEEGNFE<br>LALEALEIINEAARVLARIAHHRGNQELLEKAASLTHASAALSRAIAAILEGDVEKAVRAAQ<br>EAVKAAKEAGDNDMLRAVAIAALRIAKEAEKQGNVEVAVKAARVAVEAAKQAGDNDVL<br>RKVAEQALRIAKEAEKQGNTVVAFQALVVAASAATAAGDDDVLRKVLEQFMRIVKELKK<br>QG |
| | ho | MGSPELFLQDLRSLVEAARILARLARQRGDEHALERAARWAEQAARQAERLARQARKE<br>GNLELALKALQILVNAAYVLAEIARDRGNEELLEYAARLAEEAARQAIEIWAQAMEEGNQ<br>QLRTKAAHIILRAAEVLLEIARDRGNQELLEKAASLVDAVAALQQAAAAILEGDVEKAVRA<br>AQEAVKAAKEAGDNDMLRAVAIAALRIAKEAEKQGNVEVAVKAARVAVEAAKQAGDND<br>VLRKVAEQALRIAKEAEKQGNTVVAFEALEVAAEAASAAGDDDVLRKVLEQFKRIVKELE<br>KQGGSGGSHHHHHH |
| Tet$_{T=4}$-2 | A | Same with Crown$_{C3}$-3_chA |
| | B | Same with Crown$_{C3}$-3_chB |
| | C | MGSPELFLQDLRSLVEAARILARLARQRGDEHALERAARWAEQAARQAERLARQARKE<br>GNLELALKALQILVNAAYVLAEIARDRGNEELLEYAARLAEEAARQAIEIAAQAMEEGNFE<br>LALEALEIINEAARVLARIAHHRGNQELLEKAASLTHASAALSRAIAAILEGDVEKAVRAAQ<br>EAVKAAKEAGDNDMLRAVAIAALRIAKEAEKQGNVEVAVKAARVAVEAAKQAGDNDVL<br>RKVAEQALRIAKEAEKQGNVHVALQALAVAGEAASAAGDTDVLRKVLEQFRRIVKELEK<br>QG |
| | ho | MGSPELFLQDLRSLVEAARILARLARQRGDEHALERAARWAEQAARQAERLARQARKE<br>GNLELALKALQILVNAAYVLAEIARDRGNEELLEYAARLAEEAARQAIEIWAQAMEEGNQ<br>QLRTKAAHIILRAAEVLLEIARDRGNQELLEKAASLVDAVAALQQAAAAILEGDVEKAVRA<br>AQEAVKAAKEAGDNDMLRAVAIAALRIAKEAEKQGNVEVAVKAARVAVEAAKQAGDND<br>VLRKVAEQALRIAKEAEKQGNVHVALLALDVAATAAAVAGDTDVLRKVLEQFQRIVKELE<br>KQGGSGGSHHHHHH |
| Tet$_{T=4}$-3 | A | Same with Crown$_{C3}$-3_chA |
| | B | Same with Crown$_{C3}$-3_chB |
| | C | MGSPELFLQDLRSLVEAARILARLARQRGDEHALERAARWAEQAARQAERLARQARKE<br>GNLELALKALQILVNAAYVLAEIARDRGNEELLEYAARLAEEAARQAIEIAAQAMEEGNFE<br>LALEALEIINEAARVLARIAHHRGNQELLEKAASLTHASAALSRAIAAILEGDVEKAVRAAQ<br>EAVKAAKEAGDNDMLRAVAIAALRIAKEAEKQGNVEVAVKAARVAVEAAKQAGDNDVL<br>RKVAEQALRIAKEAEKQGNTRVAMEAAGVALEAAKVAGDGDVIQKVVEQVIRIVDEVEK<br>QG |
| | ho | MGSPELFLQDLRSLVEAARILARLARQRGDEHALERAARWAEQAARQAERLARQARKE<br>GNLELALKALQILVNAAYVLAEIARDRGNEELLEYAARLAEEAARQAIEIWAQAMEEGNQ<br>QLRTKAAHIILRAAEVLLEIARDRGNQELLEKAASLVDAVAALQQAAAAILEGDVEKAVRA<br>AQEAVKAAKEAGDNDMLRAVAIAALRIAKEAEKQGNVEVAVKAARVAVEAAKQAGDND<br>VLRKVAEQALRIAKEAEKQGNTRVAMRAVKVAAEAAKTAGDQDVIQKVLEQVGRIGGEV<br>IKQGGSGGSHHHHHH |
| Tet$_{T=4}$-4 | A | Same with Crown$_{C3}$-3_chA |
| | B | Same with Crown$_{C3}$-3_chB |
| | C | MGSPELFLQDLRSLVEAARILARLARQRGDEHALERAARWAEQAARQAERLARQARKE<br>GNLELALKALQILVNAAYVLAEIARDRGNEELLEYAARLAEEAARQAIEIAAQAMEEGNFE |

| | | |
|---|---|---|
| | | LALEALEIINEAARVLARIAHHRGNQELLEKAASLTHASAALSRAIAAILEGDVEKAVRAAQ<br>EAVKAAKEAGDNDMLRAVAIAALRIAKEAEKQGNVEVAVKAARVAVEAAKQAGDNDVL<br>RKVAEQALRIAKEAEKQGNPVVALPAVGVAWEAADWAGDPQVGRKVDEQFLRIADEIE<br>KQG |
| | ho | MGSPELFLQDLRSLVEAARILARLARQRGDEHALERAARWAEQAARQAERLARQARKE<br>GNLELALKALQILVNAAYVLAEIARDRGNEELLEYAARLAEEAARQAIEIWAQAMEEGNQ<br>QLRTKAAHIILRAAEVLLEIARDRGNQELLEKAASLVDAVAALQQAAAAILEGDVEKAVRA<br>AQEAVKAAKEAGDNDMLRAVAIAALRIAKEAEKQGNVEVAVKAARVAVEAAKQAGDND<br>VLRKVAEQALRIAKEAEKQGNYVVALEAQKVAWEAATVAGDPQVVRKVREQFRRIAAEI<br>QKQGGSGGSHHHHHH |
| Tet$_{T=4}$-5 | A | Same with Crown$_{C3}$-3_chA |
| | B | Same with Crown$_{C3}$-3_chB |
| | C | MGSPELFLQDLRSLVEAARILARLARQRGDEHALERAARWAEQAARQAERLARQARKE<br>GNLELALKALQILVNAAYVLAEIARDRGNEELLEYAARLAEEAARQAIEIAAQAMEEGNFE<br>LALEALEIINEAARVLARIAHHRGNQELLEKAASLTHASAALSRAIAAILEGDVEKAVRAAQ<br>EAVKAAKEAGDNDMLRAVAIAALRIAKEAEKQGNVEVAVKAARVAVEAAKQAGDNDVL<br>RKVAEQALRIAKEAEKQGNPGVAMDAVSVAFEAAKQAGDKGVTDKVTEQFGRIVSEIEK<br>QG |
| | ho | MGSPELFLQDLRSLVEAARILARLARQRGDEHALERAARWAEQAARQAERLARQARKE<br>GNLELALKALQILVNAAYVLAEIARDRGNEELLEYAARLAEEAARQAIEIWAQAMEEGNQ<br>QLRTKAAHIILRAAEVLLEIARDRGNQELLEKAASLVDAVAALQQAAAAILEGDVEKAVRA<br>AQEAVKAAKEAGDNDMLRAVAIAALRIAKEAEKQGNVEVAVKAARVAVEAAKQAGDND<br>VLRKVAEQALRIAKEAEKQGNPGVAMMAIGVAAEAAKEAGDKGVTDKVTEQFVRIVPEI<br>KKQGGSGGSHHHHHH |

**Table S13.** Designed sequences of T=4 octahedral cages (Symmetry: O43). Chain A: Crown$_{C4}$-2_chA. Chain B: Crown$_{C4}$-2_chB. Chain 'ho' indicates the protomer that forms homotrimer.

| ID | Chain | Protomer sequence |
|---|---|---|
| Oct$_{T=4}$-1 | A | Same with Crown$_{C4}$-2_chA |
| | B | Same with Crown$_{C4}$-2_chB |
| | C | MGSPELFLQDLRSLVEAARILARLARQRGDEHALERAARWAEQAARQAERLARQARK<br>EGNLELALKALQILVNAAYVLAEIARDRGNEELLEYAARLAEEAARQAIEIAAQAMEEGN<br>FELALEALEIINEAARVLARIAHHRGNQELLEKAASLTHASAALSRAIAAILEGDVEKAVR<br>AAQEAVKAAKEAGDNDMLRAVAIAALRIAKEAEKQGNVEVAVKAARVAVEAAKQAGDN<br>DVLRKVAEQALRIAKEAEKQGNVGVAFKAIDVAIEAAGQAGDDDVRAKVSDQMLRIIDE<br>TEKQG |
| | ho | MGSPELFLQDLRSLVEAARILARLARQRGDEHALERAARWAEQAARQAERLARQARK<br>EGNLELALKALQILVNAAYVLAEIARDRGNEELLEYAARLAEEAARQAIEIWAQAMEEG<br>NQQLRTKAAHIILRAAEVLLEIARDRGNQELLEKAASLVDAVAALQQAAAAILEGDVEKA<br>VRAAQEAVKAAKEAGDNDMLRAVAIAALRIAKEAEKQGNVEVAVKAARVAVEAAKQAG<br>DNDVLRKVAEQALRIAKEAEKQGNVKVAFKAMRVAAEAAKQAGDLDVRDKVSDQTLRI<br>QDEKEKQGGSGGSHHHHHH |
| Oct$_{T=4}$-2 | A | Same with Crown$_{C4}$-2_chA |
| | B | Same with Crown$_{C4}$-2_chB |
| | C | MGSPELFLQDLRSLVEAARILARLARQRGDEHALERAARWAEQAARQAERLARQARK<br>EGNLELALKALQILVNAAYVLAEIARDRGNEELLEYAARLAEEAARQAIEIAAQAMEEGN<br>FELALEALEIINEAARVLARIAHHRGNQELLEKAASLTHASAALSRAIAAILEGDVEKAVR<br>AAQEAVKAAKEAGDNDMLRAVAIAALRIAKEAEKQGNVEVAVKAARVAVEAAKQAGDN |

| | | |
|---|---|---|
| | | DVLRKVAEQALRIAKEAEKQGNVVVASKAIGVAEEAAYQAGDTDVQKKVSDQNYRIIQE<br>RIKQG |
| | ho | MGSPELFLQDLRSLVEAARILARLARQRGDEHALERAARWAEQAARQAERLARQARK<br>EGNLELALKALQILVNAAYVLAEIARDRGNEELLEYAARLAEEAARQAIEIWAQAMEEG<br>NQQLRTKAAHIILRAAEVLLEIARDRGNQELLEKAASLVDAVAALQQAAAAILEGDVEKA<br>VRAAQEAVKAAKEAGDNDMLRAVAIAALRIAKEAEKQGNVEVAVKAARVAVEAAKQAG<br>DNDVLRKVAEQALRIAKEAEKQGNVSVAAKANLVAIEAAKQAGDTDVQKKVSEQNQRI<br>TSEFEKQGGSGGSHHHHHH |
| Oct$_{T=4}$-3 | A | Same with Crown$_{C4}$-2_chA |
| | B | Same with Crown$_{C4}$-2_chB |
| | C | MGSPELFLQDLRSLVEAARILARLARQRGDEHALERAARWAEQAARQAERLARQARK<br>EGNLELALKALQILVNAAYVLAEIARDRGNEELLEYAARLAEEAARQAIEIAAQAMEEGN<br>FELALEALEIINEAARVLARIAHHRGNQELLEKAASLTHASAALSRAIAAILEGDVEKAVR<br>AAQEAVKAAKEAGDNDMLRAVAIAALRIAKEAEKQGNVEVAVKAARVAVEAAKQAGDN<br>DVLRKVAEQALRIAKEAEKQGNVIVAMKAIDVAVEAAGQAGDIDVIKKVADQTQRIVDE<br>WLKQG |
| | ho | MGSPELFLQDLRSLVEAARILARLARQRGDEHALERAARWAEQAARQAERLARQARK<br>EGNLELALKALQILVNAAYVLAEIARDRGNEELLEYAARLAEEAARQAIEIWAQAMEEG<br>NQQLRTKAAHIILRAAEVLLEIARDRGNQELLEKAASLVDAVAALQQAAAAILEGDVEKA<br>VRAAQEAVKAAKEAGDNDMLRAVAIAALRIAKEAEKQGNVEVAVKAARVAVEAAKQAG<br>DNDVLRKVAEQALRIAKEAEKQGNVYVAAKAVQVAAEAAKQAGDIDVLKKVIGQTQRIR<br>DEWVKQGGSGGSHHHHHH |
| Oct$_{T=4}$-4 | A | Same with Crown$_{C4}$-2_chA |
| | B | Same with Crown$_{C4}$-2_chB |
| | C | MGSPELFLQDLRSLVEAARILARLARQRGDEHALERAARWAEQAARQAERLARQARK<br>EGNLELALKALQILVNAAYVLAEIARDRGNEELLEYAARLAEEAARQAIEIAAQAMEEGN<br>FELALEALEIINEAARVLARIAHHRGNQELLEKAASLTHASAALSRAIAAILEGDVEKAVR<br>AAQEAVKAAKEAGDNDMLRAVAIAALRIAKEAEKQGNVEVAVKAARVAVEAAKQAGDN<br>DVLRKVAEQALRIAKEAEKQGNVSVAFKAIDVAVEAAGQAGDWDVWGKVYKQTWRIIN<br>EAIKQG |
| | ho | MGSPELFLQDLRSLVEAARILARLARQRGDEHALERAARWAEQAARQAERLARQARK<br>EGNLELALKALQILVNAAYVLAEIARDRGNEELLEYAARLAEEAARQAIEIWAQAMEEG<br>NQQLRTKAAHIILRAAEVLLEIARDRGNQELLEKAASLVDAVAALQQAAAAILEGDVEKA<br>VRAAQEAVKAAKEAGDNDMLRAVAIAALRIAKEAEKQGNVEVAVKAARVAVEAAKQAG<br>DNDVLRKVAEQALRIAKEAEKQGNVYVALKALKVAIEAAGQAGDGDVWGKVFVQSKRI<br>EDEWNKQGGSGGSHHHHHH |

**Table S14.** Designed sequences of T=4 icosahedral cages (Symmetry: I43). Chain A: Crown$_{C5}$-1_chA. Chain B: Crown$_{C5}$-1_chB. Chain 'ho' indicates the protomer that forms homotrimer.

| ID | Chain | Protomer sequence |
|---|---|---|
| Ico$_{T=4}$-1 | A | Same with Crown$_{C5}$-1_chA |
| | B | Same with Crown$_{C5}$-1_chB |
| | C | MGSPELFLQDLRSLVEAARILARLARQRGDEHALERAARWAEQAARQAERLARQARKE<br>GNLELALKALQILVNAAYVLAEIARDRGNEELLEYAARLAEEAARQAIEIAAQAMEEGNFE<br>LALEALEIINEAARVLARIAHHRGNQELLEKAASLTHASAALSRAIAAILEGDVEKAVRAAQ<br>EAVKAAKEAGDNDMLRAVAIAALRIAKEAEKQGNVEVAVKAARVAVEAAKQAGDNDVL<br>RKVAEQALRIAKEAEKQGNVEVAVKAARVAVEAAKQAGDNDVLRKVAEQALRIAKEAEK<br>QGNVIVAVEAAVVAVEAATLAGDQDVLRKVLEQLERIRKEAEKQGGSGGSHHHHHH |

| | | |
|---|---|---|
| | ho | MGSPELFLQDLRSLVEAARILARLARQRGDEHALERAARWAEQAARQAERLARQARKE<br>GNLELALKALQILVNAAYVLAEIARDRGNEELLEYAARLAEEAARQAIEIWAQAMEEGNQ<br>QLRTKAAHIILRAAEVLLEIARDRGNQELLEKAASLVDAVAALQQAAAAILEGDVEKAVRA<br>AQEAVKAAKEAGDNDMLRAVAIAALRIAKEAEKQGNVEVAVKAARVAVEAAKQAGDND<br>VLRKVAEQALRIAKEAEKQGNVEVAVKAARVAVEAAKQAGDNDVLRKVAEQALRIAKEA<br>EKQGNVVVATMAAIVAIEAATLAGDQDVLRKVKEQLERIAKEAAKQGGSGGSHHHHHH |
| Ico$_{T=4}$-2 | A | Same with Crown$_{C5}$-1_chA |
| | B | Same with Crown$_{C5}$-1_chB |
| | C | MGSPELFLQDLRSLVEAARILARLARQRGDEHALERAARWAEQAARQAERLARQARKE<br>GNLELALKALQILVNAAYVLAEIARDRGNEELLEYAARLAEEAARQAIEIAAQAMEEGNFE<br>LALEALEIINEAARVLARIAHHRGNQELLEKAASLTHASAALSRAIAAILEGDVEKAVRAAQ<br>EAVKAAKEAGDNDMLRAVAIAALRIAKEAEKQGNVEVAVKAARVAVEAAKQAGDNDVL<br>RKVAEQALRIAKEAEKQGNVEVAVKAARVAVEAAKQAGDNDVLRKVAEQALRIAKEAEK<br>QGNVIVALAAALVAVEAASVAGDQDVMRKVLEQLERILKELEKQGGSGGSHHHHHH |
| | ho | MGSPELFLQDLRSLVEAARILARLARQRGDEHALERAARWAEQAARQAERLARQARKE<br>GNLELALKALQILVNAAYVLAEIARDRGNEELLEYAARLAEEAARQAIEIWAQAMEEGNQ<br>QLRTKAAHIILRAAEVLLEIARDRGNQELLEKAASLVDAVAALQQAAAAILEGDVEKAVRA<br>AQEAVKAAKEAGDNDMLRAVAIAALRIAKEAEKQGNVEVAVKAARVAVEAAKQAGDND<br>VLRKVAEQALRIAKEAEKQGNVEVAVKAARVAVEAAKQAGDNDVLRKVAEQALRIAKEA<br>EKQGNVIVAAGAAAVALEAATVAGDQDVMRKVKEQLERISKEAQKQGGSGGSHHHHH<br>H |
| Ico$_{T=4}$-3 | A | Same with Crown$_{C5}$-1_chA |
| | B | Same with Crown$_{C5}$-1_chB |
| | C | MGSPELFLQDLRSLVEAARILARLARQRGDEHALERAARWAEQAARQAERLARQARKE<br>GNLELALKALQILVNAAYVLAEIARDRGNEELLEYAARLAEEAARQAIEIAAQAMEEGNFE<br>LALEALEIINEAARVLARIAHHRGNQELLEKAASLTHASAALSRAIAAILEGDVEKAVRAAQ<br>EAVKAAKEAGDNDMLRAVAIAALRIAKEAEKQGNVEVAVKAARVAVEAAKQAGDNDVL<br>RKVAEQALRIAKEAEKQGNVEVAVKAARVAVEAAKQAGDNDVLRKVAEQALRIAKEAEK<br>QGNVLVAIAAASVAVEAASLAGDQDVLRKVLEQLERIAKEAEKQGGSGGSHHHHHH |
| | ho | MGSPELFLQDLRSLVEAARILARLARQRGDEHALERAARWAEQAARQAERLARQARKE<br>GNLELALKALQILVNAAYVLAEIARDRGNEELLEYAARLAEEAARQAIEIWAQAMEEGNQ<br>QLRTKAAHIILRAAEVLLEIARDRGNQELLEKAASLVDAVAALQQAAAAILEGDVEKAVRA<br>AQEAVKAAKEAGDNDMLRAVAIAALRIAKEAEKQGNVEVAVKAARVAVEAAKQAGDND<br>VLRKVAEQALRIAKEAEKQGNVEVAVKAARVAVEAAKQAGDNDVLRKVAEQALRIAKEA<br>EKQGNVIVATQAAIVALEAATVAGDQDVLRKVKEQLERILKEARKQGGSGGSHHHHHH |
| Ico$_{T=4}$-4 | A | Same with Crown$_{C5}$-1_chA |
| | B | Same with Crown$_{C5}$-1_chB |
| | C | MGSPELFLQDLRSLVEAARILARLARQRGDEHALERAARWAEQAARQAERLARQARKE<br>GNLELALKALQILVNAAYVLAEIARDRGNEELLEYAARLAEEAARQAIEIAAQAMEEGNFE<br>LALEALEIINEAARVLARIAHHRGNQELLEKAASLTHASAALSRAIAAILEGDVEKAVRAAQ<br>EAVKAAKEAGDNDMLRAVAIAALRIAKEAEKQGNVEVAVKAARVAVEAAKQAGDNDVL<br>RKVAEQALRIAKEAEKQGNVEVAVKAARVAVEAAKQAGDNDVLRKVAEQALRIAKEAEK<br>QGNVIVAVRAALVAAQAATEAGDQDVLRKVDEQLERILKEARKQGGSGGSHHHHHH |
| | ho | MGSPELFLQDLRSLVEAARILARLARQRGDEHALERAARWAEQAARQAERLARQARKE<br>GNLELALKALQILVNAAYVLAEIARDRGNEELLEYAARLAEEAARQAIEIWAQAMEEGNQ<br>QLRTKAAHIILRAAEVLLEIARDRGNQELLEKAASLVDAVAALQQAAAAILEGDVEKAVRA<br>AQEAVKAAKEAGDNDMLRAVAIAALRIAKEAEKQGNVEVAVKAARVAVEAAKQAGDND<br>VLRKVAEQALRIAKEAEKQGNVEVAVKAARVAVEAAKQAGDNDVLRKVAEQALRIAKEA<br>EKQGNVVVATQAAIVAIQAARLAGDQDVGRKVIEQEERIAKEARKQGGSGGSHHHHHH |

| | | |
|---|---|---|
| $Ico_{T=4}$-5 | A | Same with $Crown_{C5}$-1_chA |
| | B | Same with $Crown_{C5}$-1_chB |
| | C | MGSPELFLQDLRSLVEAARILARLARQRGDEHALERAARWAEQAARQAERLARQARKE<br>GNLELALKALQILVNAAYVLAEIARDRGNEELLEYAARLAEEAARQAIEIAAQAMEEGNFE<br>LALEALEIINEAARVLARIAHHRGNQELLEKAASLTHASAALSRAIAAILEGDVEKAVRAAQ<br>EAVKAAKEAGDNDMLRAVAIAALRIAKEAEKQGNVEVAVKAARVAVEAAKQAGDNDVL<br>RKVAEQALRIAKEAEKQGNVEVAVKAARVAVEAAKQAGDNDVLRKVAEQALRIAKEAEK<br>QGNVVVALAAAAVAVRAAREAGDQDVLRKVLEQLERILKEGDKQGGSGGSHHHHHH |
| | ho | MGSPELFLQDLRSLVEAARILARLARQRGDEHALERAARWAEQAARQAERLARQARKE<br>GNLELALKALQILVNAAYVLAEIARDRGNEELLEYAARLAEEAARQAIEIWAQAMEEGNQ<br>QLRTKAAHIILRAAEVLLEIARDRGNQELLEKAASLVDAVAALQQAAAAILEGDVEKAVRA<br>AQEAVKAAKEAGDNDMLRAVAIAALRIAKEAEKQGNVEVAVKAARVAVEAAKQAGDND<br>VLRKVAEQALRIAKEAEKQGNVEVAVKAARVAVEAAKQAGDNDVLRKVAEQALRIAKEA<br>EKQGNVVVATRAAAVALEAALLAGDQDVLRKVEEQLERILKEAEKQGGSGGSHHHHHH |

**Table S15.** CryoEM data collection information of T=4 cages ($Oct_{T=4}$-3 and $Ico_{T=4}$-4).

| | $Oct_{T=4}$-3 (T=4) (EMDB-40260) | $Ico_{T=4}$-4 (T=4) (EMDB-40268) |
|---|---|---|
| **Data collection and processing** | | |
| Magnification | 45,000 | 210,000 |
| Voltage (kV) | 200 | 300 |
| Electron exposure (e–/Å$^2$) | 50 | 61 |
| Defocus range (μm) | Between -0.8 and -1.8 | Between -0.8 and -1.8 |
| Pixel size (Å) | 0.44 | 0.42 |
| Symmetry imposed | Octahedral symmetry | Icosahedral symmetry |
| Initial particle images (no.) | 35,828 + 21,472 | 12,780 |
| Final particle images (no.) | 29,069 | 6,896 |
| Map resolution (Å) | 6.87 | 12.78 |

**Table S15.** CryoEM data collection information of off target small cages (T=1 cages) found in $Oct_{T=4}$-3 and $Ico_{T=4}$-4 systems.

| | $Oct_{T=4}$-3 (T=1) (EMDB-40269) | $Ico_{T=4}$-4 (T=1) (EMDB-40267) |
|---|---|---|
| **Data collection and processing** | | |
| Magnification | 45,000 | 210,000 |
| Voltage (kV) | 200 | 300 |
| Electron exposure (e–/Å$^2$) | 50 | 61 |
| Defocus range (μm) | Between -0.8 and -1.8 | Between -0.8 and -1.8 |
| Pixel size (Å) | 0.44 | 0.42 |
| Symmetry imposed | Octahedral symmetry | Icosahedral symmetry |
| Initial particle images (no.) | 35,828 + 21,472 | 12,780 |
| Final particle images (no.) | 15,852 | 1,172 |
| Map resolution (Å) | 6.34 | 6.54 |
